# Genetic Basis, Quantitative Nature, and Functional Relevance of Evolutionarily Conserved DNA Methylation

**DOI:** 10.1101/2024.11.07.622474

**Authors:** Zheng (Joe) Dong, Samantha Schaffner, Maggie Fu, Joanne Whitehead, Julia L. MacIsaac, David H. Rehkopf, W. Thomas Boyce, Luis Rosero-Bixby, Lluis Quintana-Murci, Etienne Patin, Gregory E. Miller, Keegan Korthauer, Michael S. Kobor

## Abstract

DNA methylation (DNAm) is a key epigenetic mark that modulates regulatory elements and gene expression, playing a crucial role in mammalian development and physiological function. Despite extensive characterization of DNAm profiles across species, little is known about its evolutionary conservation. Here, we conducted a comparative epigenome-wide analysis of great apes to identify and characterize sequence- and methylation-conserved CpGs (MCCs). Using 202 DNAm arrays, alongside 6 matched genotype and 13 matched transcriptomic datasets, we identified 11,500 MCCs for which methylation was evolutionarily related to sequences of CpGs and methylation quantitative trait loci. MCCs were the most stable across healthy human tissues and exhibited weaker genetic associations than other CpGs. Moreover, MCCs showed minimal associations with demographic, environmental factors, and noncancer diseases, yet demonstrated stronger associations with certain cancers than other CpGs, particularly gastrointestinal cancers. Functional enrichment analysis revealed that genes associated with MCC methylation in cancer were enriched for cancer driver genes and canonical cancer pathways, highlighting a significant regulatory role for MCCs in tumorigenesis. Collectively, our findings reveal the extent of DNAm conservation in great ape evolution, its association with genetic conservation, and its relevance to human diseases. These integrative analyses offer evolutionary insights into epigenetic variation and its functional implications in human populations.

---

DNA methylation (DNAm) is an important epigenetic mark that plays a role in genome integrity and gene regulation. In mammals, DNAm occurs primarily at cytosine-phosphate-guanine (CpG) dinucleotides, of which ∼60%–80% are methylated in somatic cells^1^. Gene bodies of highly expressed genes are typically hypermethylated, whereas regulatory regions of genes (e.g., promoters) show a low DNAm level^2^. Moreover, the quantitative characteristics of DNAm, such as methylation level and variability, have been extensively demonstrated to be associated with tissue development, environmental exposures, and disease susceptibility^3,4^. Specifically, aberrant DNAm in malignancies has been widely characterized as global loss and focal gain of DNAm^5,6^. Altered DNAm levels have been associated with a wide range of environmental exposures, including air pollution, nutritional factors, and smoking^7^. In addition, DNAm has been shown to act as a potential mediator of the association between genetics and diseases^8^. Genetic variants associated with quantitative changes in DNAm of CpGs are typically referred to as methylation quantitative trait loci (mQTLs).

Given the role of DNAm in various biological processes, there is an emerging interest in its quantitative measurements across species to gain insight into evolution. Numerous comparative epigenomics investigations initially elucidated DNAm changes between primates in single tissue samples or in samples with multiple tissues in a tissue-specific manner^9–13^. Species-specific DNAm may be a response to environmental influences^14^ and associated with differences in underlying genomic sequences (i.e., CpG and its neighboring nucleotides)^15–19^. For example, the frequency of 3-mer DNA sequences, such as ACG, CGT, CGA, and TCG, has been reported to account for more than 80% of the observed DNAm variance across vertebrate evolution^17^. By integration with transcriptional data, species-specific DNAm has been suggested to be associated with transcriptional modification of genes involved in species-specific adaptations and phenotypic variability, particularly human-specific traits^9–13,20^. These studies have therefore improved our understanding of the evolutionary context of epigenetic roles in diseases and complex traits.

However, previous studies of DNAm in evolution focused mostly on species-specific DNAm signals, leaving evolutionarily conserved DNAm largely uncharacterized. For example, the genetic basis of methylation-conserved CpGs (MCCs) remains to be fully explored. Although DNAm levels in certain genomic regions have been reported to be evolutionarily conserved based on the positive relation between DNAm conservation and underlying sequence conservation^15,18,19,21–27^, it is still unclear whether—and to what extent—DNAm conservation is associated with the conservation of genetic variation outside of CpG and neighboring nucleotides, such as mQTLs. Moreover, as evolutionary conservation in DNA sequences is a commonly used indicator of functional importance^28^, the presence of conserved DNAm prompted interest in elucidating its potential functional relevance. However, there have been no comprehensive examinations of DNAm variability of MCCs in human populations or their associations with lifestyle and health factors.

Using more than 200 in-house and publicly available data sets containing a total of 32,060 DNAm array samples, 6 matched genotype data sets containing a total of 1058 samples, and 13 matched transcription data sets containing a total of 3860 samples, this study comprehensively characterized MCCs with regard to their genetic basis, quantitative nature, and potential functional relevance. Specifically, we identified 1) which CpGs genome-wide show conserved DNAm levels across great ape species, 2) whether mQTL sequence conservation is associated with DNAm conservation, 3) the characteristics of DNAm levels and variability of MCCs in human populations, 4) the lifestyle and health factors associated with MCCs, 5) functional enrichment of genes whose expression is associated with MCC methylation in cancer driver genes and biological pathways, and then 6) developed an integrated tool to illustrate evolutionary conservation of CpGs genome-wide. DNAm conservation was shown to be positively associated with genomic sequence conservation not only in CpGs but also in mQTLs. In this comprehensive investigation, MCCs were characterized by their extensive DNAm stability in a wide range of healthy human tissues, DNAm dynamics in multiple cancers, and their regulatory roles in cancer driver genes. Our analyses provided a better understanding of epigenomic conservation in terms of genetic associations, quantitative nature, and potential functional relevance in human populations, as well as a basis for using DNAm conservation information in further epigenetics investigations.

## Results

### Discovery of evolutionary conservation DNAm at single-base resolution

#### Genome-wide identification of 11,500 MCCs across great apes

The workflow for identification of MCCs (CpG sites conserved in both sequence and DNAm) across great apes is illustrated in Figure 1a. Briefly, we retrieved 32 Illumina HumanMethylation450 BeadChip (HM450K) samples from the NCBI Gene Expression Omnibus (GEO) database under accession number GSE41782, which consisted of whole-blood samples from nine human subjects (*Homo sapiens*), five chimpanzees (*Pan troglodytes*), six bonobos (*Pan paniscus*), six gorillas (*Gorilla gorilla gorilla* and *Gorilla beringei graueri*), and six orangutans (*Pongo pygmaeus abelii*) (data sources are described in Supplementary Table S1)^29^. The sequences of HM450K array probes designed based on the human reference genome, hg19, were aligned to reference genomes for five species of great ape (i.e., panTro5 for chimpanzee, panPan2 for bonobo, gorGor4 for gorilla, ponAbe3 for orangutan, and hg19 for human). The 169,500 probes with unique alignment targets (i.e., unambiguous or non–cross-reactive) and high sequence conservation (including ≥ 90% identity and perfect match at the CpG locus^30–33^) across all five species of great apes were retained (see Methods section “Identifying sequence conservation of CpGs”). From a biological perspective, this was done to obtain CpGs with high sequence conservation, and from a practical perspective it allowed selection of array probes expected to be effective in great apes.

**Figure 1.**
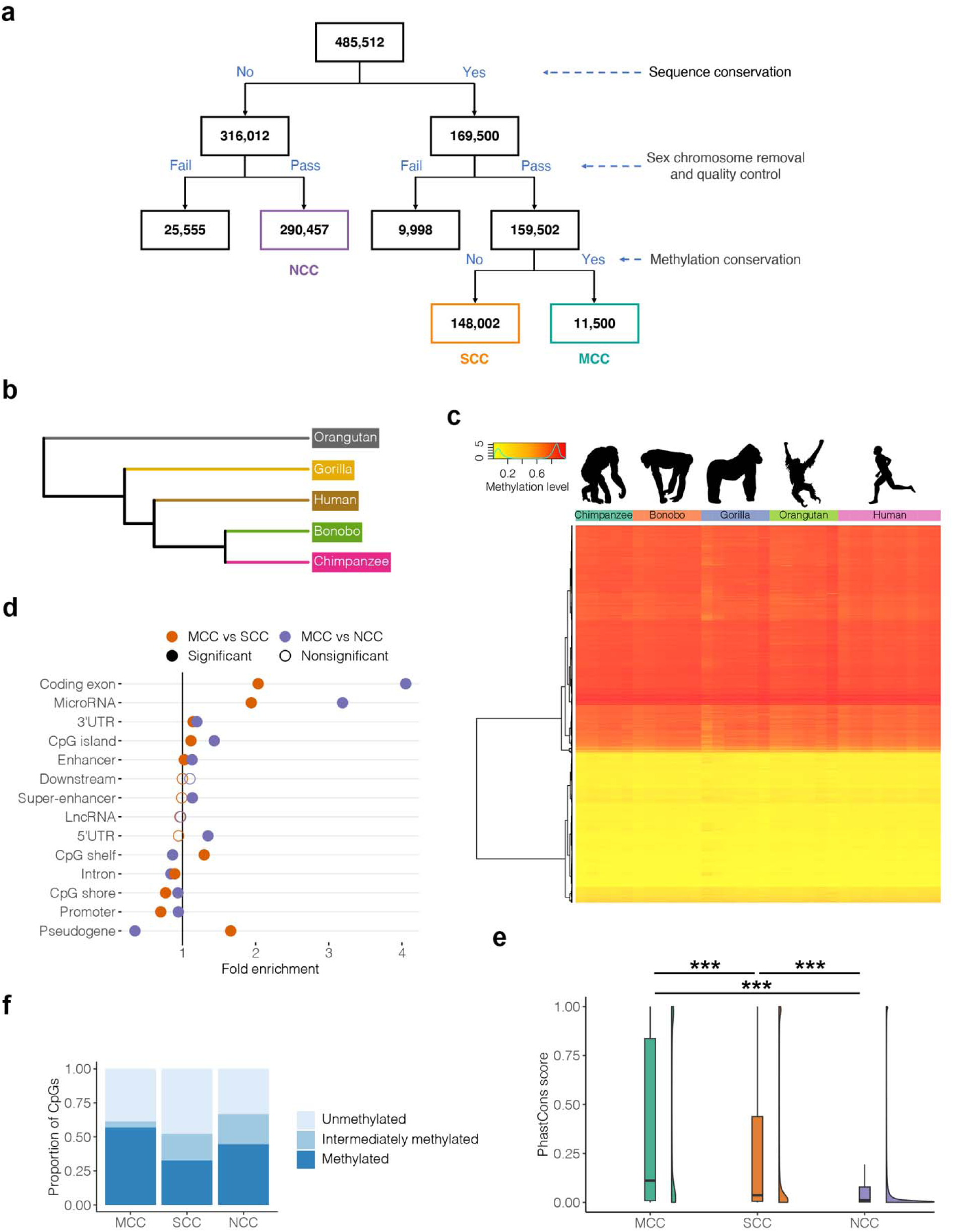
Identification and characterization of MCCs. **(a)** Flowchart of MCC discovery. **(b)** A phylogenetic tree was constructed using methylation information of probes matched to all great ape genomes (*n* = 159,502). **(c)** Methylation pattern of MCCs across great apes. Images of great apes were extracted from Phylopic and are in the public domain (“http://phylopic.org”). **(d)** Significant enrichment and depletion of MCCs in genomic features compared separately to 1000 randomizations of SCCs and NCCs. FDR < 0.05 was considered significant. **(e)** Comparison of CpG sequence conservation between MCCs, SCCs, and NCCs using phastCons scores. * FDR < 0.05; ** FDR < 0.01; *** FDR < 0.001; ^ns^ nonsignificant. **(f)** Proportions of MCCs with methylated (beta > 0.75), unmethylated (beta < 0.25), and intermediately methylated (0.25 < beta < 0.75) methylation patterns in human blood samples separately compared to SCCs and NCCs. The figure represents the proportion of MCCs identified using the threshold of methylation differences < 0.05 and FDR > 0.1.

To determine whether evaluation of DNAm in these CpGs was meaningful from an evolutionary viewpoint, we constructed a phylogenetic tree using the DNAm levels at 159,502 CpGs identified across great apes and filtered by sex chromosome and quality control (including the removal of probes with missing values in > 5% of samples and polymorphic CpG probes; see Methods section “Reconstructing a phylogenetic tree based on DNAm information”). This tree showed bonobo and chimpanzee to be most similar to humans, followed by gorilla and orangutan (Fig. 1b). These relationships of great apes based on DNAm levels in these CpGs were consistent with previous evolutionary relationships estimated using genomic data^34,35^. This confirmed that DNAm patterns at these CpGs can reflect evolutionary history, and provided a basis for identification and characterization of DNAm conservation.

Differences in DNAm patterns in the 159,502 CpGs across great ape species were then calculated to establish which of these loci had conserved DNAm. These so-called MCCs were defined as CpGs with DNAm differences (absolute delta beta) between any two species < 0.05 and statistically nonsignificant according to the Kruskal–Wallis test for DNAm differences across five great ape species with false discovery rate (FDR) > 0.1 (see Methods section “Identifying methylation conservation of CpGs”). Using this approach, we identified 11,500 MCCs (2.56% of all CpGs tested) at a genome-wide level with conservation of both DNAm and sequence (Fig. 1a, c and Supplementary Table S2).

For comparison, two other types of CpGs were also characterized: 148,002 sequence-conserved CpGs (SCCs), defined as CpGs with conserved sequences but nonconserved methylation patterns; and 290,457 nonconserved CpGs (NCCs), defined as CpGs with nonconserved sequences (Fig. 1a and Supplementary Table S2). The comparison of MCCs to SCCs isolated DNAm conservation as the key difference. In comparison to NCCs, MCCs differ in both DNAm and sequence conservation. When characterizing MCCs, these comparisons of groups of CpGs allowed us to associate key findings to conservation of DNAm, sequence, or both. For example, we carried out pathway analysis by comparing genes whose promoters and gene bodies overlapped MCCs to those for SCCs and NCCs separately. When compared to genes associated with SCCs, those associated with MCCs were significantly enriched in crucial human biological pathways involved in biological regulation, cell functions, and development (all Benjamini–Hochberg FDR < 0.05) (Supplementary Table S3). Genes associated with MCCs were enriched in 3624 additional biological pathways in comparison to those associated with NCCs (all Benjamini–Hochberg FDR < 0.05), which included almost all pathways identified when compared to SCCs (*n* = 2951) (Supplementary Table S3).

#### Cross-tissue validation of MCC methylation conservation

To validate the evolutionary conservation of MCC methylation, we examined four additional datasets from whole blood, prefrontal cortex, lateral cerebellum, and dermal fibroblast samples, containing both chimpanzee and human data (data sources are described in Supplementary Table S1). Using the independent whole blood data set, we confirmed that the DNAm patterns were more highly conserved (as measured by absolute values of DNAm differences [delta beta] between chimpanzee and human samples) on average at MCCs than at SCCs between 14 chimpanzees and 20 humans (0.032 vs. 0.051, respectively, *P* < 0.001, 1000 permutation test with randomly shuffled labels of the values in the two groups for comparison between MCCs and SCCs) (Supplementary Fig. S1a). Although MCCs were identified from blood, similar results were also observed between 8 chimpanzee and 7 human prefrontal cortex samples (0.029 vs. 0.041, respectively, *P* < 0.001, 1000 permutation test), between 8 chimpanzee and 7 human lateral cerebellum samples (0.037 vs. 0.046, respectively, *P* < 0.001, 1000 permutation test), and between 7 chimpanzee and 2 human dermal fibroblast samples (0.046 vs. 0.053, respectively, *P* < 0.001, 1000 permutation test), suggesting the potential expansion of DNAm conservation of MCCs to other tissues (Supplementary Fig. S1a). The cross-tissue DNAm conservation in MCCs may be related to their enrichment in genes associated with basic cell functions mentioned above and is consistent with previous findings that evolutionarily conserved genes tend to impact more organs or anatomical systems than evolutionarily new genes^36^.

We further validated MCCs in these data sets with M values (calculated as the log2 ratio of the intensities of methylated probes versus unmethylated probes) to avoid the potential uneven variability of beta values, as the DNAm range of beta values can result in extremes having considerably lower variability^37^. Smaller average values of DNAm differences between chimpanzee and human samples were observed in MCCs rather than SCCs when using M values, suggesting the reliability of our approach for investigation of DNAm conservation (Supplementary Fig. S1b). Detailed results using M values are provided in Supplementary Results.

#### MCCs are enriched in coding and regulatory elements but depleted in promoters and introns

Next, we mapped MCCs to 14 typical human genomic elements to better understand their potential biological roles. Separately compared to the same number of randomly selected SCCs and NCCs through the 1000 permutation test, MCCs were found to show greater enrichment in several genomic elements, including coding exons (fold enrichment = 2.03 and 4.05, respectively, both Benjamini–Hochberg FDR < 0.001), CpG islands (fold enrichment = 1.12 and 1.43, respectively, both Benjamini–Hochberg FDR < 0.001), and enhancers (fold enrichment = 1.02 and 1.13, respectively, Benjamini–Hochberg FDR = 0.008 and 0.001), but were depleted in some other genomic elements, such as promoters (fold enrichment = 0.70 and 0.94, respectively, Benjamini–Hochberg FDR = 0.005 and 0.003) and introns (fold enrichment = 0.90 and 0.84, respectively, both Benjamini–Hochberg FDR < 0.001) (Fig. 1d and Supplementary Fig. S2). These findings were consistent with the previously observed strong evolutionary constraint of DNA sequences in coding regions and enhancers and weak constraint in introns, but inconsistent with the strong constraint of DNA sequences in promoters^38,39^. Thus, the evolutionary conservation of DNAm was not always consistent with that of its corresponding genomic elements and suggested a complex relationship between DNAm and genomic sequence in the context of evolution.

### Evolutionary conservation between DNAm and genomic sequence

#### Methylation conservation is positively associated with CpG sequence conservation

Prior investigations suggested a positive relationship between CpG methylation conservation and its underlying sequence conservation^15–19^ (i.e., CpG and its neighboring nucleotides). To validate this in our study, we compared CpG sequence conservation scores evaluated using phastCons between MCCs and other types of CpGs (see Methods section “Evolutionary conservation score calculation”). The phastCons score (ranging from 0 to 1) represents the probability that a base is in an evolutionarily conserved element^40^. These scores were downloaded from the UCSC Genome Browser and estimated based on sequence alignment across 10 primates: human (*H. sapiens*), chimpanzee (*P. troglodytes*), gorilla (*G. gorilla gorilla*), orangutan (*P. pygmaeus abelii*), rhesus macaque (*Macaca mulatta*), baboon (*Papio hamadryas*), marmoset (*Callithrix jacchus*), tarsier (*Tarsius syrichta*), mouse lemur (*Microcebus murinus*), and bushbaby (*Otolemur garnettii*)^41^. These primates included but were not limited to great apes, as MCCs and SCCs were defined based on highly conserved CpG sequences (i.e., 100% identical CpG sequences) in great apes. As expected, MCCs had the highest phastCons scores, indicating greater conservation than the other two types of CpGs (0.361 vs. 0.255 vs. 0.125, respectively, all Benjamini–Hochberg FDR < 0.001, 1000 permutation test with randomly shuffled labels of the values in the two groups for comparison between any two CpG types) (Fig. 1e). This supported a positive relationship between CpG sequence and DNAm conservation, and demonstrated that sequence conservation at MCCs extends to some extent beyond the great ape lineage.

#### Methylation conservation is positively associated with mQTL sequence conservation

The relationship between CpG sequence and DNAm conservation was unable to explain why phylogenetic relationships among great apes can be successfully captured by DNAm levels in highly sequence-conserved CpGs (Fig. 1b). It has been proposed that DNAm conservation would be associated with the conservation of genetic variation outside of CpG nucleotides^20^. As DNAm variation has been shown to be associated with the most common type of genetic variation, single-nucleotide polymorphisms (SNPs)^42^, we examined the associations between mQTL sequence conservation and DNAm conservation. For this purpose, we first identified *cis-*mQTLs in six human tissues with both DNAm and genotype data. These tissues were tested in our downstream analysis and contained the corresponding genotype data (see Methods section “Genotype data preprocessing”). We identified mQTLs in samples of each tissue by testing SNPs located within ± 10-kb windows of each CpG using linear modeling with adjustments for sex and age and a family-wise error rate (FWER) threshold of 0.05 ^43^ (see Methods section “Mapping of mQTLs”). Through our analyses, we identified 92,577 *cis-*mQTLs in blood, 39,146 in brain, 69,652 in buccal swabs, 16,111 in lymphoblastoid cell lines (LCLs), 12,472 in saliva, and 7590 in skin (data sources are described in Supplementary Table S1). Then, we compared human mQTL sequence conservation scores of great ape and primate genomes evaluated individually using phastCons between MCCs and other types of CpGs. The phastCons scores of great apes were calculated based on the alignments of gorilla, chimpanzee, and human genomes^44^, while the phastCons scores of primates were calculated based on sequence alignment across 10 primates, as mentioned above. mQTLs for MCCs showed 3.05%–4.64% greater sequence conservation in great ape genomes than for SCCs in three tissues and 0.53%–9.68% greater than for NCCs in five tissues with Benjamini–Hochberg FDR < 0.05 (1000 permutation test with randomly shuffled labels of the values in the two groups for comparison between any two mQTL sets, i.e., mQTL sets for MCCs, SCCs, and NCCs) (Table 1). These differences were larger when using phastCons scores of primate genomes, where mQTLs for MCCs were 7.22%–18.68% more highly conserved than those for SCCs in five tissues and 16.00%–45.95% more highly conserved than those for NCCs in all six tissues (Benjamini–Hochberg FDR < 0.05, 1000 permutation test). These findings suggested that mQTL sequence conservation and CpG methylation conservation are positively related (Table 1).

**Table 1.** Comparison of Human mQTL Evolutionary Conservation Scores in Great Apes and Primates Between MCCs, SCCs, and NCCs.

| Species | Tissue | Average score |  |  | FDR |  |  |
| --- | --- | --- | --- | --- | --- | --- | --- |
|  |  | MCC | SCC | NCC | MCC vs. SCC | MCC vs. NCC | SCC vs. NCC |
| Great apes | Blood | 0.190 | 0.191 | 0.189 | 0.724 | 0.015 | 0.003 |
|  | Brain | 0.189 | 0.189 | 0.186 | 0.469 | 0.001 | 0.001 |
|  | Buccal | 0.188 | 0.189 | 0.187 | 0.060 | 0.131 | 0.003 |
|  | LCL | 0.203 | 0.197 | 0.187 | 0.003 | 0.001 | 0.001 |
|  | Saliva | 0.204 | 0.195 | 0.186 | 0.002 | 0.001 | 0.001 |
|  | Skin | 0.203 | 0.194 | 0.186 | 0.005 | 0.001 | 0.001 |
| Primates | Blood | 0.091 | 0.083 | 0.076 | 0.001 | 0.001 | 0.001 |
|  | Brain | 0.089 | 0.083 | 0.074 | 0.001 | 0.001 | 0.001 |
|  | Buccal | 0.087 | 0.081 | 0.075 | 0.001 | 0.001 | 0.001 |
|  | LCL | 0.091 | 0.087 | 0.071 | 0.278 | 0.001 | 0.001 |
|  | Saliva | 0.097 | 0.086 | 0.075 | 0.037 | 0.001 | 0.001 |
|  | Skin | 0.108 | 0.091 | 0.074 | 0.015 | 0.001 | 0.001 |

Abbreviations: LCL, lymphoblastoid cell line; MCC, methylation-conserved CpGs; NCC, nonconserved CpGs; SCC, sequence-conserved CpGs.
Conservation scores in great ape genomes were calculated based on the alignments of human (*Homo sapiens*), chimpanzee (*Pan troglodytes*), and gorilla (*Gorilla gorilla gorilla*) genomes.
Conservation scores in primate genomes were calculated based on the alignments of human (*Homo sapiens*), chimpanzee (*Pan troglodytes*), gorilla (*Gorilla gorilla gorilla*), orangutan (*Pongo pygmaeus abelii*), rhesus macaque (*Macaca mulatta*), baboon (*Papio hamadryas*), marmoset (*Callithrix jacchus*), tarsier (*Tarsius syrichta*), mouse lemur (*Microcebus murinus*), bushbaby (*Otolemur garnettii*) genomes.
The comparisons were analyzed using the 1000 permutation test. The Benjamini–Hochberg FDR < 0.05 was taken to indicate statistical significance.

To better explore their possible relationships, we mapped mQTLs for MCCs to 14 typical genomic features. The analysis was conducted on each tissue separately as the mQTL set identified for MCCs differed between tissues. The top three mapped features in each of the six tissues were consistently introns, super-enhancers, and enhancers (Supplementary Fig. S3). In contrast to random sets of mQTLs for SCCs or NCCs in each tissue, mQTLs for MCCs exhibited 1.07–73.17-fold or 1.21–19.74-fold stronger enrichments in 13 genomic features in at least one tissue with Benjamini–Hochberg FDR < 0.05 (1000 permutation test for per genomic feature, per tissue) (Supplementary Fig. S4). These enriched features included coding exons and a variety of regulatory regions, such as promoters and enhancers, and 3′-UTRs. Although the enrichment results varied with the different tissue data sets, mQTLs for MCCs were most frequently (five of the six tissues) enriched in promoters, CpG islands, super-enhancers, and enhancers, all of which contain dense clusters of transcription factor (TF) binding sites. This was consistent with previous findings regarding the evolutionary relationship between the epigenome and TF binding events at these genomic features^16,18,45^.

### Quantitative nature of MCCs in human populations

#### Global DNAm patterns of MCCs in human blood samples

Investigation of the global distribution of MCCs in nine human blood samples for MCC discovery (accession number GSE41782) according to the methylation status revealed that 6542 (56.9%) MCCs were methylated (beta > 0.75), 4450 (38.7%) were unmethylated (beta < 0.25), and 508 (4.4%) were intermediately methylated (0.25 < beta < 0.75) (Fig. 1f). The proportions of methylated CpGs in SCCs and NCCs were 32.7% and 44.7%, those of unmethylated CpGs were 47.7% and 33.1%, and those of intermediately methylated CpGs were 19.6% and 22.2%, respectively. Compared to SCCs and NCCs separately, MCCs were 1.74-fold and 1.27-fold more methylated and 4.45-fold and 5.05-fold less intermediately methylated, respectively (all Benjamini–Hochberg FDR< 2.2 × 10^−16^, Chi-squared test) (Fig. 1f). These observations indicated that the distributions of DNAm levels of MCCs in human populations differed from those of other CpG types.

Given the association between intermediately methylated CpGs and high DNAm variability^37,46^, we assessed whether the small proportion of intermediately methylated CpGs in MCCs is a result of the DNAm difference threshold (i.e., absolute delta beta of any two species < 0.05) used in MCC identification. To test this, we removed this threshold (i.e., only used the Benjamini–Hochberg FDR threshold for *P*-values calculated by the Kruskal–Wallis rank-sum test) in MCC identification and obtained another set of MCCs (*n* = 15,475). This set contained 8702 (56.2%) methylated, 5015 (32.4%) unmethylated, and 1758 (11.4%) intermediately methylated CpGs (Supplementary Fig. S5). Although the proportion of intermediately methylated CpGs was increased by 2.59-fold following removal of the DNAm difference threshold (*P* < 2.2 × 10^−16^, Chi-squared test), they still represented only a small subset of MCCs. Consistently, MCCs were 1.75-fold and 1.26-fold more methylated and 1.70-fold and 1.95-fold less intermediately methylated compared to SCCs (*n* = 144,027; the numbers also changed as a subset were regarded as MCCs when changing the threshold) and NCCs, respectively (all Benjamini–Hochberg FDR< 2.2 × 10^−16^, Chi-squared test) (Supplementary Fig. S5). Taken together, these observations indicated that the different distributions of DNAm levels of MCCs compared to other types of CpG in human populations were not primarily driven by use of the DNAm difference threshold. Given that intermediate methylation is predominantly tissue-specific^47^, one possible explanation for this difference may be the lower relation of MCCs to tissue specificity, which was consistent with the cross-species MCC conservation observed in multiple tissues. We provided an overview of analysis of the tissue specificity of MCCs in the section “Associations of MCCs with tissue specificity, environmental exposures, and human diseases.” As the DNAm difference threshold is commonly used in epigenome-wide association studies (EWASs) to represent the percentage of a site that has a change in DNAm and to obtain CpGs more likely to be associated with biological rather than technical variation^37,48^, MCCs (*n* = 11,500) identified with this threshold were used in our subsequent analyses.

#### Moderate positive correlation between methylation conservation and DNAm stability

Evolutionarily conserved genomic sequences are subject to strong selection pressure and can be correlated with reduced variation within species^49^. Given the observed positive relationship between sequence conservation and DNAm conservation, it is logical to expect DNAm variability to be correlated with interspecies conservation, and for DNAm in MCCs to be stable across human individuals. In contrast to genetic variation, however, DNAm variability is sensitive to environmental exposures^7^, making it difficult to reach definitive conclusions. To determine whether and how evolutionary conservation is associated with the variability of DNAm within human populations, we used the same whole blood data set as for MCC discovery (accession number GSE41782) to examine the correlation between DNAm conservation (measured by interspecies standard deviation [SD] of beta values) of CpGs that were available in great apes (i.e., MCCs and SCCs) and their DNAm variability (measured by intraspecies SD of beta values) across human individuals. We found a moderate positive rank correlation between evolutionary conservation and DNAm stability within human populations (Pearson’s *r* = 0.50, *P* < 2.2 × 10^−16^; Spearman’s rho = 0.69, *P* < 2.2 × 10^−16^) (Fig. 2a). This correlation was stronger than that observed previously between coding sequence constraint and conservation (Pearson’s *r* = 0.22)^49^, suggesting that it may not be merely due to the relationship between sequence conservation and DNAm conservation.

**Figure 2.**
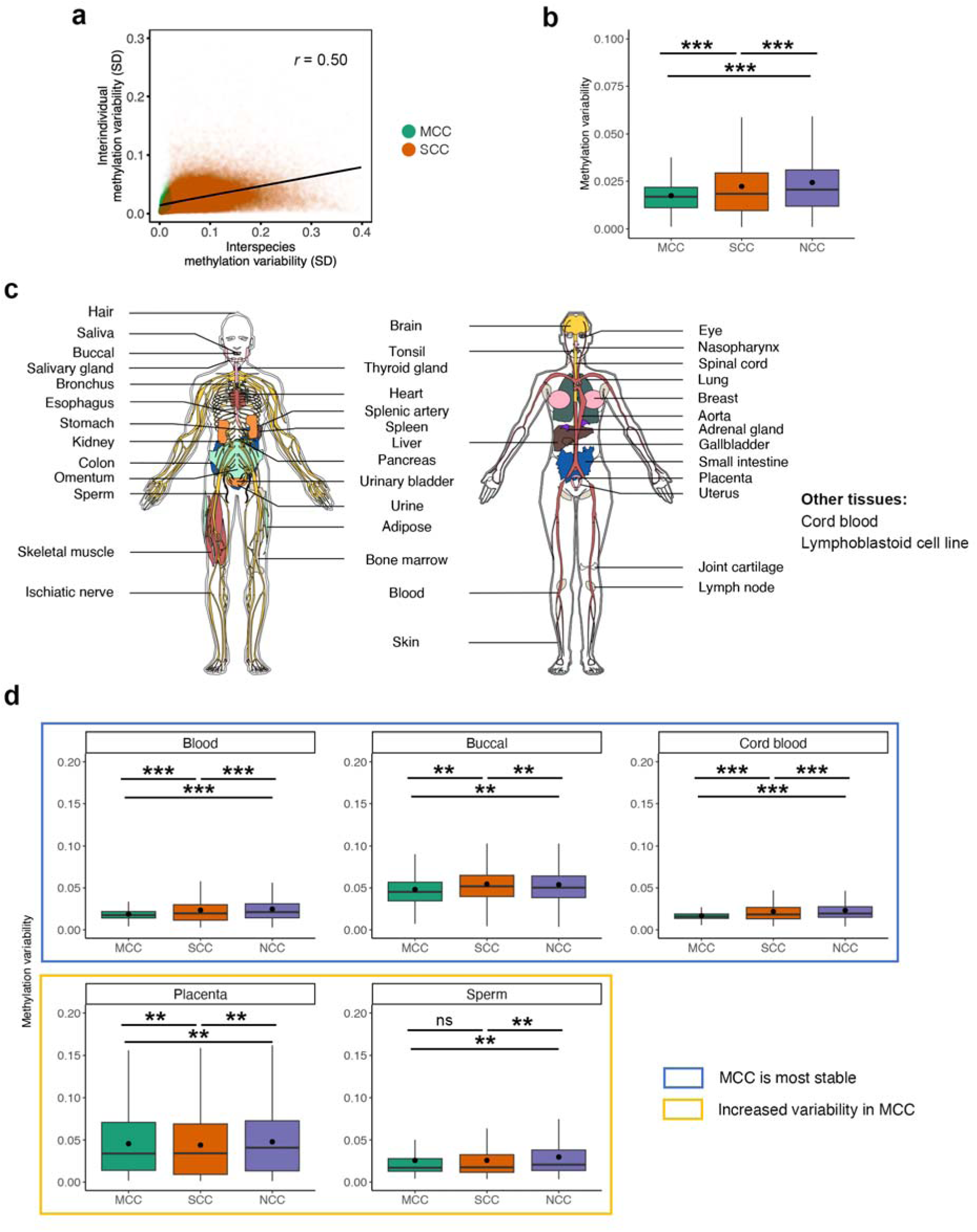
Methylation patterns of MCCs across human individuals in different tissues. **(a)** Smooth scatterplots of intraspecies and interspecies methylation variability within probes matched to all great ape genomes (*n* = 159,502). Methylation variability was calculated as the standard deviation (SD) of CpG methylation across samples. **(b)** Box plots summarizing differential methylation variability between MCCs, SCCs, and NCCs in nine human blood samples. Methylation variability was calculated separately as the SD of CpG methylation across samples. Mean values are presented as circles in each box plot. * FDR < 0.05; ** FDR < 0.01; *** FDR < 0.001; ^ns^ nonsignificant. Outliers are not shown in box plots. **(c)** Illustration of the 42 human tissues included in the analysis. Sample sizes are shown in Supplementary Table S1. **(d)** Box plots summarizing differential methylation variability between MCCs, SCCs, and NCCs in human tissues. MCCs were the most stable in 40 tissues, with three common tissues shown as examples. The complete results are shown in Supplementary Fig. S16. MCCs were not the most stable or variable (defined as increased variability) in placenta and sperm samples. Methylation variability was calculated as the SD of CpG methylation across samples. Mean values are presented as circles in each box plot. Outliers are not shown in box plots.

Although the correlation was significant, its magnitude was moderate. Therefore, we next examined whether and to what extent DNAm variability of MCCs across human individuals would be significantly lower than that of other CpG types. Note that MCC selection was based only on interspecies variability, and was made independently of intraspecies variability. We found MCC methylation to be 22.03% and 28.68% more stable (i.e., with lower variability) on average than SCC and NCC methylation, respectively, across these human blood samples (both Benjamini–Hochberg FDR = 0.001, 1000 permutation test with randomly shuffled labels of the values in the two groups for comparison between any two CpG types) (Fig. 2b). These results showed that evolutionarily conserved DNAm can be characterized by reduced variability within species.

#### Extensive DNAm stability of MCCs in a wide range of healthy human tissues

To further validate MCC methylation stability (measured by intraspecies SD of beta values) in a large data set of blood samples and extend the analysis to other tissues, a total of 10,665 samples from 42 healthy human tissues were examined (Fig. 2c, data sources are described in Supplementary Table S1). Analyses using a data set of > 4000 human blood samples showed that MCCs were consistently 20.41% and 24.57% more stable on average than SCCs and NCCs, respectively (both Benjamini–Hochberg FDR < 0.001, 1000 permutation test with randomly shuffled labels of the values in the two groups for comparison between any two CpG types) (Fig. 2d). Although different blood cell type proportions can influence DNAm^50^, similar trends in MCC methylation stability were found across 28 individuals within each of five major cell types in human peripheral blood (i.e., B cells, CD4^+^ T cells, CD8^+^ T cells, granulocytes, and monocytes) using data from purified cell types, with 14.21%–21.22% and 18.81%–28.15% (depending on cell type) greater stability on average compared to SCCs and NCCs, respectively (all Benjamini–Hochberg FDR < 0.001, 1000 permutation test) (Supplementary Fig. S6a). We then investigated the DNAm variability of MCCs within each of 41 other human tissues and 6 cord blood cell types (B cells, CD4^+^ T cells, CD8^+^ T cells, natural killer cells, granulocytes, and monocytes) across individuals (data sources are described in Supplementary Table S1). We found a similar tendency in all but two tissues, with 1.97%–27.14% and 7.19%–32.04% (depending on tissue and cord blood cell type) greater stability on average compared to SCCs and NCCs, respectively, using a Benjamini–Hochberg FDR threshold of 0.05 via the 1000 permutation test (Fig. 2c and d, Supplementary Fig. S6b and 7). The two exceptions were the placenta and sperm, where MCCs were 4.72% and 13.42% more stable on average than NCCs, respectively (both Benjamini–Hochberg FDR = 1.5 × 10^−3^, 1000 permutation test), but 3.80% more variable (Benjamini–Hochberg FDR = 0.003, 1000 permutation test) and similar to SCCs, respectively (Benjamini–Hochberg FDR = 0.48, 1000 permutation test). Consistent results were also obtained using M values, addressing the concern of whether the relative stability of MCCs could be explained by fewer intermediately methylated CpGs in MCCs compared to SCCs and NCCs separately (Supplementary Fig. S8 and 9; Detailed results are provided in the Supplementary Results). Although MCCs were identified using whole blood samples, the extensive DNAm stability of MCCs relative to SCCs and NCCs in various healthy human tissues was consistent with the cross-species MCC conservation observed in multiple tissues. Taken together, our analyses revealed the important quantitative nature of MCCs, which presented extensive DNAm stability across a wide range of human tissues

### Genetic aspects of MCC methylation in human populations

#### Characteristics of the association between genetic variation and MCC methylation

DNAm variation has been associated with SNPs in human populations^42^. Given the abovementioned relationship between mQTL sequence conservation and CpG methylation conservation across great apes, we further examined whether and how evolutionary conservation would be related to the association between mQTLs and DNAm in human populations. We identified significant associations between CpGs and SNPs in six tissues (methods described in the above section “Methylation conservation is positively associated with mQTL sequence conservation”) and characterized associations in MCCs by comparing them separately to those in SCCs and NCCs. We found that MCCs generally had 9.51%–41.00% and 29.23%–46.41% lower proportions of mQTL-CpGs (depending on tissues) than SCCs in blood, brain, buccal, and saliva and NCCs in all tissues separately with a Benjamini–Hochberg FDR threshold of 0.05 using the two-proportions z-test (Fig. 3a). These observations showed that MCC methylation was less likely to be associated with SNPs, indicating that DNAm conservation is related to the differences in associations between SNPs and DNAm.

**Figure 3.**
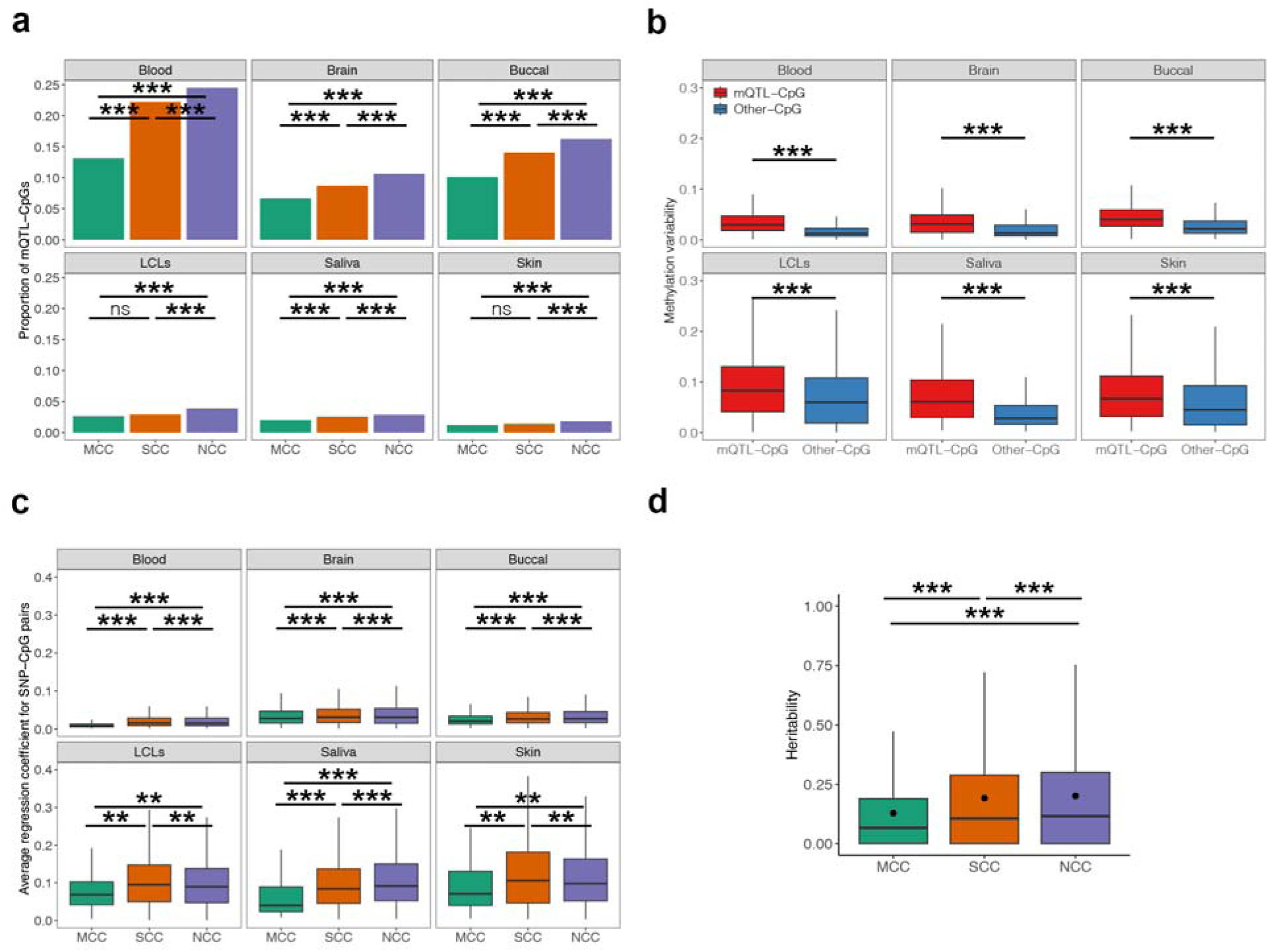
Characterization of genetic impact on MCCs. **(a)** Comparison of mQTL-CpG proportions in MCCs, SCCs, and NCCs in six common tissues. Abbreviation: LCL, lymphoblastoid cell line. The two-proportions z-test was used to compare each pairwise combination of proportions. * FDR < 0.05; ** FDR < 0.01; *** FDR < 0.001; ^ns^ nonsignificant. **(b)** Methylation variability differences between mQTL-CpGs and other CpGs (SNP-independent CpGs) in six common tissues. Methylation variability was calculated as the SD of CpG methylation across samples in each tissue. Outliers are not shown in box plots. **(c)** Comparison of regression coefficient for SNP-CpG pairs in MCCs, SCCs, and NCCs in six common tissues. The absolute values of regression coefficients were used. Outliers are not shown in box plots. **(d)** Box plot summarizing the genetic heritability differences between MCCs, SCCs, and NCCs in a twin cohort. Heritability was estimated by Falconer’s formula [H^2^ = 2(*r*_MZ_ − *r*_DZ_)]. Mean values are presented as circles in each box plot. Outliers are not shown in box plots.

Next, we examined whether the different proportions of CpGs in association with SNPs among the three CpG types would be linked to the differences DNAm variability discussed above. CpGs associated with SNPs showed 31.61%–103.51% (depending on tissue) increased average DNAm variability over SNP-independent CpGs using all CpGs (all Benjamini– Hochberg FDR < 0.001, 1000 permutation test) (Fig. 3b). With regard to each CpG type, MCCs associated with SNPs showed 10.23%–79.68% increased average DNAm variability (depending on tissue) over SNP-independent MCCs, while increases of 30.24%–111.79% and 30.78%– 99.69% were observed for SCCs and NCCs, respectively (all Benjamini–Hochberg FDR < 0.05 with the exception of MCC in skin, 1000 permutation test) (Supplementary Fig. S10). These results suggested a relationship between genetic association and high DNAm variability in CpGs, and provided a potential genetic context for MCC stability in human populations.

Focusing on mQTL-CpG pairs, MCCs were associated with SNPs with smaller effect sizes (absolute value of regression coefficient). Indeed, the average effect sizes of SNPs on MCCs were 11.69%–56.19% and 14.27%–56.72% (depending on tissue) smaller than those on SCCs and NCCs, respectively, across the six tissues examined (all Benjamini–Hochberg FDR < 0.05, 1000 permutation test) (Fig. 3c). The comparison of effect sizes among CpGs may be skewed by the various numbers of SNPs within haplotypes, as numerous linked SNPs within haplotypes are likely to be associated with CpGs simultaneously. To overcome this challenge, we chose a representative SNP (tag-SNP) for all SNPs within each haplotype using PLINK and repeated the analysis^51^. The findings were similar, with effect sizes (absolute value of regression coefficient) 12.55%–53.83% and 16.42%–54.86% (depending on tissue) smaller for MCCs on average than SCCs and NCCs, respectively, across the six tissues (all Benjamini–Hochberg FDR < 0.05, 1000 permutation test) (Supplementary Fig. S11). These observations suggested that the degree of association between genetic variants and DNAm variability differed by DNAm conservation, and the weak effect size of genetic variation on MCC methylation variability may provide an additional potential genetic context for MCC stability in human populations. Notably, these analyses were further validated by converting beta values to M values to reduce the potential confounding effect of uneven DNAm variability across the DNAm beta value range when investigating associations between CpGs and SNPs^37^. Consistent results were obtained (Supplementary Fig. S12; Detailed results are provided in the Supplementary Results).

#### Low heritability of MCCs relative to other CpG types

Although the findings outlined above indicated the association of the most common type of genetic variation, SNPs, with MCC methylation variability, other types of genetic variation, such as structural and copy number variations, which have been reported to be associated with DNAm variability have not been investigated^52,53^. To estimate the global influence of genetic variation on MCCs, we estimated MCC DNAm heritability, which is a statistic used to estimate the proportion of phenotypic variation controlled by genetic variation. Here, we used Falconer’s formula to estimate the broad-sense heritability of CpG methylation in a twin cohort, H^2^ = 2(*r*_MZ_ − *r*_DZ_), where *r*_MZ_ and *r*_DZ_ correspond to the Pearson correlation of DNAm in monozygotic (MZ) pairs and dizygotic (DZ) pairs, respectively^54^. In a whole blood data set containing samples from 426 MZ and 306 same-sex DZ twin pairs (GSE105018), we found that the average heritability was significantly lower for MCCs than for SCCs and NCCs separately using both beta values (0.129 vs. 0.192 vs. 0.201, respectively; all Benjamini–Hochberg FDR < 0.001, 1000 permutation test with randomly shuffled labels of the values in the two groups for comparison between any two CpG types) and beta-converted M values (0.121 vs. 0.185 vs. 0.192, respectively; all Benjamini–Hochberg FDR < 0.001, 1000 permutation test) (Fig. 3d and Supplementary Fig. S13). These observations indicated that MCCs were less dependent on genetic variation and suggested that the association between genetic variation and DNAm differed according to DNAm conservation. These results were consistent with those of the above SNP analysis but extended to the global influence of genetic variation.

### Associations of MCCs with tissue specificity, environmental exposures, and human diseases

The analyses outlined above depicted a comprehensive picture of the quantitative nature of MCCs across individuals in a range of tissues and their underlying genetic bases. As evolutionary conservation is a commonly used indicator of functionally important genomic regions^28^, it is logical to expect MCCs to be associated with important biological functions and human diseases. However, the potential functional relevance of DNAm conservation remains largely unknown. Therefore, it was necessary to examine the functionally important biological processes and human diseases with which MCCs would be associated. To systematically characterize the possible associations of MCCs with specific phenotypes of interest or confounders in EWASs, we performed a diverse suite of assessments ranging from tissue specificity to lifestyle and health factors that have previously been shown to be associated with DNAm patterns^4,5,7,55,56^. For this purpose, in-house and publicly available DNAm data sets with available information on multiple tissues, 49 demographic and environmental factors, 34 cancers, and 69 noncancer diseases were collected (data sources are described in Supplementary Table S1). Diseases were divided into cancers and non-cancer conditions, as aberrant DNAm in cancers has been widely characterized as global loss and focal gain of DNAm^5,6^. In addition, to better explore MCCs on the whole, we calculated the average values of MCC methylation variability/differences for each trait instead of identifying trait-related CpGs in MCCs.

#### MCCs are less likely to be associated with tissue specificity than SCCs and NCCs separately

As the evolutionary conservation and DNAm stability of MCCs relative to SCCs and NCCs can be observed in multiple nonblood tissues and MCCs had a low proportion of intermediately methylated CpGs, we hypothesized that MCCs may be less likely to be associated with tissue specificity than SCCs and NCCs. Therefore, we assessed DNAm variability (measured by the SD of beta values) of each CpG type across distinct human tissues and blood cell types (data sources are described in Supplementary Table S1). Only samples with multiple tissues or cell types collected from the same individual were used in this analysis. As expected, we found that DNAm levels in MCCs were 14.22%–61.94% and 10.84%–61.86% more stable on average across tissues, blood cell types, and cord blood cell types than those in SCCs and NCCs, respectively (all Benjamini–Hochberg FDR < 0.05, 1000 permutation test with randomly shuffled labels of the values in the two groups for comparison between any two CpG types) (Supplementary Fig. S14a). Similar results were also observed in other independent tissue and cell type data sets: MCCs were 8.36%–44.81% and 6.42%–43.89% more stable on average than SCCs and NCCs, respectively, across tissues, blood cell types, and cord blood cell types (all Benjamini–Hochberg FDR < 0.05, 1000 permutation test) (Supplementary Fig. S14b). Interestingly, SCCs showed the greatest DNAm variability among multiple tissues, and were 2.12%–6.31% more variable on average compared to NCCs, suggesting that some SCCs among great apes are likely to be tissue-specific (all Benjamini–Hochberg FDR < 0.05, 1000 permutation test). This was consistent with the previous finding that rat tissue-specific differentially methylated regions for whole blood, whole brain, and sperm were mostly in regions orthologous in mice and humans^19^. Similar results were observed when using M values instead of beta values (Supplementary Fig. S15; Detailed results are provided in the Supplementary Results).

#### MCCs are more likely to be associated with certain cancers than SCCs and NCCs, but not with demographic and environmental factors or noncancer diseases

To characterize the possible associations of MCCs as a whole with each specific demographic and environmental factor and disease, we calculated their differences in DNAm (defined as the absolute delta beta) between phenotypic groups of each factor or disease. With regard to demographic and environmental factors, the average DNAm differences for MCCs according to these factors were consistently less than 0.03, which is a commonly used DNAm change threshold for EWASs^57^ (Supplementary Fig. S16 and Table S4). Similarly, the average DNAm differences for MCCs were less than 0.03 in almost all noncancer diseases examined (66/69) (Supplementary Fig. S17a). Only three noncancer diseases had average MCC methylation differences slightly greater than 0.03 (atherosclerotic lesions, 0.035; psoriasis, 0.032; and polymicrogyria, 0.039). These results suggested that the DNAm differences in MCCs associated with these factors were consistently small on average.

To further determine if the considerably small DNAm changes in these phenotypic factors and noncancer diseases represent a specific property of MCCs or a general property of all CpGs, we then examined whether the average DNAm differences in these factors were smaller in MCCs than SCCs and NCCs separately. For almost all demographic and environmental factors (44/49) and noncancer diseases examined (65/69), MCCs showed significantly 2.45%–42.85% and 4.96%–44.28% (depending on factors or noncancer diseases) smaller average DNAm differences than SCCs and NCCs, respectively, with a Benjamini–Hochberg FDR threshold of 0.05 using the 1000 permutation test (Fig. 4a). Taken together, these findings suggested that stable DNAm patterns of MCCs relative to SCCs and NCCs are maintained in the presence of demographic and environmental factors as well as noncancer diseases.

**Figure 4.**
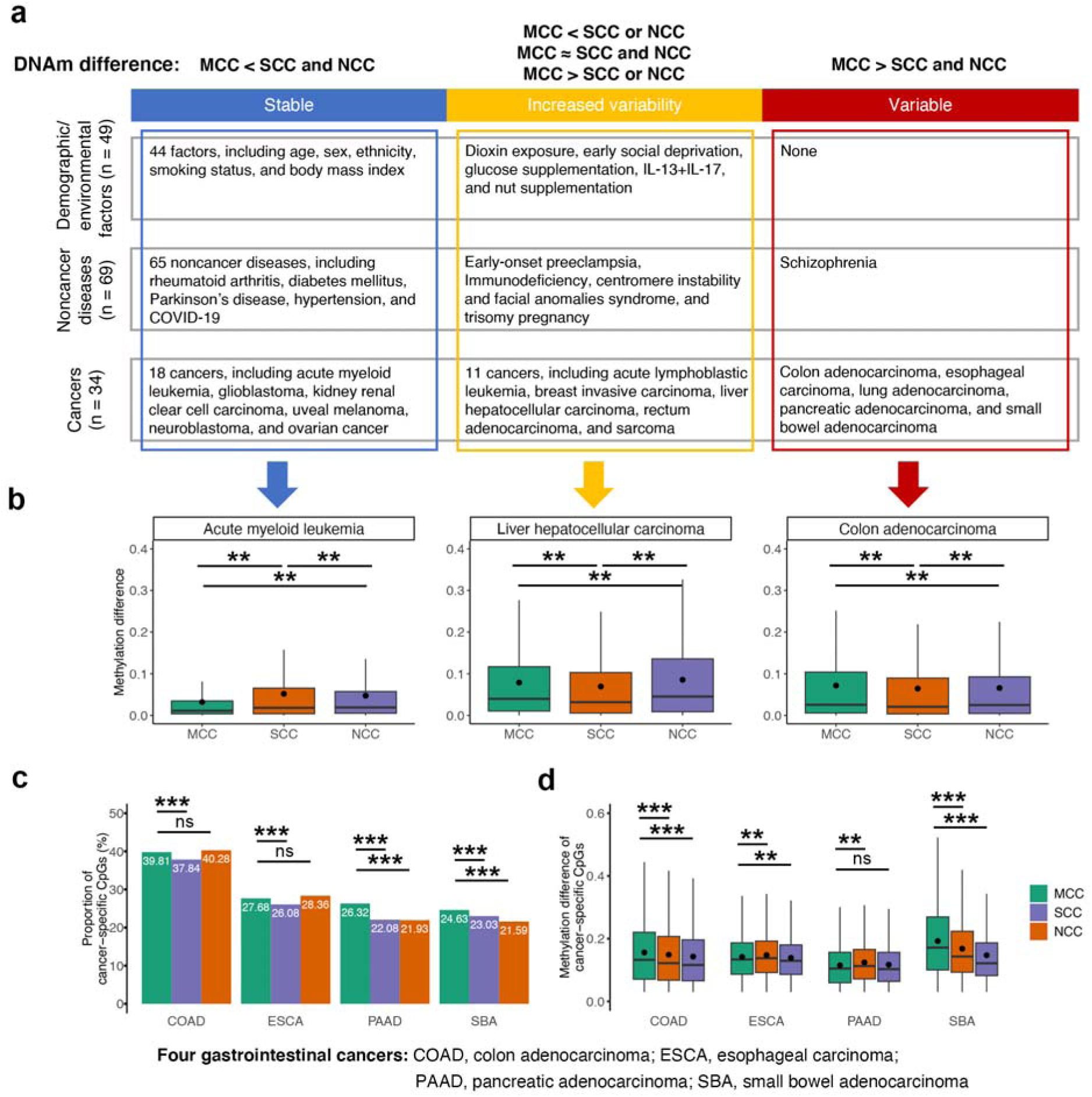
Associations of MCCs with various demographic/environmental factors and diseases. **(a)** Comparison of MCCs with SCCs and NCCs separately with regard to methylation differences (absolute beta value changes) between groups according to factors divided into three types. Abbreviations: DNAm, DNA methylation; IL-13, interleukin-13; IL-17, interleukin-17. **(b)** Box plots showing three examples. Mean values are represented as circles in each box plot. * FDR < 0.05; ** FDR < 0.01; *** FDR < 0.001; ^ns^ nonsignificant. Outliers are not shown in box plots. **(c)** Four gastrointestinal cancers were used as examples to depict the proportions of cancer-specific CpGs among MCCs, SCCs, and NCCs. Cancer-specific CpGs were identified using thresholds of FDR < 0.05 and methylation difference > 0.03. The two-proportions z-test was used to compare each pairwise combination of proportions. **(d)** Four gastrointestinal cancers were used as examples to depict the methylation variability of cancer-specific CpGs among MCCs, SCCs, and NCCs. Mean values are presented as circles in each box plot. Outliers are not shown in box plots.

However, large average methylation differences (> 0.03) for MCCs between primary tumors and normal tissues were found in nearly all cancer types (33/34, except thyroid carcinoma [THCA]), ranging from 0.017 to 0.080, which was consistent with previous reports of extensive aberrant DNAm in malignancies^5,6^ (Supplementary Fig. S17b). To further determine if the considerably large DNAm changes in cancers represent a specific property of MCCs or a general property of all CpGs, we further separately compared MCC methylation differences to those for SCCs and NCCs. Unlike their stability in almost all demographic and environmental factors and noncancer diseases examined, we found that average MCC methylation differences were lower in only approximately half of all cancer types (18/34) compared to SCCs and NCCs, with a Benjamini–Hochberg FDR threshold of 0.05 using the 1000 permutation test (Fig. 4a and b). In particular, MCCs showed 4.43%–14.70% and 2.10%–28.92% greater average DNAm differences than SCCs and NCCs, respectively, in five different cancer types, i.e., colon adenocarcinoma (COAD), esophageal carcinoma (ESCA), pancreatic adenocarcinoma (PAAD), small bowel adenocarcinoma (SBA), and lung adenocarcinoma (LUAD) (all Benjamini– Hochberg FDR < 0.05, 1000 permutation test) (Fig. 4a and b). Taken together, these findings suggested that MCCs tend to be more likely than SCCs and NCCs to be differentially methylated in certain malignancies, but not across demographic and environmental variables, or noncancer disorders.

To be noted, these analyses were further validated by converting beta values to M values to reduce the potential confounding effect of uneven DNAm variability across the DNAm beta value range when investigating associations between CpGs and SNPs^37^. We observed comparable results (Supplementary Fig. S18 and 19; Detailed results are provided in the Supplementary Results).

The stability of DNAm patterns of MCCs in demographic and environmental factors was unsurprising, as we observed their stability across human individuals. The varied average DNAm differences in MCCs between noncancer and cancer diseases may be explained by the different roles of MCCs in the proposed mechanisms of these diseases. As illustrations, we discussed some examples. First, some age-associated diseases, such as Alzheimer’s disease and Parkinson’s disease, have been shown to be associated with the age-related accumulation of epigenetic alterations^58^. The stability of MCCs in these diseases is consistent with the observed stability of MCCs relative to other CpG types in aging. In addition, given that MCCs are less reliant on genetic variation than other CpG types, DNAm stability may be expected in certain genetic disorders, including ankylosing spondylitis and Huntington’s disease. As a final example, the role of DNAm alterations in cancer stem cells (CSCs) has been recognized as one possible mechanism underlying the association between DNAm alterations and oncogenesis^58^. Genes whose promoters and gene bodies overlapped MCCs were found to be enriched in a number of signaling pathways involved in the maintenance of stem cell properties and survival of CSCs through our above pathway enrichment analysis compared to genes associated with SCCs and NCCs (Supplementary Table S3). These pathways in CSCs included the Wnt, Notch, and PI3K-Akt signaling pathways, epithelial-mesenchymal transition, as well as stem cell differentiation, development, and proliferation^59,60^. This may partly explain the increased DNAm differences in MCCs between tumors and normal tissues. Further exploration of the roles of MCCs in cancer biology will be introduced in the next section with integration of gene expression data.

#### Characteristics of the associations between MCCs and gastrointestinal cancers

Although large average DNAm differences between primary tumors and normal tissues (> 0.03) for MCCs were found in nearly all cancer types tested due to extensive aberrant DNAm in malignancies described previously^5,6^, MCCs showed the greatest DNAm differences (absolute delta beta) among the three CpG types only in five cancer types. Notably, four of these five malignancies (COAD, ESCA, PAAD, and SBA) were gastrointestinal cancers, which are cancers that develop along the gastrointestinal tract from the esophagus to the anus and account for 26% of the global cancer incidence and 35% of all cancer-related deaths^61^. The other two gastrointestinal cancers tested here, liver hepatocellular carcinoma (LIHC) and rectum adenocarcinoma (READ), also showed higher average DNAm differences (absolute delta beta values) in MCCs than SCCs but not NCCs (for LIHC: 0.079 vs. 0.069 vs. 0.086, respectively, Benjamini–Hochberg FDR = 0.003 and 1.5 × 10^−3^; for READ, 0.073 vs. 0.067 vs. 0.072, respectively, Benjamini–Hochberg FDR = 0.001 and 0.263) (Fig. 4a and b). In comparison to other cancers (i.e., non-gastrointestinal cancers), MCCs were 16.79-fold more likely to show the greatest DNAm differences between tumors and normal samples in gastrointestinal cancers (*P* = 0.01, Fisher’s exact test). This was consistent with a previous study showing that biologically similar cancers may share more similar epigenetic mechanisms^62^.

Large average DNAm differences in MCCs for one trait are expected to account for a high proportion and/or large effect size (i.e., greater absolute values of delta-beta) of trait-related CpGs. To further elucidate the associations of MCCs with gastrointestinal cancer, we subsequently examined cancer-specific CpGs in four cancers, COAD, ESCA, PAAD, and SBA, with a threshold of Mann–Whitney U test Benjamini–Hochberg FDR < 0.05 and DNAm difference > 0.03 (see Methods section “Identification of cancer-specific CpGs”). The results showed that MCCs had 5.23%–19.20% and 14.07%–20.01% higher proportions of gastrointestinal cancer-specific CpGs than SCCs in all four cancers examined and NCCs in PAAD and SBA, respectively, with a Benjamini–Hochberg FDR threshold of 0.05 using the two-proportions z-test (Fig. 4c). In addition, gastrointestinal cancer-specific CpGs for MCCs presented 4.97%–14.52% and 2.33%–30.68% greater average DNAm changes (absolute delta beta values) than those for SCCs in COAD and SBA and NCCs in COAD, ESCA, and SBA, respectively, with a Benjamini–Hochberg FDR threshold of 0.05 using the 1000 permutation test (Fig. 4d). The observation that MCCs were slightly more likely to be or/and were more strongly associated with these gastrointestinal cancers than other types of CpG, supported the suggestion that the complex combination of proportion and effect size of trait-related MCCs can be reflected in average DNAm differences in MCCs for a given trait. Similar results were also determined when using different thresholds of DNAm difference (i.e., 0, 0.05, and 0.1) with a combination of Benjamini–Hochberg FDR < 0.05 in cancer-specific CpG identification (Supplementary Table S5).

### MCC methylation, gene expression, and functional relevance

#### Characteristics of the association between MCC methylation and gene expression in cancer

Due to the well-recognized role of DNAm in modification of gene expression^2^, to better characterize the functional roles of MCCs in cancer, especially in gastrointestinal cancer, we leveraged the availability of TCGA RNA-sequencing data from the same individuals with 13 cancer types with sample sizes ranging from 140 to 448 to separately identify genes whose expression was associated with MCC methylation (data sources are described in Supplementary Table S1). The 13 TCGA cancers included four gastrointestinal cancers (COAD, ESCA, LIHC, and PAAD) and nine non-gastrointestinal cancers (bladder urothelial carcinoma [BLCA], breast invasive carcinoma [BRCA], head and neck squamous cell carcinoma [HNSC], kidney renal clear cell carcinoma [KIRC], kidney renal papillary cell carcinoma [KIRP], prostate adenocarcinoma [PRAD], sarcoma [SARC], THCA, and uterine corpus endometrial carcinoma [UCEC]). We associated the levels of expression of 24,144 genes in each cancer type with those of DNAm (M values) at CpGs located within their promoters and gene bodies, for a total of 320,740 CpG sites (9077 for MCCs, 110,877 for SCCs, and 200,786 for NCCs). Using linear regression with adjustment for age, sex, and disease status (i.e., tumor or normal tissue) and a Benjamini-Hochberg FDR threshold of 0.05 in each cancer type (see Methods section “Association of DNAm with gene expression”), we identified 71,857–145,702 (2044–4487 for MCCs, 27,183–53,800 for SCCs, and 42,630–87,415 for NCCs) CpGs whose DNAm levels were significantly associated with the expression of 13,253–15,993 genes across these cancer types (Supplementary Table S6).

Comparing proportions of CpGs whose levels of DNAm were associated with gene expression in three CpG types, we found that the proportion in MCCs was the lowest in COAD, KIRC, KIRP, PRAD, and THCA, and the highest in ESCA, with a Benjamini–Hochberg FDR threshold of 0.05 using the two-proportions z-test (Supplementary Fig. S20). The remaining comparisons were not statistically significant for MCCs compared to SCCs and/or NCCs. Similar trends were also observed in the comparison of coefficients of determination (R^2^) for CpG-gene pairs in MCCs, SCCs, and NCCs in the 13 cancers: MCCs showed the lowest R^2^ values in KIRC, KIRP, and THCA, but the largest R^2^ values in ESCA and LIHC with a Benjamini–Hochberg FDR threshold of 0.05 using the 1000 permutation test (Fig. 5a). Notably, ESCA and LIHC were gastrointestinal cancers that presented the largest DNAm differences in MCCs. Taken together, MCCs were more likely to be or were more strongly associated with gene expression than SCCs and NCCs separately in certain gastrointestinal cancers, suggesting the importance of the regulatory role of MCCs in particular gastrointestinal cancers.

**Figure 5.**
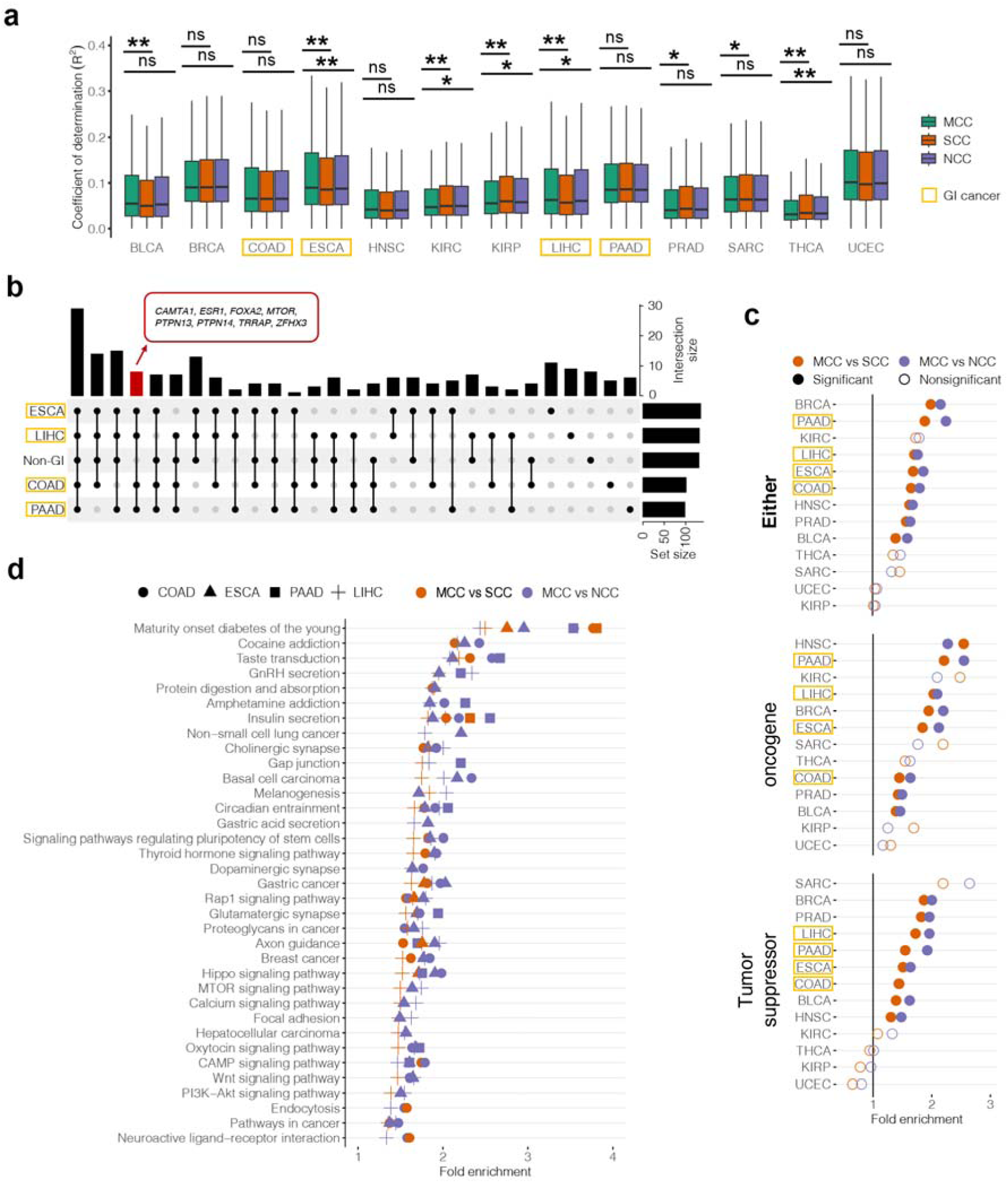
Associations between DNA methylation and the expression of genes with diverse biological functions. Abbreviations: BLCA, bladder urothelial carcinoma; BRCA, breast invasive carcinoma; COAD, colon adenocarcinoma; ESCA, esophageal carcinoma; GI, gastrointestinal; HNSC, head and neck squamous cell carcinoma; KIRC, kidney renal clear cell carcinoma; KIRP, kidney renal papillary cell carcinoma; LIHC, liver hepatocellular carcinoma; PAAD, pancreatic adenocarcinoma; PRAD, prostate adenocarcinoma; SARC, sarcoma; THCA, thyroid carcinoma; UCEC, uterine corpus endometrial carcinoma. **(a)** Comparison of coefficient of determination (*R*^2^) for CpG-gene pairs in MCCs, SCCs, and NCCs in the 13 cancers. These cancers included four gastrointestinal cancers (COAD, ESCA, LIHC, and PAAD) and nine non-gastrointestinal cancers (BLCA, BRCA, HNSC, KIRC, KIRP, PRAD, SARC, THCA, and UCEC). Outliers are not shown in box plots. * FDR < 0.05; ** FDR < 0.01; *** FDR < 0.001; ^ns^ nonsignificant. **(b)** An upset plot of cancer driver genes across genes whose expression is associated with MCC methylation in various cancers. Non-gastrointestinal cancer (non-GI) is a combination of pathways from nine non-gastrointestinal cancers (BLCA, BRCA, HNSC, KIRC, KIRP, PRAD, SARC, THCA, and UCEC). Cancer driver genes for at least one of the 73 cancer types in Integrative OncoGenomics were used. The red bar shows the unique cancer driver genes shared in gastrointestinal cancers. **(c)** Significant enrichment of genes whose expression is associated with MCC methylation in cancer-specific cancer driver genes. FDR < 0.05 is considered significant. **(d)** Shared KEGG pathways for genes whose expression is associated with MCC methylation in four gastrointestinal cancers. Detailed pathway information is shown in Supplementary Table S9.

#### Genes whose expression was associated with MCC methylation are enriched in cancer driver genes

As DNAm-mediated modulation may be an important mechanism influencing the regulation of cancer driver genes^63^, to obtain a snapshot of the potential roles of genes with association between MCC DNAm and expression in cancers, we mapped these genes to cancer driver genes, mutations of which affect cancer progression^64^. The cancer driver gene set was collected from Integrative OncoGenomics, which is a database of the compendium of mutational driver genes across 73 tumor types (Release date: 2023.05.31)^64^. We showed that 4.76%–6.01% of genes whose expression was significantly associated with MCC methylation across the 13 cancers were cancer driver genes for at least one of 73 cancer types in Integrative OncoGenomics (Supplementary Table S7). Compared to random genes in RNA-sequencing data, genes whose expression was associated with MCC methylation showed strong enrichment of 1.67–2.11-fold among cancer driver genes, depending on cancer type (all Benjamini–Hochberg FDR < 0.001, 1000 permutation test). This suggested that MCC methylation may play an important role in the regulation of cancer driver genes.

These cancer driver genes with significant association between MCC methylation and expression shared at most 5 cancer types, and none were shared among the 13 cancers. Given that MCCs were more likely to present the greatest DNAm differences between tumors and normal tissue samples in gastrointestinal cancers, we next examined whether any of these cancer driver genes were shared in gastrointestinal cancers. We found that 37 cancer driver genes were associated with MCCs in all four gastrointestinal cancers, with eight genes (*CAMTA1*, *ESR1*, *FOXA2*, *MTOR*, *PTPN13*, *PTPN14*, *TRRAP*, *ZFHX3*) uniquely identified in gastrointestinal cancers (i.e., not identified in any of the nine non-gastrointestinal cancers) (Fig. 5b and Supplementary Table S8). Focusing on cancer-specific driver genes (e.g., ESCA driver genes were used for mapping genes associated with MCCs in ESCA), *GNAS* was the only gene shared among the four gastrointestinal cancers and was also found in three non-gastrointestinal cancers (BRCA, HNSC, and PRAD). *GNAS* can activate the cAMP-PKA signaling pathway and hence stimulate mucin secretion, thus providing a crucial link between inflammation and cancer^65,66^. Given that the integrity of the mucus barrier plays a primary role in protection of the gastrointestinal tract, it was unsurprising that *GNAS* was a shared cancer driver gene expression of which was associated with MCCs in the four gastrointestinal cancers.

Separately compared to genes whose expression was significantly associated with SCC and NCC methylation, genes with associations involving MCCs were significantly enriched (1.39– 1.99-fold and 1.59–2.25-fold) in cancer-specific driver genes in eight cancers, with a Benjamini– Hochberg FDR threshold of 0.05 using the 1000 permutation test (Fig. 5c). Of the eight cancer types, six (except BRCA and PRAD) exhibited larger DNAm differences in MCCs compared to SCCs and/or NCCs. Notably, all four gastrointestinal cancers tested showed significant enrichment of genes whose expression was associated with MCC methylation in cancer driver genes. In addition, nonsignificant enrichment was observed in five cancer types, four of which (except SARC) exhibited the lowest DNAm differences in MCCs compared to SCCs and NCCs. The high but not total concordance between MCC methylation differences in cancer and enrichment of their corresponding genes in cancer driver genes suggested that MCCs may be related to the pathogenesis of multiple cancers, especially gastrointestinal cancers, partly through regulation of cancer driver gene transcription.

Cancer driver genes can be functionally classified as tumor suppressors or oncogenes based on their role in cancer progression^64^. To better decipher the role of MCCs in oncogenesis, we next compared the enrichment of genes with associations involving MCCs, SCCs, and NCCs in the different roles of cancer driver genes. We demonstrated similar MCC enrichment results as described above (Fig. 5c), suggesting that aberrant DNAm in MCCs may be associated with cancer progression through oncogene activation and tumor suppressor gene silencing. Particularly, the CpG island methylator phenotype (CIMP) is a proposed epigenetic mechanism for cancer biology, as it is associated with the oncogenesis and progression of cancer by transcriptionally silencing tumor suppressor genes through promoter hypermethylation^67^. We found that MCCs were more enriched than other CpG types in the regulation of tumor suppressor genes in multiple malignancies, consistent with reports of CIMP.

#### Functional enrichment of genes whose expression was associated with MCC methylation in biological pathways

Pathway enrichment analysis for genes whose promoter and gene bodies overlapped MCCs showed enrichment in cancer-related pathways, including signaling pathways for CSCs. However, not all CpGs in gene promoters and gene bodies are expected to be associated with gene expression, and these associations may be tissue-specific^63,68^. Here, we performed enrichment analysis for genes whose expression was significantly associated with MCC methylation identified in each cancer type separately and performed comparative pan-cancer pathway analysis, which revealed possible common cancer and specific biological processes for MCCs. Applying a Benjamini–Hochberg FDR threshold of 0.05, these genes showed greater enrichment in varied numbers of biological pathways compared to genes whose expression was associated with SCC methylation (*n* from 409 to 823) and NCC methylation (*n* from 631 to 1228) separately across the 13 cancer types (fold enrichment: 1.01–6.51 and 1.02–7.79, respectively) (Supplementary Table S9). Among them, numerous pathways were shared among multiple cancer types. For example, 35 KEGG pathways shared by multiple gastrointestinal cancers, including CSC signaling pathways: Wnt and PI3K-Akt signaling pathways (Fig. 5d).

We next investigated whether any of these pathways were shared in four gastrointestinal cancers or even in the 13 cancers tested. We discovered that 315 and 508 pathways were shared among four gastrointestinal cancer types when compared separately to genes with associations involving SCCs and NCCs (Supplementary Table S10). In light of our earlier finding that the largest DNAm changes in MCCs were more likely to occur in gastrointestinal malignancies, we hypothesized that gastrointestinal cancers may share more biological processes involving MCCs. For this purpose, we assessed shared pathways from four randomly selected cancers of the 13 cancers tested and calculated the proportion of shared pathways to the minimum number of pathways in randomly selected cancers. This process was repeated 1000 times. As expected, we found a significantly higher proportion of shared pathways among gastrointestinal cancers than randomly selected cancers (compared to genes for SCCs: 77.02% vs. 58.32%, respectively, *P* < 0.001; compared to genes for NCCs, 73.73% vs. 60.73%, respectively, *P* < 0.001). This may provide an explanation for the enrichment of MCCs exhibiting the greatest differential DNAm in gastrointestinal cancers.

Although these pathways were shared in gastrointestinal cancers, they were also observed in at least one non-gastrointestinal cancer, i.e., none was exclusive to gastrointestinal cancers (Supplementary Fig. S21). Of them, 163 and 261 pathways were enriched for genes whose expression was significantly associated with MCC methylation in all 13 cancer types tested when compared separately to genes with associations involving SCCs and NCCs, including cellular development processes (e.g., cell morphogenesis, cell development, and differentiation), transcriptional regulation (e.g., transcription *cis*-regulatory region binding, DNA-binding transcription factor activity, and regulation of transcription by RNA polymerase II), and canonical cancer pathways (e.g., the Wnt signaling pathway, cell population proliferation, blood vessel development, and cell migration) (Supplementary Table S11). Consistent with our above findings using genes whose promoters and gene bodies overlapped MCCs, genes whose expression was associated with MCC methylation were significantly enriched in numerous CSC signaling pathways shared among the 13 cancers tested. These observations showed that MCCs may play important roles in shared biological processes across cancers, suggesting the extensive functional relevance of MCCs to cancer.

In contrast to pathways shared by multiple cancers, 395 and 514 enriched pathways for genes whose expression was significantly associated with MCC were specific to each cancer type when compared to genes whose expression was significantly associated with SCC and NCC methylation, respectively, such as the regulation of histone acetylation in ESCA, the insulin-like growth factor receptor signaling pathway in LIHC, and the HIF-1 signaling pathway in UCEC (Supplementary Table S12). These observations suggested that MCCs may also be associated with cancer-specific functions. Taken together, our results suggested that MCCs may be involved in common and specific biological processes across cancers and play extensive roles in cell development, transcriptional regulation, and tumorigenesis.

### Application and extension of DNAm evolutionary conservation

#### Building a user-friendly web interface to integrate our findings

EWASs are frequently used to detect genome-wide epigenetic variants (predominantly DNAm) that are significantly associated with phenotypes of interest. However, the rapid interpretation of EWAS results is often complex and requires additional annotation sources. Identification of the quantitative nature of evolutionarily conserved CpGs among human individuals and their relations to human disease provided an opportunity to develop a community resource for the exploration and interpretation of DNAm signatures of interest from an evolutionary standpoint. Therefore, we developed a user-friendly R Shiny web interface (QNMEC [Quantitative Nature of Methylation under Evolutionary Conservation]; https://epigenome.shinyapps.io/QNMEC/) to depict CpG-based results of comparative epigenomics analyses and EWASs. QNMEC includes DNAm variability across individuals in 42 tissues and DNAm associations with 152 specific phenotypes, at the CpG-type level (i.e., MCCs, SCCs, and NCCs). More importantly, this tool provides information regarding both methylation and sequence conservation for each HM450K CpG, which may be leveraged to infer the potential functional importance of DNAm alterations in EWASs (see Discussion section).

#### An example of DNAm conservation annotation of EWAS findings

To illustrate the application of DNAm conservation, we used a previous EWAS that identified 1858 CpGs associated with atherosclerosis^69^. Based on evolutionary conservation information in QNMEC, 63 of these CpGs were annotated as MCCs and 601 were annotated as SCCs (Supplementary Table S13). Compared to the SCCs associated with atherosclerosis, these MCCs were strongly enriched at the promoters and gene bodies of genes involved in cortisol synthesis and secretion (fold enrichment = 24.53, Benjamini–Hochberg FDR = 0.044), Cushing syndrome (fold enrichment = 14.72, Benjamini–Hochberg FDR = 0.044), and vasopressin-regulated water reabsorption (fold enrichment = 24.53, Benjamini–Hochberg FDR = 0.044). The three biological pathways were linked to glucocorticoid homeostasis, which plays critical roles in atherosclerosis^70,71^. More enriched biological pathways were found when comparing these MCCs to NCCs that were associated with atherosclerosis, i.e., including the three pathways observed above and an additional nine pathways linked to atherosclerosis, such as thyroid hormone synthesis^72^ (fold enrichment = 22.42, Benjamini–Hochberg FDR = 0.048) and insulin secretion^73^ (fold enrichment = 14.95, Benjamini–Hochberg FDR = 0.049) (Supplementary Table S14). There was no significant enrichment when comparing SCCs that were associated with atherosclerosis to those for NCCs. In this example, QNMEC can effectively pick candidate CpGs most likely involved in the pathogenesis of atherosclerosis and markedly reduce the number of candidate CpGs tested in downstream experiments. Therefore, annotation of epigenetic alterations found by EWASs using QNMEC can help prioritize candidate CpGs and advance our understanding of their potential functional roles.

#### Depicting the evolutionary conservation of CpGs on the EPIC array

To further facilitate their use by the epigenetics community, evolutionary conservation information for EPIC CpGs was identified and provided in QNMEC. The EPIC array measures DNAm at over 850,000 CpG sites and shares ∼450,000 of the CpG sites represented on the HM450K array. Therefore, we determined the conservation of additional CpGs (*n* = 413,406) on the EPIC array by aligning probe sequences to great ape genomes and then computing the DNAm differences between species using a data set consisting of eight chimpanzee and seven human prefrontal cortex samples (GSE154403; see Methods “Identifying sequence conservation of CpGs” and “Identifying methylation conservation of CpGs” sections). As described for HM450K CpGs, we classified EPIC CpGs into three types based on the same thresholds of sequence identify and between-species methylation differences; 43,774 CpGs on the EPIC array were defined as MCCs, 75,008 as SCCs, and 285,580 as NCCs (Supplementary Fig. S22). The DNAm conservation of MCCs was further confirmed in the lateral cerebellum samples (eight chimpanzee and seven human samples) in the same data set and yielded consistent results using both beta values and beta-converted M values (beta value: 0.042 vs. 0.091 vs. 0.155, respectively; M value: 0.435 vs. 0.947 vs. 1.356, respectively, all Benjamini–Hochberg FDR < 0.001, 1000 permutation test) (Supplementary Fig. S23).

For these additional EPIC CpGs, we found a significant positive correlation between DNAm conservation among great apes (interspecies SD) and DNAm variability across human individuals (intraspecies SD) using the same data set (Pearson’s *r* = 0.36, *P* < 2.2 × 10^−16^; Spearman’s rho = 0.48, *P* < 2.2 × 10^−16^) (Supplementary Fig. S24a). Consequently, MCCs were more stable than SCCs and NCCs separately in human prefrontal cortex samples (0.033 vs. 0.043 vs. 0.101, respectively, all Benjamini–Hochberg FDR < 0.001, 1000 permutation test) (Supplementary Fig. S24b). Similar results were also found in our in-house EPIC data with ∼500 blood samples (0.016 vs. 0.022 vs. 0.023, respectively, all Benjamini–Hochberg FDR < 0.001, 1000 permutation test) and recapitulated after the conversion of beta values to M values (0.392 vs. 0.475 vs. 0.457, respectively, all Benjamini–Hochberg FDR = 1.5 × 10^−3^, 1000 permutation test) (Supplementary Fig. S25). Taken together, these results showcased the reliability of our approach for the investigation of DNAm conservation and provided a basis for extending our findings on the quantitative nature of MCCs from HM450K to EPIC arrays.

## Discussion

Studying these conserved sequences is essential to gain an understanding of the genetic basis of health and disease, and it has important implications for diagnostics, therapeutics, and personalized medicine^28,74^. Although species-specific epigenetic variation has been suggested to be associated with species-specific adaptations and phenotypic variability^9–13^, the characteristics of epigenomic conservation are still largely unclear. To address this issue, we studied the evolutionary conservation status of DNAm at the genome-wide level and performed a multiomics investigation to uncover its genetic associations, DNAm variability, and potential functional relevance. We found that MCCs can be characterized by their extensive DNAm stability in a wide range of healthy human tissues, DNAm dynamics in multiple cancers, and their roles in regulation of cancer driver genes. The characteristics of MCCs are similar to those of evolutionarily conserved genomic sequences, with involvement in maintaining normal development and function as well as alterations associated with disease. However, as 72.13% of MCCs were found outside of coding regions, characterizing MCCs provides important insight into the functional epigenetics underpinning regulatory evolution in contrast to genomic sequence comparisons between species, which mostly concentrate on coding sequences^75^. In addition, we found a positive relationship between DNAm conservation and genomic sequence conservation not only in CpGs but also in mQTLs. Although not all CpGs were associated with genetic variation, this partly explained the phylogenetic relationships among species that can be successfully captured by measuring DNAm levels in highly sequence-conserved cytosines, as observed in this and previous studies^20,29^. Our findings provide insight into epigenetic evolution and may help to elucidate the potential biological functions of DNAm conservation.

DNAm can provide information on evolutionary conservation that is complementary to underlying sequence data. We found that CpG sequence conservation showed a positive relationship with DNAm conservation, consistent with the previous literature on the genetic contexts (i.e., CpG and their neighboring nucleotides) underlying DNAm conservation^15–19^. Interestingly, the CpG sequence conservation at MCCs extended beyond the great ape lineage used to identify MCCs. This is reasonable as CpG sequence conservation is a prerequisite for DNAm conservation at CpG sites. More importantly, the mQTL sequence conservation was determined here to have a positive relationship with DNAm conservation. Considering the strong enrichment of mQTLs for MCCs in TF binding site-rich genomic features, such as promoters and enhancers, it is tempting to speculate that their relationship could be partially mediated by TF binding site turnover in epigenome conservation^16^. In addition to genomic sequence, the negative correlations of DNAm with other epigenetic marks, such as the histone modifications H3K4me2, H3K4me3, and H3K9me3, have been shown previously to be conserved across humans, mice, and pigs^9,18^. Although the mechanisms underlying DNAm evolution are still largely unclear, these observations suggest that DNAm can provide complementary evolutionary conservation information on CpGs over their underlying sequences alone by pointing toward CpG sequence conservation beyond the lineage tested for sequence and DNAm, linking with mQTL sequence conservation, and correlated with other epigenetic marks. As conservation analysis is one of the most widely used approaches for inferring these functionally important DNA sequences^76,77^, measuring DNAm conservation in addition to assessing genomic sequence conservation will help to complement elucidation of the functional capacity of genomic regions amenable to DNAm, particularly in noncoding regions, as 72.13% and 86.79% of MCCs and SCCs were outside of coding regions, respectively.

The information regarding DNAm conservation may serve as an important proxy for the functional importance of epigenetic variation. Evolutionarily conserved genomic information is a robust indicator of the functional importance of genetic variation and is widely leveraged for the biological interpretation of the findings of genome-wide association studies^28,78^. Our investigations provided evidence for the possible functional importance of MCCs over other CpGs. Specifically, by integration with transcription data, we identified an association of MCCs with expression of genes enriched in numerous crucial human biological pathways, including cell development, transcriptional regulation, and canonical cancer pathways. Therefore, understanding the conservation of DNAm could be beneficial in the functional interpretation of DNAm alterations in EWASs. Here, we leveraged an EWAS in atherosclerosis as an example and suggested that annotation of conservation information can improve our understanding of the potential functional roles of DNAm alterations. To facilitate the use of DNAm conservation information in epigenetics research, we developed QNMEC, a user-friendly web interface to systematically depict DNAm and sequence conservation information of CpGs at the genome-wide level.

MCCs have potential as an initial panel in cancer research. DNAm has been used as an early diagnostic and prognostic biomarker of various cancers^79,80^. Even if DNAm biomarkers perform well in terms of sensitivity and specificity, their application to clinical practice is still limited. One of the main limitations is the confounding effect of non-neoplastic factors (e.g., demographic characteristics) on DNAm patterns of biomarkers^81,82^. Therefore, the ideal DNAm-based cancer biomarker for clinical translation should exhibit strong associations with cancer biology and avoid unnecessary biases from non-neoplastic factors. In 5/34 cancer types, MCCs showed the largest DNAm changes between tumor and normal tissue samples compared to SCCs and NCCs due to their higher proportion and/or greater DNAm changes of cancer-specific CpGs. More importantly, in separate comparison to genes associated with SCCs and NCCs, genes whose expression was associated with MCC methylation showed stronger enrichment in cancer driver genes and a variety of canonical cancer pathways, including the Wnt signaling pathway and pathways involved in cell population proliferation, blood vessel development, and cell migration. These observations highlighted the potential roles of MCCs in cancer biology. In addition, MCC methylation is generally more stable than at other types of CpGs across common confounding factors, such as sex and ethnicity, in human populations, although these factors may be associated with the development of some cancers^83^. Given these benefits, MCCs represent a robust tool for the discovery of potential DNAm-based cancer biomarkers. Therefore, we advocate using MCCs as an initial panel to facilitate biomarker discovery studies and enable the development of clinical prediction models for cancer diagnosis and prognosis.

As we conducted separate analyses for each DNAm-based factor using different data sets, it was difficult to directly compare MCC methylation variability across factors. No methods are available that can completely remove batch effects across multiple data sets while retaining biological differences of interest. Therefore, the relative variability of MCCs in comparison to SCCs and NCCs associated with each factor was investigated indirectly. Although this study demonstrated the quantitative nature of MCCs in 42 different normal tissue types as well as the main peripheral blood and cord blood cell types, further research in other tissues and cell types is needed. While we successfully identified MCCs at the genome-wide level using the HM450K and EPIC platforms, identification was still limited as these microarrays cover at most ∼3.0% of human genomic CpG sites, and the true number of MCCs in great ape genomes will be substantially higher. Given that 2.56% of the CpGs tested here were MCCs, the T2T-CHM13 complete human genome assembly would be expected to contain roughly 0.8 million MCCs^84^ if the same percentage applies to the whole genome. Further studies using additional DNAm data sets with greater numbers of CpGs measured, larger sample sizes, and more diverse DNAm-based factors across all human tissues may be helpful to obtain a comprehensive understanding of the characteristics of MCCs.

## Conclusion

This study identified evolutionarily conserved CpGs in great apes and quantitatively characterized a set of CpGs with both sequence and methylation conservation (termed MCCs) in human populations. MCCs can be characterized as generally more stable across individuals than other CpGs and being less closely associated with demographic and environmental factors, tissue types, and noncancer diseases, while showing stronger associations with some cancer types. While correlated, DNAm conservation complemented sequence conservation of CpG sites. Functional enrichment analysis indicated that genes whose expression was associated with MCC methylation were enriched for cell development, transcriptional regulation, and canonical cancer pathways. Taken together, our findings provided insights into the genetic associations, DNAm variability, and potential functional relevance of evolutionarily conserved DNAm, laying a foundation for the application of conservation information in the functional interpretation of DNAm alterations in EWASs. As a resource for the epigenetics research community, we developed QNMEC, a user-friendly R Shiny web interface to illustrate evolutionary conservation and DNAm variability at the genome-wide level. This tool may accelerate the translation of epigenomic discoveries into the biological and clinical knowledge necessary to understand human disease.

## Methods

### Study subjects

To identify and characterize methylation-conserved CpGs (MCCs) in the context of as many methylation-related factors as possible, 202 DNAm array data sets with a total of 32,060 were retrieved, including in-house and publicly available data sets (Supplementary Table S1). Our study was performed using data from the Illumina Human Methylation 450 BeadChip (HM450K) and Illumina HumanMethylationEPIC BeadChip (EPIC) arrays. In addition, to explore MCC’s genetic association and potential functional relevance, 6 matched genotype data sets containing a total of 1058 samples and 13 matched transcription data sets containing a total of 3860 samples were included. Publicly available data sets were obtained from the NCBI Gene Expression Omnibus (GEO; https://www.ncbi.nlm.nih.gov/geo/), The Cancer Genome Atlas (TCGA; https://tcga-data.nci.nih.gov), the International Cancer Genome Consortium (ICGC; https://dcc.icgc.org/), and ArrayExpress databases (https://www.ebi.ac.uk/arrayexpress/)^85–88^. Detailed information for all data sets used in this study is provided in Supplementary Table S1.

Since most of the datasets used in this study were publicly available, four procedures were implemented to ensure data quality: for each tissue or group (case or control groups), data sets with sample sizes fewer than four were eliminated, multiple data sets for the same tissue were generally merged using the *ComBat* function in the sva R package^89^, independent data sets for the different tissues were used to validate the results, and additional quality control was performed after downloading the data set (see “DNA methylation array data preprocessing” subsection below).

### DNA methylation measurements

Regarding the in-house DNAm data, peripheral blood or buccal samples were collected from participants in cohorts (cohort descriptions are provided in Supplementary Data) and DNA was extracted from either the buffy coat, monocytes, or buccal swabs. 750 ng of purified genomic DNA was bisulfite converted using the EZ DNA Methylation Kit (Zymo Research) as per the manufacturer’s instructions. Specific incubation conditions for the Illumina Infinium Methylation Assay were used as per the manufacturer’s protocol Appendix. Bisulfite-converted DNA was assessed for concentration and quality using a NanoDrop^TM^ 8000 Spectrophotometer (Thermo Fisher Scientific), and 160 ng of the conversion product was used for genome-wide DNA methylation evaluation at over 485,000 CpG sites using the HM450K or over 850,000 CpG sites using the EPIC (Illumina, San Diego, CA, USA). Bisulfite-converted DNA from each sample was randomized across BeadChip arrays, as well as across sentrix rows, and run in one batch according to the manufacturer’s protocol.

### DNA methylation array data preprocessing

The in-house raw DNAm data were color corrected and background subtracted using GenomeStudio software. Data analysis was carried out using R version 3.5.1. Preprocessing was performed using the `minfi` and `methylumi` R packages^90,91^. Specifically, a subset of probes was removed for quality control purposes, including SNP control probes, probes within X and Y chromosomes, probes with a detection *P* > 0.05 in more than 1% of samples, probes with missing values in more than 5% of samples, probes with limited bead count (< 3) in more than 5% of samples, and polymorphic CpG probes^32^. Furthermore, samples with more than 1% of the probes missing were also excluded. To account for differences between probe types I and II, a beta-mixture quantile (BMIQ) normalization method was applied^92^. Known batch effects (chip and row) were corrected using the *ComBat* function in the `sva` R package^89^. For samples containing multiple cell types, cell proportions were estimated using the Houseman method and then regressed out from the DNAm data^50,93^. DNAm levels for each CpG site were presented as beta values, which were a ratio of the intensities of methylated bead signals over the sum of methylated and unmethylated bead signals.

The preprocessed and normalized data sets for the publicly available array samples used in this study were downloaded from GEO, TCGA, ICGC, and ArrayExpress databases^85–88^, and additional quality control was carried out. The same quality control processes as in-house DNAm data were conducted, with the exception of some processes that have been done (e.g., normalization) or cannot be performed due to the unavailable information for the publicly available data (e.g., some data sets with missing detection *P*-values).

### Identifying sequence conservation of CpGs

To capture CpGs that were comparable across humans and great apes, we mapped HM450K (*n* = 485,512) probes to chimpanzee (panTro5), bonobo (panPan2), gorilla (gorGor4), orangutan (ponAbe3), and human (hg19) reference genomes downloaded from the National Center for Biotechnology Information (NCBI). In addition to ensuring their mappability to reference genomes of great apes, we also uniquely excluded probes that could have multiple targets in these genomes (nonspecific probes), following the approach described previously^31–33^.

BLAT was used to map probe sequences to four different types of genomes for each reference genome: (1) an unmethylated bisulfite-treated genome with all Cs converted to Ts; (2) a methylated bisulfite-treated genome with only non-CpG Cs converted to Ts; (3) the reverse-complement of (1); and (4) the reverse-complement of (2)^94^. BLAT was implemented using the following parameters: tileSize = 11, stepSize = 5, and repMatch = 1,000,000. For a probe to be considered sequence conserved and nonspecific, the sequence alignment had to have at least 90% identity, at least 40 of 50 matching bases, no gaps, the CpG locus had to be perfectly matched, with only one specific hit aligning to the genome. Following selection, 435,255 human, 343,891 chimpanzee, 348,713 bonobo, 325,794 gorilla, and 264,329 orangutan probes were retained in HM450K. As HM450K were designed based on the human reference genome, the numbers of CpGs kept in this step were consistent with the evolutionary distance of these species to human^34,35^. Ultimately, we defined only the 169,500 HM450K probes that were successfully aligned to all great ape genomes as sequence-conserved. The additional EPIC (*n* = 413,406) probes that were not covered by the HM450K array were also mapped to great ape reference genomes, and the 121,547 additional probes shared by these species were retained.

### Identifying methylation conservation of CpGs

To identify methylation-conserved CpGs (MCCs), the Kruskal–Wallis rank-sum test was used to compare DNAm patterns among species. *P*-values were corrected for multiple testing using the Benjamini–Hochberg method^95^. Furthermore, DNAm changes for each CpG were measured between each pairwise combination of species. CpGs that showed highly stable methylation patterns among great apes with FDR > 0.1 and between-species methylation differences (absolute beta value changes of any two species) < 0.05 were defined as MCCs.

### Reconstructing a phylogenetic tree based on DNAm information

A phylogenetic tree of great apes was reconstructed based on the DNAm of 159,502 selected CpG probes from HM450K after elimination of 9998 probes from the total of 169,500 as a consequence of DNAm data quality control (Fig. 1a). DNAm levels were discretized into five fraction states using the default method described in the `PhyloEpiGenomics` R package^22^. We reconstructed a maximum likelihood tree based on a Markov model of the evolution of DNAm fractions^22^. We used the “No Jump” evolutionary model that assumed a methylation fraction status could not change into a nonadjacent state during the course of a short period of time^22^.

### Pathway enrichment analysis of genes

We mapped genes for MCCs to biological pathways to investigate their functional significance. The analysis was restricted to CpGs from promoter and gene body regions. Gene Ontology (GO) molecular functions, GO biological processes, GO cellular components, and the Kyoto encyclopedia of genes and genomes (KEGG) pathway analyses were performed using the g:Profiler toolset (https://biit.cs.ut.ee/gprofiler/gost)^96^. Genes for SCCs and NCCs were treated separately as background. Pathways were deemed to be significantly enriched at a threshold of Benjamini–Hochberg FDR < 0.05.

### Enrichment analysis of MCCs and mQTLs for known genomic elements

To calculate enrichment statistics for MCCs presented in a specific genomic element, the same numbers of SCCs and NCCs were selected separately at random. For each genomic element, the fold enrichment was determined by comparing the number of these MCCs that were mapped to the genomic element with the expected number estimated through 1000 resamplings of SCCs and NCCs separately. Statistical significance was defined as FDR < 0.05 after multiple test correction of the empirical *P*-value using the Benjamini-Hochberg method^95^. The UCSC Genome Browser (https://genome.ucsc.edu/) was used to retrieve annotation files for the coding exon, intron, promoter (1 kb), 1 kb downstream, 5′-UTR, 3′-UTR, and CpG island regions. CpG shores were derived from CpG island coordinates by taking 2-kb flanking regions, and CpG shelves were obtained by taking 2-kb flanking regions outward from shores. Annotations for long noncoding RNAs (lncRNAs) and pseudogenes were obtained from the GENCODE database (https://www.gencodegenes.org/releases/19.html)^97^. The microRNA (miRNA) coordinates were downloaded from FANTOM5 (http://slidebase.binf.ku.dk/human_mirna/)^98^. Super-enhancer coordinates were obtained from the dbSUPER database (http://bioinfo.au.tsinghua.edu.cn/dbsuper/)^99^. Enhancer data were obtained from ENCODE (http://ftp.ebi.ac.uk/pub/databases/ensembl/encode/integration_data_jan2011/byDataType/segme ntations/jan2011/Combined_7_state/), FANTOM5 (http://slidebase.binf.ku.dk/human_enhancers/), and RoadMap (http://egg2.wustl.edu/roadmap/data/byDataType/dnase/BED_files_enh/)^98,100,101^. All genomic coordinates in this study were based on the GRCh37/hg19 human reference genome.

Similarly, for analysis of genomic element enrichment of methylation quantitative trait loci (mQTLs) for MCCs, 1000 permutations were applied to mQTLs for SCCs and NCCs, separately. We calculated the fold enrichment of these mQTLs in each genomic feature in comparison to randomly selected mQTLs for SCCs or NCCs. Enrichments were considered statistically significant at a threshold of Benjamini-Hochberg FDR < 0.05.

### Evolutionary conservation score calculation

To estimate the sequence conservation of CpG sites, we used two sets of PhastCons scores separately based on the alignment of primate genomes and great ape genomes^40^. The PhastCons scores in primates had been estimated based on multiple sequence alignments of 10 primate genomes: human (*H. sapiens*), chimpanzee (*P. troglodytes*), gorilla (*G. gorilla gorilla*), orangutan (*P. pygmaeus abelii*), rhesus macaque (*M. mulatta*), baboon (*P. hamadryas*), marmoset (*C. jacchus*), tarsier (*T. syrichta*), mouse lemur (*M. murinus*), bushbaby (*O. garnettii*)^41^. The file containing PhastCons scores for multiple alignments of nine nonhuman primate genomes to the human genome at single-base resolution was downloaded from the UCSC Genome Browser at https://hgdownload.cse.ucsc.edu/goldenPath/hg19/phastCons46way/primates/ ^41^. The PhastCons scores of CpG sites were extracted from this download file based on their genomic coordinates. PhastCons scores of MCCs were compared to those of other CpG types using the 1000 permutation test. Statistical significance was defined as FDR < 0.05 after multiple test correction of the empirical *P*-value using the Benjamini-Hochberg method^95^.

In addition, we calculated PhastCons scores in great ape genomes with the alignment of gorilla, chimpanzee, and human genomes. Briefly, we first downloaded a pairwise human–ape alignment from https://github.com/mcfrith/last-genome-alignments^44^. In pairwise alignment, each great ape genome had been aligned to the human genome via the “lastal” command in LAST version 861 with the parameters “-m50 -E0.05 -C2” and then further processed using the “maf-swap” command to change the sequence order in alignments^44^. Next, using MULTIZ version 11.2, we combined the pairwise alignments into multiple genome alignments with the human genome as a reference^102^. Ultimately, we obtained PhastCons scores for each base of the human genome using the PHAST program (version 1.5) with the commands: “phastCons – target-coverage 0.25 –expected-length 12 –rho 0.4”^40,103^. As we did with PhastCons scores in primates, we extracted PhastCons scores of CpG sites in great apes and conducted comparisons of PhastCons scores of MCCs to other CpG types.

### Measurement of DNA methylation variability across individuals and DNA methylation differences between phenotypic groups

Two metrics, DNAm difference and DNAm variability, were used to assess the quantitative nature of methylation for each CpG type. Subtraction was performed to calculate methylation delta beta values for CpGs between groups, and the absolute delta beta values were defined as the DNAm differences. As some variables are continuous, such as age and body mass index, a median value or suggested cutoff point was used to assign them into two groups (Supplementary Table S15). For example, 25 kg/m² was used here as a body mass index cutoff between healthy weight and overweight as proposed by the World Health Organization^104^. The commonly used DNAm change threshold of 0.03 for EWASs, which tends to be above the technical noise of Illumina arrays, was used^57^. The SD was used to calculate DNAm variability, which reflects methylation change at a specific CpG site across individuals. We did not set a fixed numerical threshold for DNAm variability because the SD is affected by sample size. As DNAm beta value is associated with SD^46^, we repeated the investigations of DNAm variability by converting beta values to M values in all human tissues tested to avoid the potential uneven variability across the DNAm range^37^.

### Comparison of DNA methylation variability

A permutation test was used to compare the average methylation variability among MCCs, SCCs, and NCCs. Unless otherwise specified, beta values are utilized for analysis since they are easier to interpret from a biological perspective than M values^37^. The labels of the values in the two groups for three comparisons (MCCs vs. SCCs, MCCs vs. NCCs, and SCCs vs. NCCs) were randomly shuffled. The randomization procedures were repeated 1000 times for SCCs (for MCCs vs. SCCs) and NCCs (for MCCs vs. NCCs and SCCs vs. NCCs) separately. Comparisons were performed between two groups by a one-tailed test, and changes in DNAm variability were considered statistically significant at a threshold of Benjamini–Hochberg FDR < 0.05.

### Genotype data preprocessing

Human genotype data from six common tissues in which MCC variability was tested in the above analysis were used in this study to examine the association between CpG methylation and genotype (Supplementary Table S1). These in-house genotype data sets included 504 blood samples measured with the Infinium Global Screening Array-24 v1.0 BeadChip and 279 buccal swab samples measured with the Infinium PsychArray-24 v1.2 BeadChip. Further details of genotype profiling are provided in Supplementary Data. The publicly available genotype data sets included 78 brain samples measured with the Illumina PsychArray-24 v1.1 BeadChip (GSE112525), 89 LCL samples measured with the Illumina HumanHap650Yv3 Genotyping BeadChip (GSE24277), 48 saliva samples measured with the Illumina HumanHap550-Duov3 Genotyping BeadChip, HumanOmniExpress-12 v1.0 BeadChip, and Illumna HumanOmniExpressExome-8 v1.2 BeadChip (GSE99091), and 60 skin samples measured with the Human1M-Duo v3.0 DNA Analysis BeadChip (GSE53261). For each genotype data set, individuals with discordant sex information, elevated outlying heterozygosity rate or missing genotype rate, high relatedness, and distinct population stratification were checked and removed using PLINK v1.90b4 ^51^. SNPs with a minor allele frequency (MAF) < 0.01, missing genotype rate > 10%, or Hardy–Weinberg equilibrium exact test *P* < 1.0 × 10^−6^ were also excluded. The Michigan Imputation Server was used for genotype imputation, and the reference panel 1000 Genomes Project Phase 3 or haplotype reference consortium version 1.1 was used^105–107^. We estimated the haplotype phase using Eagle version 2.4 and Minimac4 to impute the missing SNPs^105,108^. Each tissue sample was imputed separately. Imputed SNPs with MAF < 0.05, missing genotype rate > 10%, or Hardy–Weinberg equilibrium exact test *P* < 1.0 × 10^−6^ were excluded. Finally, 5,556,332 SNPs in blood, 5,412,051 in brain, 5,498,067 in buccal swabs, 9,530,480 in LCLs, 8,088,158 in saliva, and 5,479,474 in skin were retained for further analysis.

### Mapping of mQTLs

We scanned for human mQTLs by testing SNPs located within ± 10-kb windows of each CpG, consistent with previous studies indicating that *cis*-mQTLs are usually close to their corresponding SNPs^43^. A linear additive model adjusted for age and sex was applied to compute the associations between CpG methylation and neighboring SNPs using the `MatrixEQTL` R package^109^. mQTL-CpGs were defined in each of the six tissues separately. The FWER was calculated by investigating the association between CpG methylation and genotypes in 100 randomly sampled sets of DNAm levels, and recomputing the *P*-values for associations for each permutation^110^. After each permutation, we kept the most significant *P*-value per CpG-SNP pair and determined the *P*-value thresholds that provided a 5% FWER in the six tissues: 5.01 × 10^−4^ in blood; 4.25 × 10^−4^ in brain; 4.42 × 10^−4^ in buccal swabs; 2.23 × 10^−4^ in LCLs; 3.07 × 10^−4^ in saliva, and 2.70 × 10^−4^ in skin.

### Identification of cancer-specific CpGs

Four gastrointestinal cancers (COAD, ESCA, PAAD, and SBA) were used as examples in this analysis. There were 294, 180, 177, and 28 tumors and 37, 16, 10, and 28 normal tissues, separately. These normal samples were healthy tissues adjacent to tumors. To identify cancer-specific CpGs in each cancer type, we used the Mann–Whitney U test to measure the DNAm differences between tumor and normal tissues. FDR values were computed using the Benjamini– Hochberg method^95^. CpGs with DNAm difference > 0.03 and FDR < 0.05 were considered statistically significant. The DNAm difference cutoff of 0.03 is a commonly used threshold for EWASs and tends to be above the technical noise level of Illumina arrays^57^.

### Association of DNAm with gene expression

To identify genes whose expression is associated with MCC methylation, the TCGA level 3 gene expression data of 17 cancer types that were used in the above analyses were downloaded from UCSC Xena data hubs (https://xenabrowser.net/datapages/)^111^. The Illumina HiSeq 2000 RNA sequencing platform was used by the University of North Carolina TCGA genome characterization lab to generate the gene expression data. Gene-level transcription in the expression data had been estimated as the log2(x+1) transformed RSEM normalized count. After matching the individuals with the DNAm data, 13 cancer types with sample sizes ranging from 140 to 448, including four gastrointestinal cancers and nine gastrointestinal cancers, were retained for subsequent analyses (Supplementary Table S1). CHOL (*n* = 24), LUAD (*n* = 15), LUSC (*n* = 33), and READ (*n* = 74) were excluded due to their limited sample sizes. Clinical data for these samples were also downloaded from the UCSC Xena data hubs.

We associated DNAm at 320,740 CpG sites (9077 for MCCs, 110,877 for SCCs, and 200,786 for NCCs) within the promoters and gene bodies of 24,144 genes to the levels of expression in each cancer type. We found CpGs whose levels of DNAm (M values) were associated with gene expression using linear regression with adjustment for age, sex, and disease status (i.e., tumor or normal tissue). FDR values were calculated using the Benjamini–Hochberg method^95^. Associations were considered significant at a threshold of FDR < 0.05.

## Supporting information

Supplementary Results

## Abbreviations

5mC: 5-methylcytosine
BLCA: bladder urothelial carcinoma
BMIQ: beta-mixture quantile normalization method
BRCA: breast invasive carcinoma
CIMP: CpG island methylator phenotype
COAD: colon adenocarcinoma
CpG: cytosine-phosphate-guanine dinucleotide
CSC: cancer stem cell
DNAm: DNA methylation
DZ: dizygotic
EPIC: Illumina HumanMethylationEPIC BeadChip
ESCA: esophageal carcinoma
EWAS: epigenome-wide association study
FDR: false discovery rate
GEO: NCBI Gene Expression Omnibus
HM450K: Illumina Human Methylation 450 BeadChip
HNSC: head and neck squamous cell carcinoma
ICGC: International Cancer Genome Consortium
IL-13: interleukin-13
IL-17: interleukin-17
KIRC: kidney renal clear cell carcinoma
KIRP: kidney renal papillary cell carcinoma
LCL: lymphoblastoid cell line
LIHC: liver hepatocellular carcinoma
lncRNAs: long noncoding RNAs
LUAD: lung adenocarcinoma
MAF: minor allele frequency
MCCs: methylation-conserved CpGs
miRNAs: microRNAs
mQTLs: methylation quantitative trait loci
MZ: monozygotic
NCBI: National Center for Biotechnology Information
NCCs: nonconserved CpGs
PAAD: pancreatic adenocarcinoma
PRAD: prostate adenocarcinoma
READ: rectum adenocarcinoma
SARC: sarcoma
SBA: small bowel adenocarcinoma
SCCs: sequence-conserved CpGs
SD: standard deviation
SNP: single-nucleotide polymorphism
TCGA: The Cancer Genome Atlas
TF: transcription factor
THCA: thyroid carcinoma
UCEC: uterine corpus endometrial carcinoma

## Acknowledgements

We are grateful to Compute Canada’s National Systems for Scientific Computation for computational support. We are extremely grateful to our colleague, Alan Kerr, whose wonderful contribution to writing and style greatly improved the manuscript. We are grateful to the individuals who participated in the Costa Rican Longevity and Healthy Aging Study (CRELES) study for providing their personal information and biological specimens without payment or compensation, as well as to the Costa Rican institutions (CONAPAM, CCSS, INEC, TSE, and INISA) that collaborated in data and specimen collection. We are also grateful to the staff and fieldworkers at the Central American Population Center (CCP) of the University of Costa Rica (UCR) who made the CRELES study possible.

## Author contributions

ZD conceived, designed, and performed the computational analyses, analyzed the data, and interpreted the results. JLM, DHR, WTB, LRB, LQM, EP, GEM, and NRB contributed DNA methylation and genotype data. SS, MF, JW, and KK contributed ideas and participated in evaluating results and discussions. MSK supervised the study and obtained funding. ZD wrote the manuscript, with input from all authors. All authors approved the final version of the manuscript.

## Data availability

All data sets used in this study are listed in Supplementary Table S1. The publicly available genotype and methylation array data can be accessed in the NCBI Gene Expression Omnibus (GEO; https://www.ncbi.nlm.nih.gov/geo/), The Cancer Genome Atlas (TCGA; https://tcga-data.nci.nih.gov), the International Cancer Genome Consortium (ICGC; https://dcc.icgc.org/), and ArrayExpress databases (https://www.ebi.ac.uk/arrayexpress/). The CRELES DNA methylation and genotype data are not publicly available; requests for restricted access to this data can be submitted at http://www.creles.berkeley.edu/ following institutional review approval. The Expression Collaborative for Kids Only (GECKO), CareGiver (CG), and EvoImmunoPop DNA methylation data utilized in this study have been deposited in the GEO under accession code <u>GSE124366</u> ^112^, <u>GSE137903</u> ^113^, GSE281200, and GSE281199. Summary statistics are available in https://epigenome.shinyapps.io/QNMEC/.

## Code availability

The scripts used to generate the figures and support the findings of this study are available in https://github.com/functionalepigenomics/Methylation-conserved-CpGs.

## Ethics approval and consent to participate

These in-house data sets were approved by the institutional review board policies at the University of British Columbia and C&W Research Ethics Board (UBC Ethics #: H15-02143 for the CRELES data, H16-02224 for the EVOIMMUNOPOP data, H07-02773 for the GECKO data, and H12-03101 for the CG data). All participants provided written informed consent.

