## Supplementary Data for "Genetic Basis, Quantitative Nature, and Functional Relevance of Evolutionarily Conserved DNA Methylation"

**Costa Rican Longevity and Healthy Aging Study (CRELES) cohort**

The CRELES (http://creles.berkeley.edu) is a prospective, longitudinal study involving a nationally representative cohort of 2,827 Costa Rican residents aged 60 and older at baseline (2004–2006)^1^. A second wave of interviews and data collection took place between 2006 and 2008. Additionally, a complementary sample of Nicoyan quasi-centenarians (aged 95 and older) was included. Data collection, physical examinations, and specimen collection were conducted in participants’ homes. Detailed information on the study’s sampling, field procedures, and laboratory protocols has been previously published^2^. The Ethical Science Committee of the University of Costa Rica granted approval (VI-763-CEC-23-04), and all participants provided written informed consent in accordance with the Declaration of Helsinki. The DNA samples used in this study, which are owned by the UCR, were transferred to the University of California Berkeley under a Non Commercial Material Transfer Agreement and Notification of Transfer on July 3, 2013.

For the present work, we randomly selected 512 individual samples from blood collected during waves 1 and 2. Whole blood samples were collected via venipuncture and processed at the University of Costa Rica, as previously described^3,4^. Genomic DNA was extracted from 2 mL of frozen whole blood using the phenol-chloroform method. DNA methylation (DNAm) data were generated using the Illumina HumanMethylationEPIC BeadChip (EPIC) array (Illumina, San Diego, CA, USA). Genotyping data was measured at 618,540 single nucleotide polymorphism (SNP) sites using the Infinium Global Screening Array (GSA) BeadChips according to the Illumina’s standard protocol (Illumina, CA, USA). GenomeStudio 2.0 Genotyping software was used to transform the raw intensity files into clusters, and subsequently make genotype calls by producing cohort-specific clustering files and manifest GSA-24v1-0_C1 (Version 1 A2, Illumina).

**Gene Expression Collaborative Kids Only (GECKO) cohort**

The GECKO cohort is a prospective Canadian cohort of 402 families with 6–11-year-old children designed to examine how social environment relates to child development and gene regulation^5^. Mothers were recruited from Vancouver, British Columbia, and provided informed consent for collection of survey data for themselves as well as buccal DNA samples from their children (age range 6–11 years, *n* = 376). This study was approved by the Clinical Research Ethics Board at the University of British Columbia.

For the present work, 345 buccal samples were obtained from children using Isohelix Buccal Swabs (Cell Projects Ltd., Kent, UK) and Genomic DNA from stabilized buccal samples was isolated using Isohelix Buccal DNA Isolation Kits (Cell Projects Ltd., Kent, UK) and was purified and concentrated using DNA Clean & Concentrator (Zymo Research, CA, USA). DNA extracted from buccal samples was applied to the Illumina Infinium HumanMethylation450K (HM450K) BeadChip array (Illumina, San Diego, CA, USA). Saliva samples were also collected from 279 children to extract DNA using the Oragene OG-500 DNA all-in-one system, following the manufacturer’s protocol (DNA Genotek Inc., ON, Canada). Genotyping data was measured at 588,454 SNP sites using the Illumina Infinium PsychChip BeadChip (PsychChip), as per manufacturer’s protocols (Illumina, CA, USA). All experimental procedures were conducted in accordance with institutional review board policies at the University of British Columbia. Written informed consent was obtained from a parent or legal guardian and assent was obtained from each participant.

**EVOIMMUNOPOP dataset**

This dataset consisted of 171 individuals living in Belgium, from the EVOIMMUNOPOP project^6^. For each participant, 300 ml of whole blood was collected into anticoagulant EDTA-blood collection tubes and peripheral blood mononuclear cells (PBMCs) were purified using Ficoll-paque density gradients (#17-1440-03, GE Healthcare). Monocytes were positively selected from purified PBMCs using magnetic CD14 microbeads (#130-050-201, MiltenyiBiotec), according to the manufacturer’s instructions. All samples had a monocyte purity higher than 90%, with a mean value of 97%. Genomic DNA from each sample was isolated from the monocyte fraction using the phenol/chloroform protocol, followed by ethanol precipitation. Quantitative DNAm measurements were performed using the HM450K array (Illumina, San Diego, CA, USA), following the manufacturer’s experimental procedures. Four technical replicates (two in each population) were applied during HM450K processing. Note that 156 individuals in this dataset were also analyzed using the EPIC array, and the array data were deposited in GEO: GSE120610^6^. All healthy donors provided informed consent. All experiments were approved by the Ethics Board of Institut Pasteur (EVOIMMUNOPOP-281297) and the relevant French authorities (CPP, CCITRS and CNIL), in accordance with applicable laws, regulations, and ethical principles consistent with the Declaration of Helsinki.

**CareGiver (CG) cohort**

The CG participates were recruited from the central nervous system tumor clinics at the British Columbia Cancer Agency, Vancouver Centre, as previously described^7^. Caregivers of patients with glioblastoma who attended clinic prior to the onset of radiotherapy were approached for participation. Controls were recruited from the broader Vancouver community using advertisements in local media. They had to be without major stressors in their lives during the past year, including divorce, bereavement, unemployment, victimization, significant illness, hospitalization, or care giving responsibilities of their own. Collection of all human samples used in this study was approved by the joint University of British Columbia and Children and Women’s Hospital Ethics board (Certificate: H08-02773), and all subjects gave written consent.

For the present work, 88 samples from healthy participants in the CG cohort were used. Whole blood was collected for these participants, and PBMCs were isolated. Monocytes, defined as CD14^+^, were separated using immune-magnetic capturing methods (Miltenyi AutoMACS) as previously reported. DNA was isolated from the purified monocytes using a column-based method and assessed for quality using NanoDrop, bisulfite converted using EZ-96 DNA Methylation kits (Zymo Research, Irvine, CA) and run on the HM450K array (Illumina, San Diego, CA, USA).
