## Supplementary Results for "Genetic Basis, Quantitative Nature, and Functional Relevance of Evolutionarily Conserved DNA Methylation"

**Cross-tissue validation of MCC methylation conservation**

We further validated MCCs in these data sets with M values (calculated as the log2 ratio of the intensities of methylated probes versus unmethylated probes) to avoid the potential uneven variability of beta values, as the DNAm range of beta values can result in extremes having considerably lower variability^1^. Smaller average values of DNAm differences between chimpanzee and human samples were observed in MCCs rather than SCCs in blood (0.447 vs. 0.654, respectively, *P* < 0.001, 1000 permutation test), prefrontal cortex (0.442 vs. 0.569, respectively, *P* < 0.001, 1000 permutation test), lateral cerebellum (0.482 vs. 0.603, *P* < 0.001, 1000 permutation test), and dermal fibroblasts (0.395 vs. 0.475, respectively, *P* < 0.001, 1000 permutation test) (Supplementary Fig. S1b).

**Extensive DNAm stability of MCCs in a wide range of healthy human tissues**

One major concern about these findings was whether the relative stability of MCCs could be explained by fewer intermediately methylated CpGs in MCCs compared to SCCs and NCCs separately, as previous studies reported a link between the intermediately methylated status and large interindividual DNAm variability^1,2^. To test this, we converted beta values to M values to avoid the potential uneven variability across the DNAm range^1^ and repeated the comparison of DNAm variability among CpG types in all 42 human tissues as well as peripheral blood and cord blood cell types. Consistent with the initial analysis, we found the highest stability of MCCs relative to SCCs and NCCs in all tissues and cell types tested, with 0.79%–43.49% and 1.04%–46.02% (depending on tissues and cell types) greater stability on average compared to SCCs and NCCs, respectively (all Benjamini–Hochberg FDR < 0.05, 1000 permutation test) (Supplementary Fig. S8 and 9).

**Characteristics of the association between genetic variation and MCC methylation**

Further validations were conducted by converting beta values to M values to reduce the possible cofounding effect of uneven DNAm variability across the DNAm range of DNAm beta values in investigating associations between CpGs and SNPs^1^. We found that in all six tissues MCCs had 10.60%–40.94% and 31.84%–46.30% fewer and 5.47%–33.25% and 11.44%–30.19% weaker associations with SNPs in comparison to SCCs and NCCs, respectively, across tissues examined (all Benjamini–Hochberg FDR < 0.05, with the exception of the comparison of MCCs to SCCs in skin) (Supplementary Fig. S12).

**MCCs are less likely to be associated with tissue specificity than SCCs and NCCs**

To exclude the possible cofounding effect of uneven DNAm variability across the DNAm range of beta values^1^, we repeated these analyses after converting beta values to M values and obtained similar results: MCCs were 7.59%–41.90% and 2.75%–41.36% more stable on average compared to SCCs and NCCs, respectively (all Benjamini–Hochberg FDR = 1.5 × 10^−3^, 1000 permutation test) (Supplementary Fig. S15). Meanwhile, SCCs showed the largest DNAm variability among tissues and cell types, with 0.56%–7.95% more variability on average over NCCs (all Benjamini–Hochberg FDR = 1.5 × 10^−3^, 1000 permutation test). These results further confirmed the findings outlined above.

**MCCs are more likely to be associated with certain cancers than SCCs and NCCs separately, but not with demographic and environmental factors or noncancer diseases**

To avoid the potential confounding effect of uneven variability across the DNAm range of beta values^1^, we converted them to M values and repeated the comparison of the DNAm differences among CpG types between phenotypic groups of each factor or disease. Consistent with the above findings, using an M value difference threshold of 0.4 as previously suggested^1^, the average DNAm differences for MCCs, SCCs, and NCCs were less than this threshold in all 49 demographic and environmental factors and almost all noncancer diseases examined (68/69 for MCCs, 65/69 for SCCs, and 65/69 for NCCs), but larger in the majority of cancers (28/34 for MCCs, 30/34 for SCCs, and 30/34 for NCCs) (Supplementary Fig. S18 and 19). We also showed that MCCs were more stable than SCCs and NCCs in 39 out of 49 demographic and environmental factors using a Benjamini–Hochberg FDR threshold of 0.05 via the 1000 permutation test, with 2.85%–36.73% and 1.48%–35.22% smaller average methylation differences, respectively. Of 69 noncancer diseases examined, 55 showed the most stability with 1.92%–33.45% and 1.64%–33.33% smaller average DNAm differences over SCCs and NCCs, respectively, using a Benjamini–Hochberg FDR threshold of 0.05 via the 1000 permutation test, but only 12 of 34 cancers showed the most stability with 5.16%–30.55% and 4.18%–23.07% smaller average methylation differences over SCCs and NCCs, respectively, (Supplementary Fig. S18 and 19). That is, MCCs exhibited similar or greater methylation differences than SCCs and/or NCCs in 22/34 cancer types. Notably, eight cancers showed the largest average DNAm difference in MCCs, including all five cancers tested with beta values. These findings generally supported our results with beta values.
