## Supplementary Figures for "Genetic Basis, Quantitative Nature, and Functional Relevance of Evolutionarily Conserved DNA Methylation"

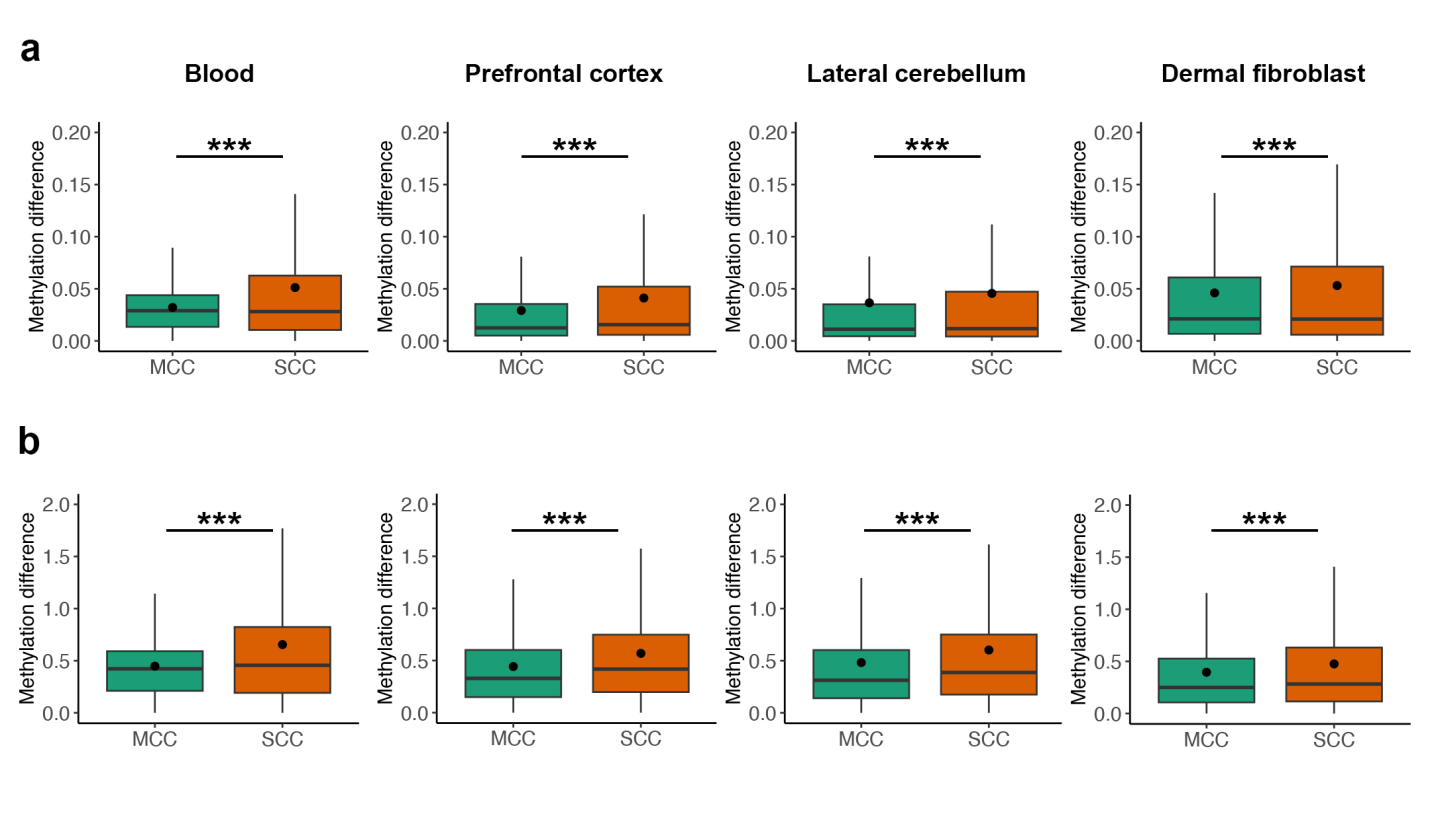

**Figure S1.** Validation of MCC methylation conservation in multiple tissues between chimpanzees (*n* = 14, 8, 8, and 7 in whole blood, prefrontal cortex, lateral cerebellum, and dermal fibroblasts, respectively) and humans (*n* = 20, 7, 7, and 2 in whole blood, prefrontal cortex, lateral cerebellum and dermal fibroblasts, respectively). **(a)** Comparison based on beta values. *** *P* < 0.001. Mean values are presented as circles in each box plot. Outliers are not shown in box plots. **(b)** Comparison based on beta-converted M values.

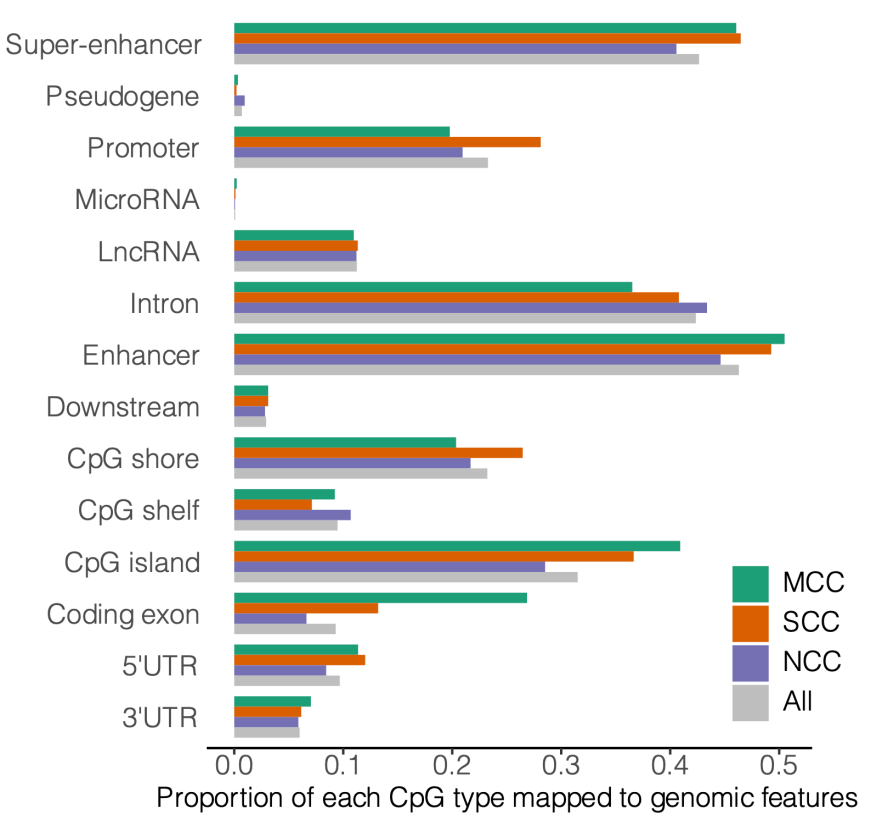

**Figure S2.** Colocalization of MCCs (*n* = 11,500) with 14 known genomic features.

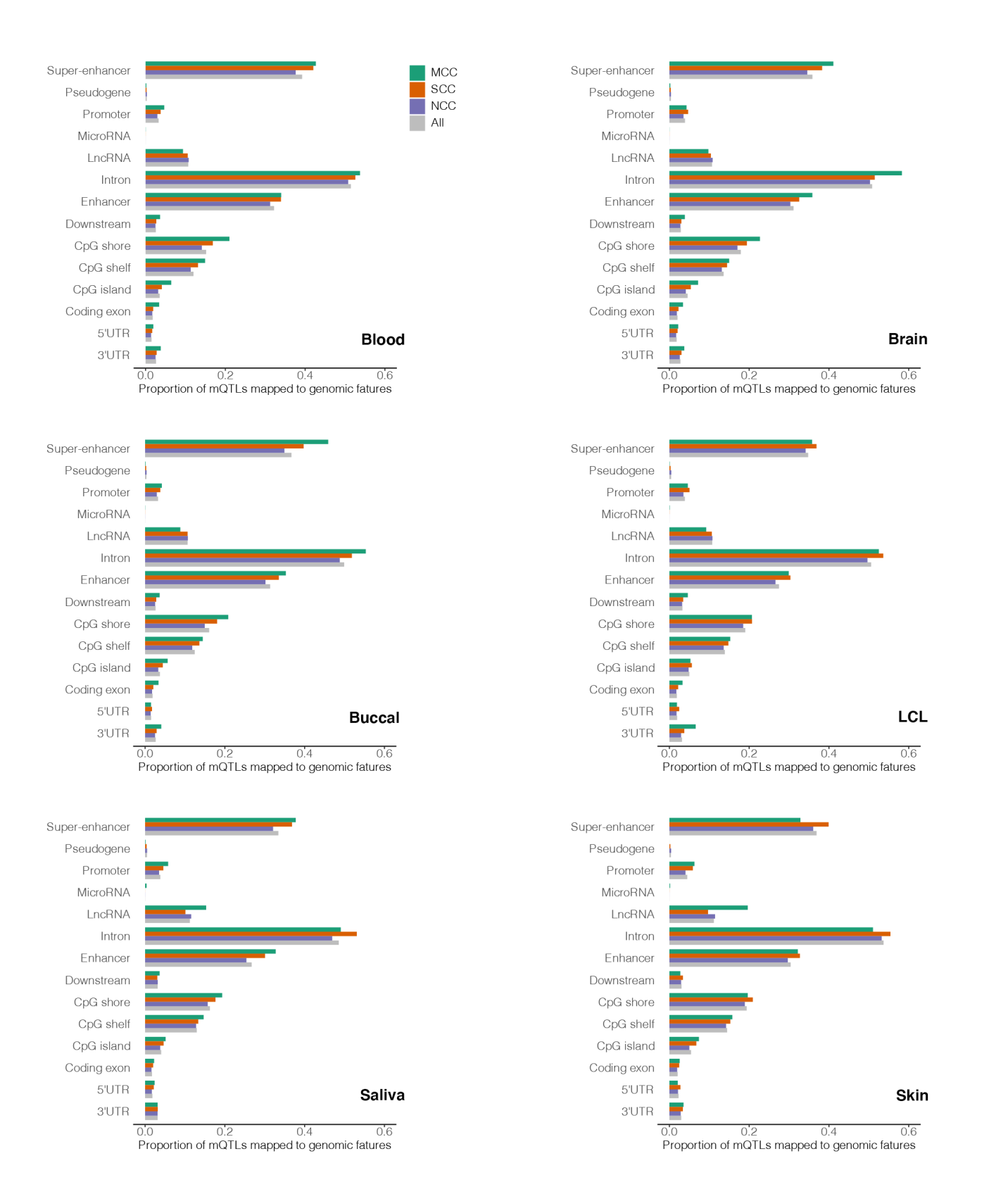

**Figure S3.** Colocalization of mQTLs for MCCs with 14 known genomic features. Abbreviation: LCL, lymphoblastoid cell line.

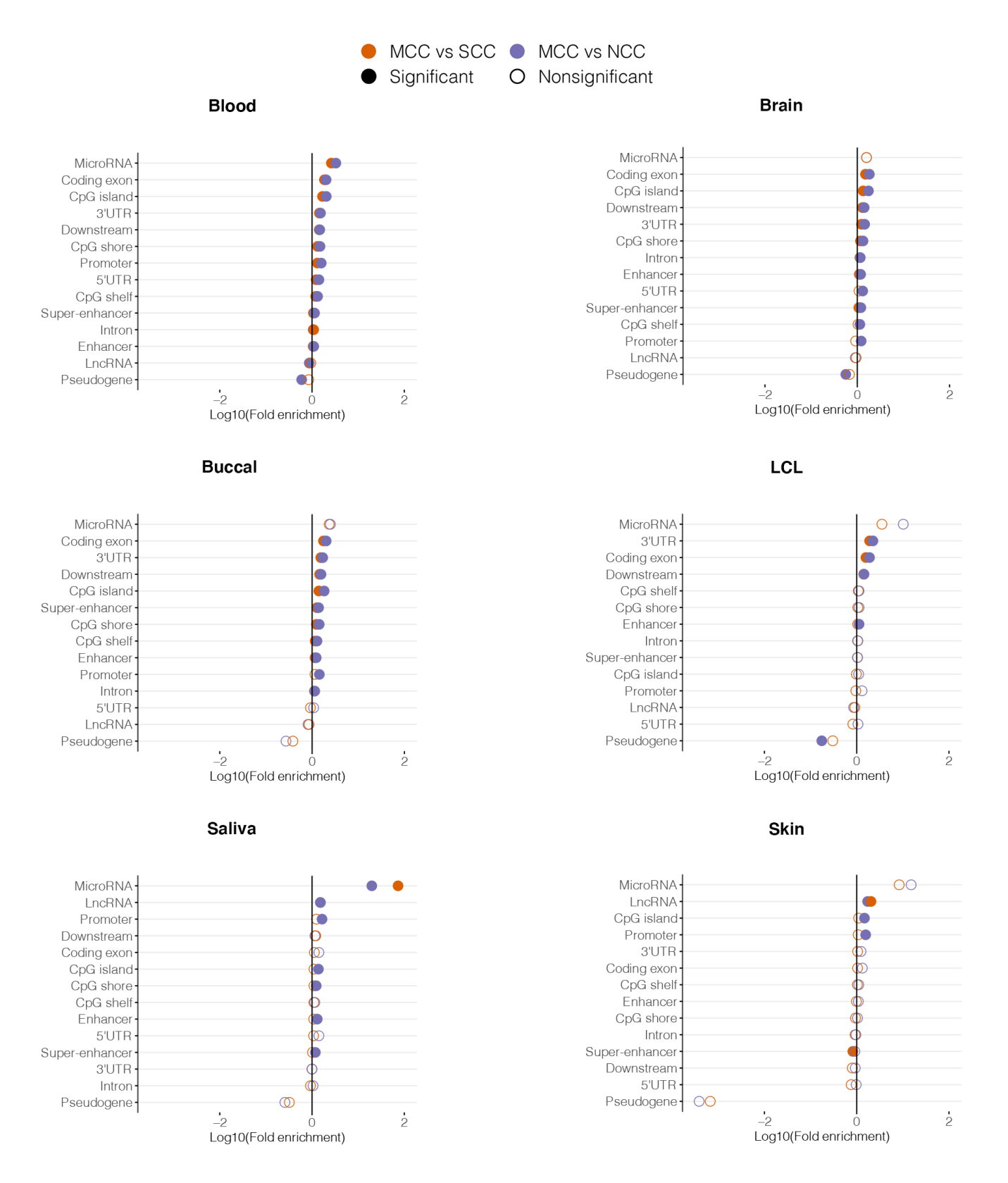

**Figure S4.** Significant enrichment and depletion of mQTLs for MCCs in genomic features compared separately to 1000 randomizations of mQTLs for SCCs and NCCs. Features with enrichment FDR < 0.05 are considered significant. Abbreviation: LCL, lymphoblastoid cell line.

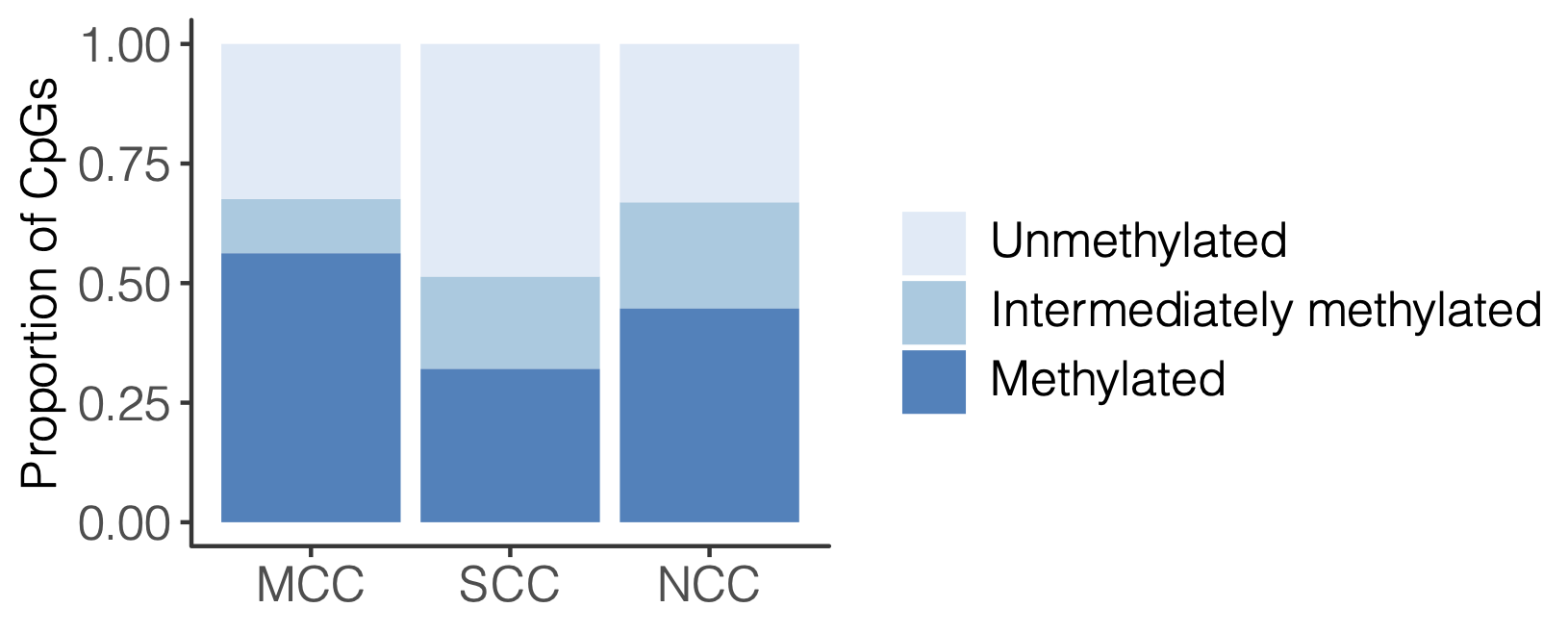

**Figure S5.** Proportions of MCCs with methylated (beta > 0.75), unmethylated (beta < 0.25), and intermediately methylated (0.25 < beta < 0.75) patterns in human blood samples separately compared to SCCs and NCCs. The figure represents the proportion of MCCs identified using the threshold of FDR > 0.1.

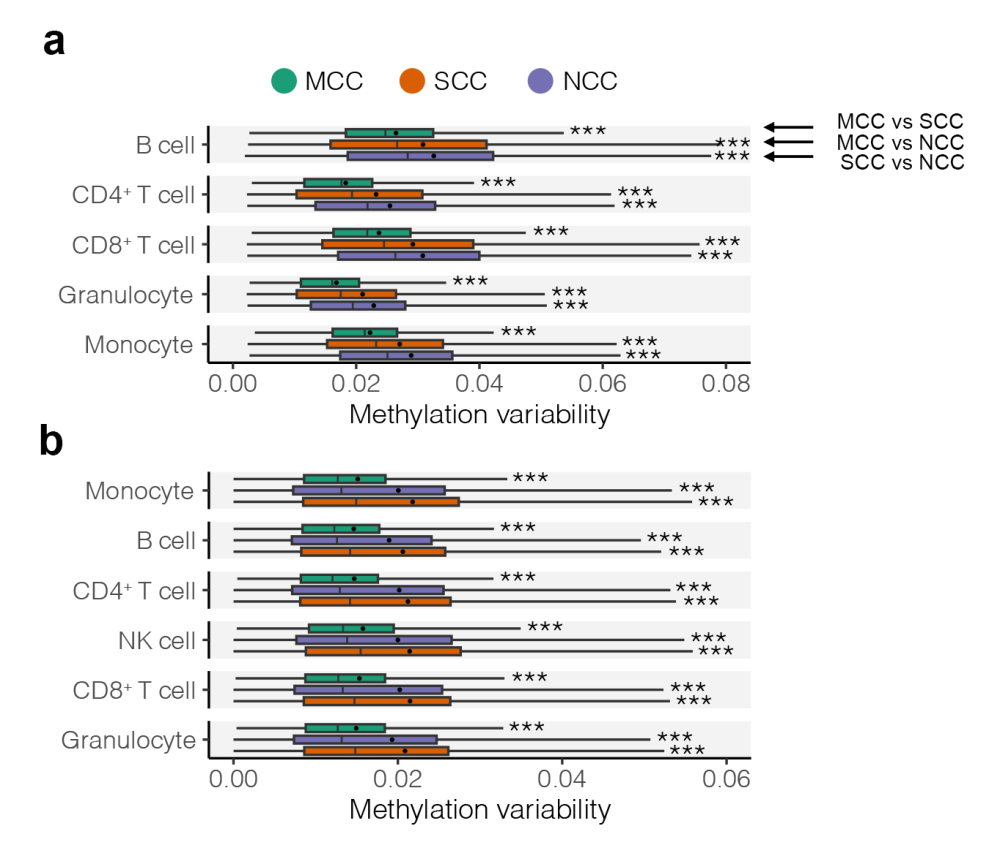

**Figure S6.** Methylation patterns of MCCs across samples in each **(a)** purified peripheral blood cell type (*n* = 28 for each cell type) and **(b)** purified cord blood cell type (*n* = 11 for each cell type). Methylation variability was calculated as the standard deviation (SD) of CpG methylation (beta values) across samples in each cell type. Abbreviation: NK cell, natural killer cell. Mean values are presented as circles in each box plot. Outliers are not shown in box plots. Statistical significance is denoted separately for the comparisons of MCCs with SCCs and NCCs by asterisks (from top to bottom: MCC vs. SCC; MCC vs. NCC; SCC vs. NCC). * FDR < 0.05; ** FDR < 0.01; *** FDR < 0.001.

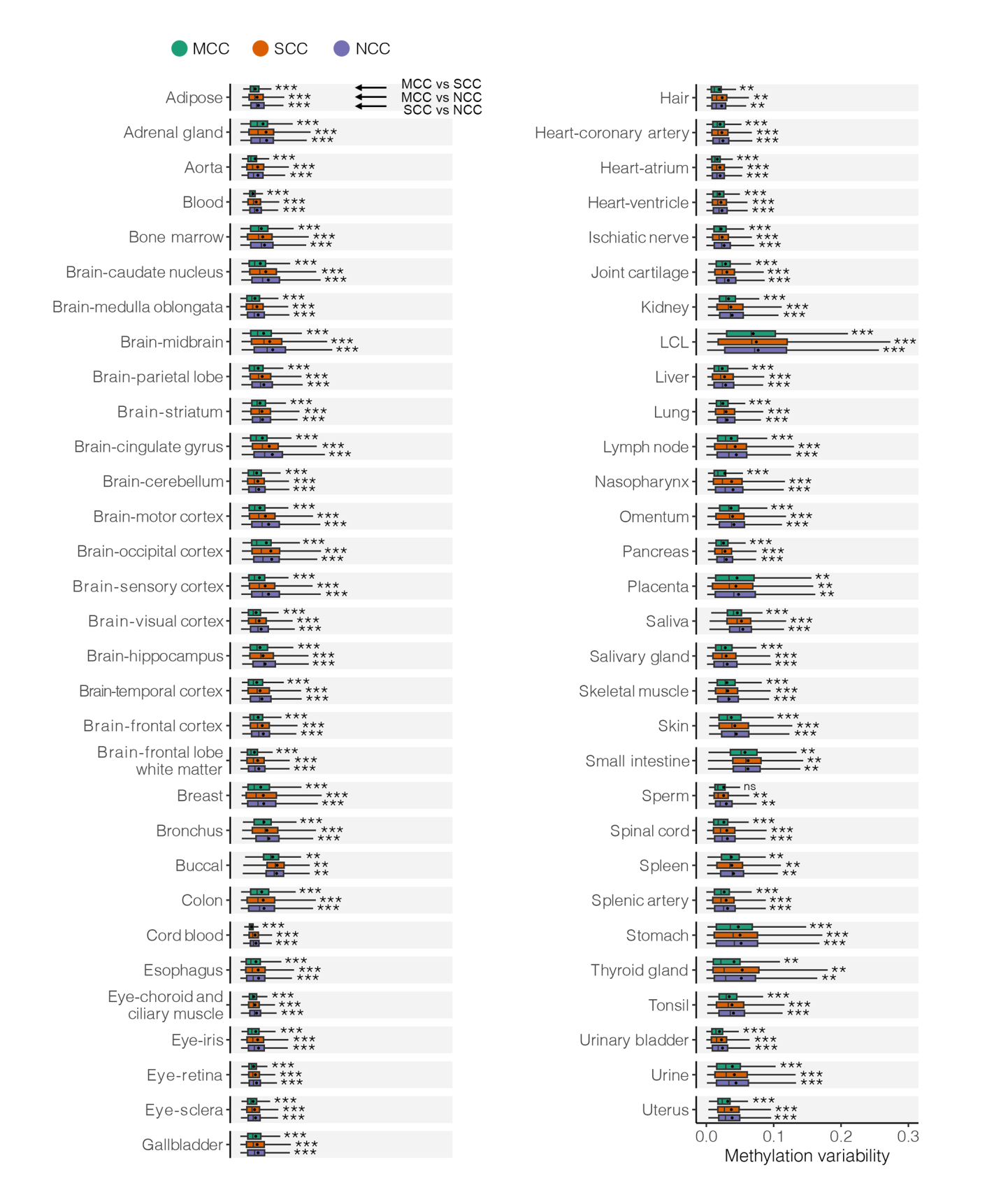

**Figure S7.** Box plots summarizing differential methylation variability (SD) between MCCs, SCCs, and NCCs in 42 tissues. Some tissues included samples from multiple regions. The average sample size was 165 (range 4–4097). Abbreviation: LCL, lymphoblastoid cell line. For some complex organs, distinct regions in the organ were used. Methylation variability was calculated as the SD of CpG methylation (beta values) across samples in each tissue. Mean values are presented as circles in each box plot. Outliers are not shown in box plots. Statistical significance is denoted separately for the comparisons of MCCs with SCCs and NCCs by asterisks (from top to bottom: MCC vs. SCC; MCC vs. NCC; SCC vs. NCC). * FDR < 0.05; ** FDR < 0.01; *** FDR < 0.001.

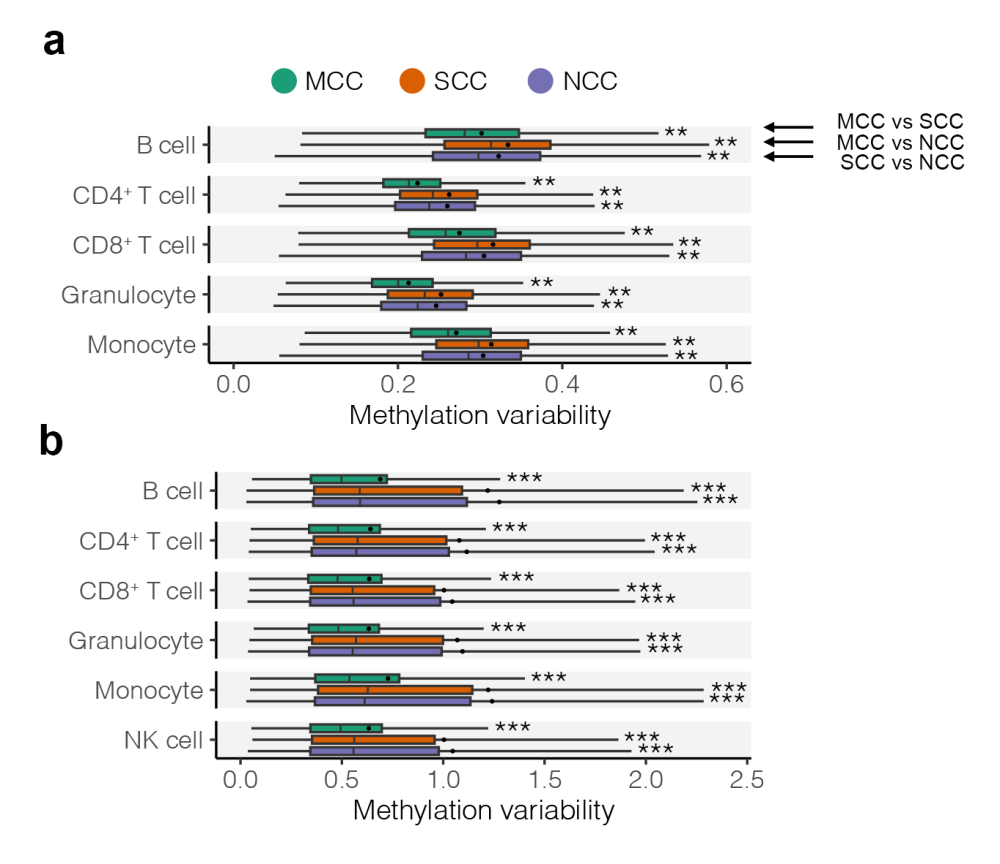

**Figure S8.** Validation of Methylation patterns of MCCs across samples in **(a)** purified peripheral blood cell types (*n* = 28 for each cell type) and **(b)** purified cord blood cell types (*n* = 11 for each cell type) using M values. Methylation variability was calculated as the SD of CpG methylation (M values) across samples in each tissue. Mean values are presented as circles in each box plot. Outliers are not shown in box plots. Statistical significance is denoted separately for the comparisons of MCCs with SCCs and NCCs by asterisks (from top to bottom: MCC vs. SCC; MCC vs. NCC; SCC vs. NCC). * FDR < 0.05; ** FDR < 0.01; *** FDR < 0.001.

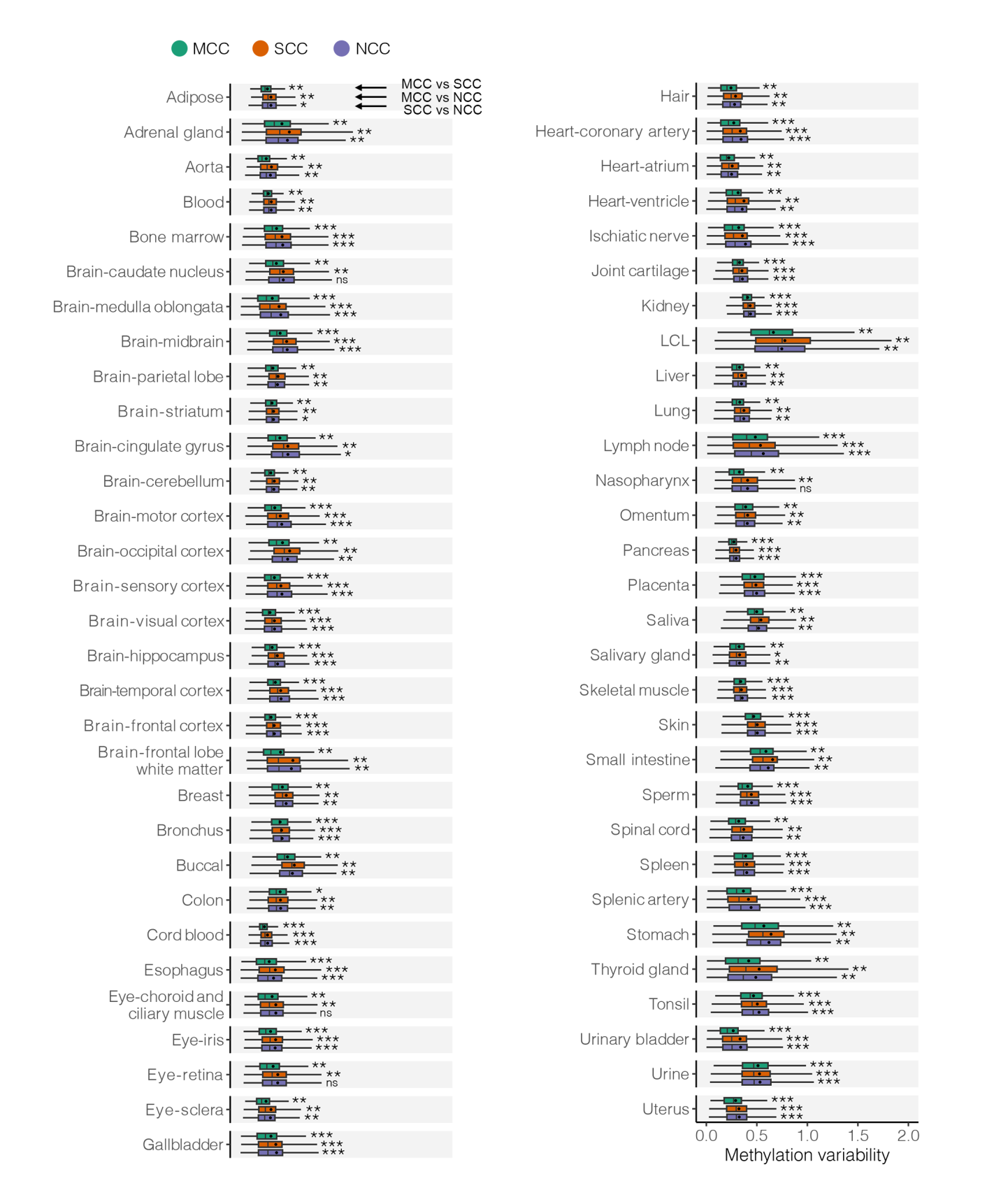

**Figure S9.** Validation of Methylation patterns of MCCs across samples in 42 tissues (*n* ranged from 127 to 4097) using M values. Abbreviation: LCL, lymphoblastoid cell line. Methylation variability was calculated as the SD of CpG methylation (M values) across samples in each tissue. Mean values are presented as circles in each box plot. Outliers are not shown in box plots. Statistical significance is denoted separately for the comparisons of MCCs with SCCs and NCCs by asterisks (from top to bottom: MCC vs. SCC; MCC vs. NCC; SCC vs. NCC). * FDR < 0.05; ** FDR < 0.01; *** FDR < 0.001.

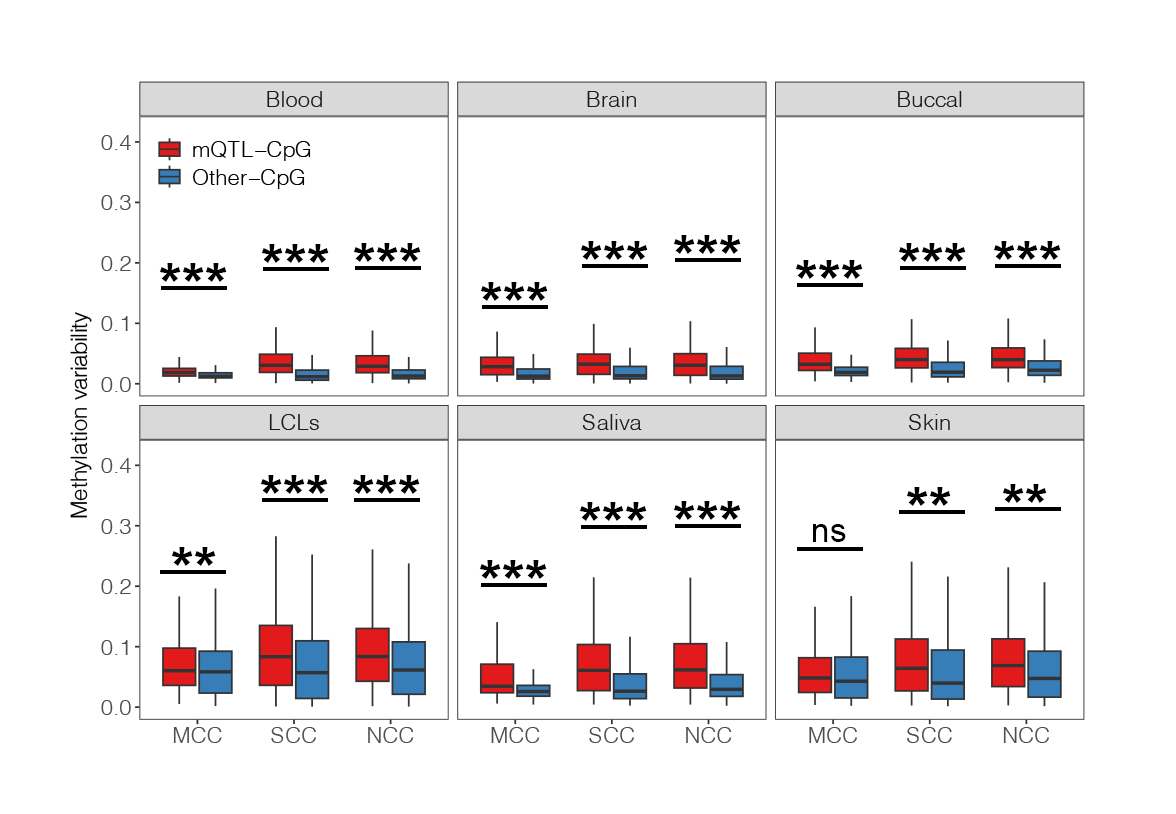

**Figure S10.** CpG-type subgroups of methylation variability differences between mQTL-CpGs and other CpGs (SNP-independent CpGs) in six common tissues. A total of 504 blood, 78 brain, 279 buccal swab, 89 lymphoblastoid cell line (LCL), 48 saliva, and 60 skin samples were used in our analyses. Abbreviation: LCL, lymphoblastoid cell line. Methylation variability was calculated as the SD of CpG methylation (beta values) across samples in each tissue. Outliers are not shown in box plots. * FDR < 0.05; ** FDR < 0.01; *** FDR < 0.001; ^ns^ nonsignificant.

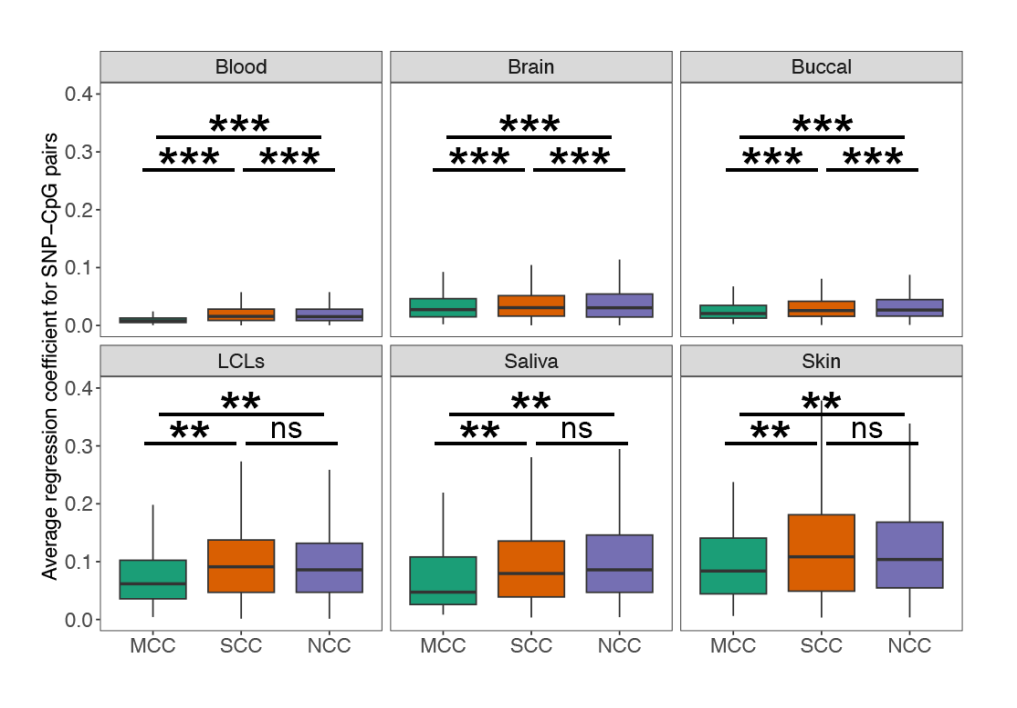

**Figure S11.** Comparison of regression coefficients for SNP-CpG pairs in MCCs, SCCs, and NCCs in six common tissues. A total of 504 blood, 78 brain, 279 buccal swab, 89 lymphoblastoid cell line (LCL), 48 saliva, and 60 skin samples were used in our analyses. Only tag-SNPs selected by PLINK were considered to reduce possible bias due to the distinct numbers of SNPs within haplotypes. Abbreviation: LCL, lymphoblastoid cell line. Outliers are not shown in box plots. * FDR < 0.05; ** FDR < 0.01; *** FDR < 0.001; ^ns^ nonsignificant.

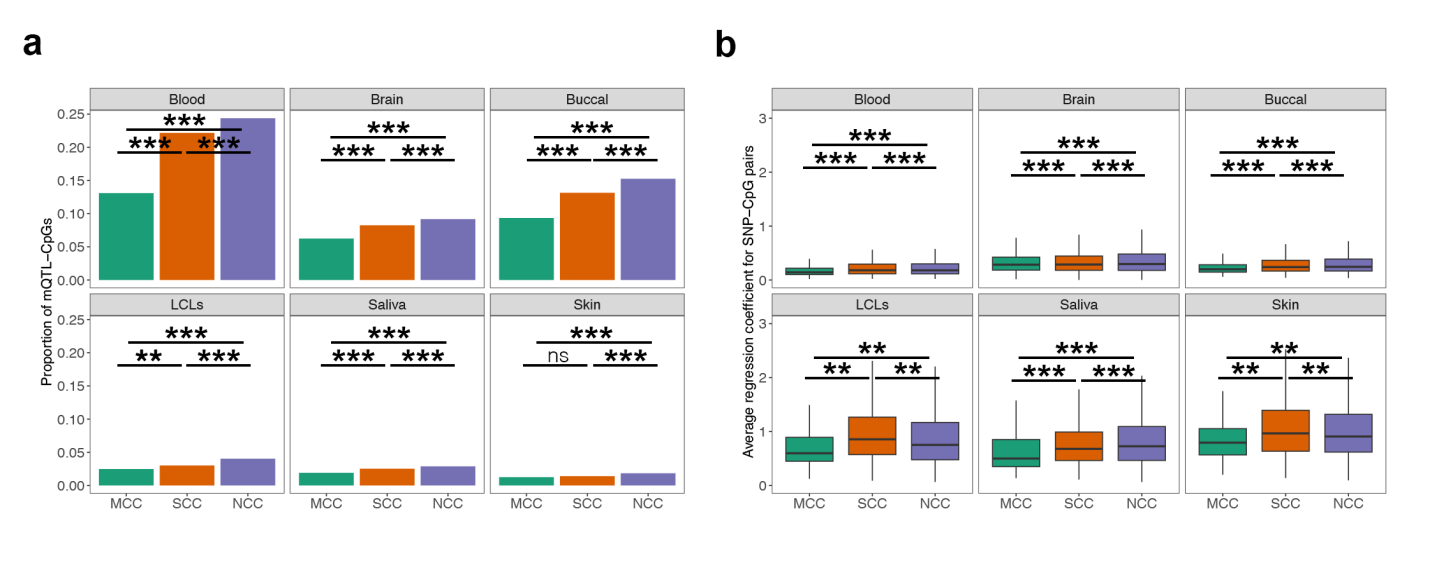

**Figure S12.** Validation of characteristics of the association between genetic variation and MCCs using M values. A total of 504 blood, 78 brain, 279 buccal swab, 89 lymphoblastoid cell line (LCL), 48 saliva, and 60 skin samples were used in our analyses. **(a)** Comparison of mQTL-CpG proportions in MCCs, SCCs, and NCCs in the six tissues. The two-proportions z-test was used to compare each pairwise combination of proportions. * FDR < 0.05; ** FDR < 0.01; *** FDR < 0.001; ^ns^ nonsignificant. **(b)** Comparison of regression coefficient for SNP-CpG pairs in MCCs, SCCs, and NCCs in the six tissues. The absolute values of regression coefficients were used. Outliers are not shown in box plots.

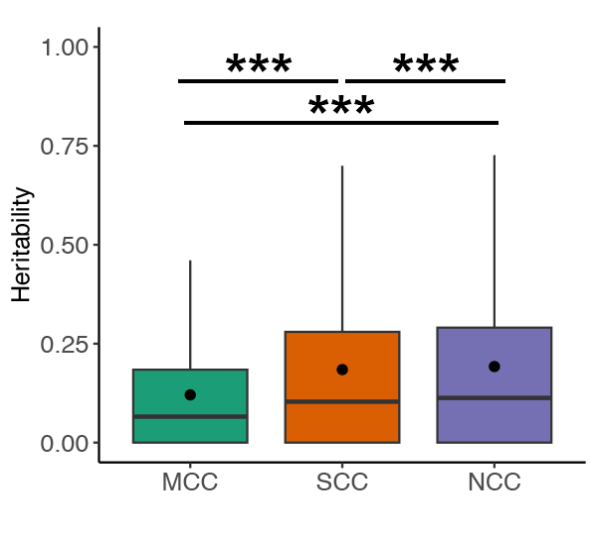

**Figure S13.** Box plot summarizing the genetic heritability differences between MCCs, SCCs, and NCCs in a twin cohort using M values. Heritability was estimated by Falconer’s formula [H^2^ = 2(*r*_MZ_ − *r*_DZ_)]. Mean values are presented as circles in each box plot. Outliers are not shown in box plots. *** FDR < 0.001.

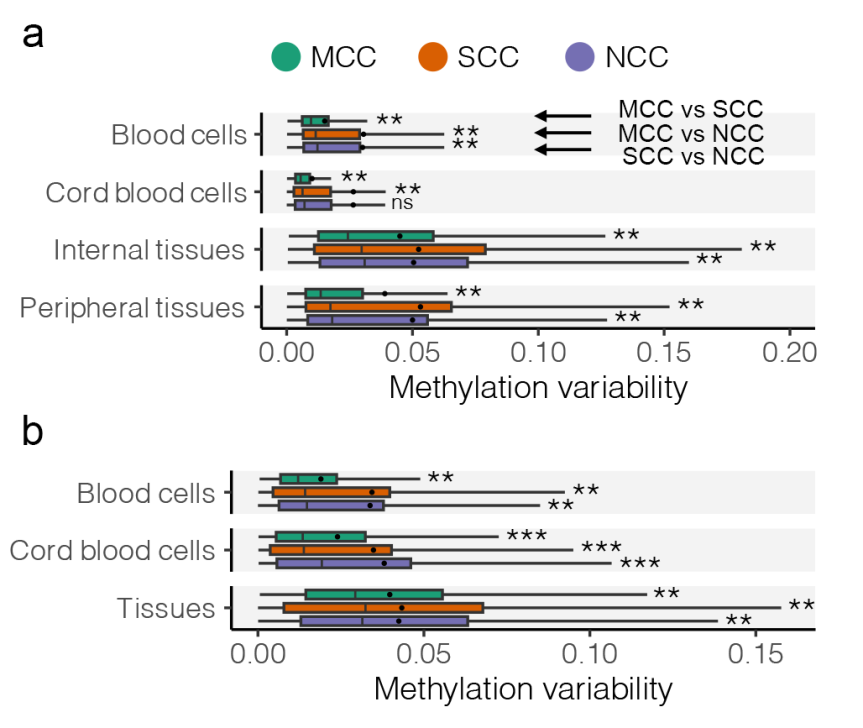

**Figure S14.** Investigation **(a)** and validation **(b)** of methylation variability of MCCs across tissues (*n* = 30 and 26 for internal and peripheral tissues) and cell types (*n* = 140 and 66 in blood and cord blood cells). Validation was conducted on independent data sets (*n* = 31, 48, and 66 for blood cells, cord blood cells, and tissues). Blood cells included B cells, CD4^+^ T cells, CD8^+^ T cells, granulocytes, and monocytes (validation data also included neutrophils). Cord blood cells included B cells, CD4^+^ T cells, CD8^+^ T cells, granulocytes, monocytes, and natural killer (NK) cells (validation data also included red blood cells). Internal tissues included liver, muscle, pancreas, subcutaneous fat, omentum, and spleen. Peripheral tissues included blood, saliva, buccal swab, and hair follicles. Tissues in validation data set included adipose abdominal, adipose subcutaneous, aorta abdominal, aorta thoracic, bladder, bone, bone marrow red, bone marrow yellow, coronary artery, gallbladder, gastric mucosa, ischiatic nerve, joint cartilage, lymph node, medulla oblongata, splenic artery, and tonsils. Only multiple tissue samples collected from the same individual were used in the analysis. Methylation variability was calculated as the SD of CpG methylation (beta values) across tissues or cell types. Mean values are presented as circles in each box plot. Outliers are not shown in box plots. Statistical significance is denoted separately for the comparisons of MCCs with SCCs and NCCs by asterisks (from top to bottom: MCC vs. SCC; MCC vs. NCC; SCC vs. NCC). * FDR < 0.05; ** FDR < 0.01; *** FDR < 0.001.

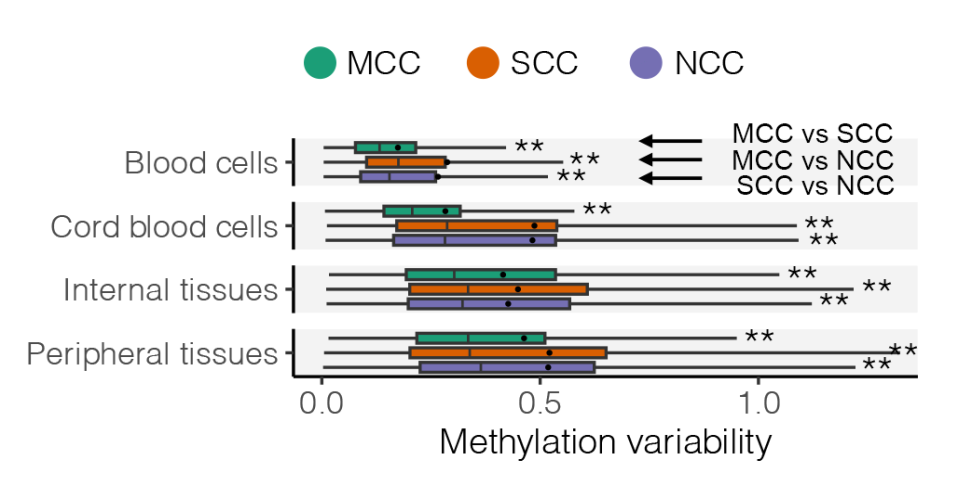

**Figure S15.** Validation of methylation variability of MCCs across tissues (*n* = 30 and 26 for internal and peripheral tissues) and cell types (*n* = 140 and 66 in blood and cord blood cells) using M values. Blood cells included B cells, CD4^+^ T cells, CD8^+^ T cells, granulocytes, and monocytes. Cord blood cells included B cells, CD4^+^ T cells, CD8^+^ T cells, granulocytes, monocytes, and natural killer (NK) cells. Internal tissues included liver, muscle, pancreas, subcutaneous fat, omentum, and spleen. Peripheral tissues included blood, saliva, buccal swab, and hair follicles. Only multiple tissue samples collected from the same individual were used in the analysis. Methylation variability was calculated as the SD of CpG methylation (M values) across tissues or cell types. Mean values are presented as circles in each box plot. Outliers are not shown in box plots. Statistical significance is denoted separately for the comparisons of MCCs with SCCs and NCCs by asterisks (from top to bottom: MCC vs. SCC; MCC vs. NCC; SCC vs. NCC). * FDR < 0.05; ** FDR < 0.01; *** FDR < 0.001.

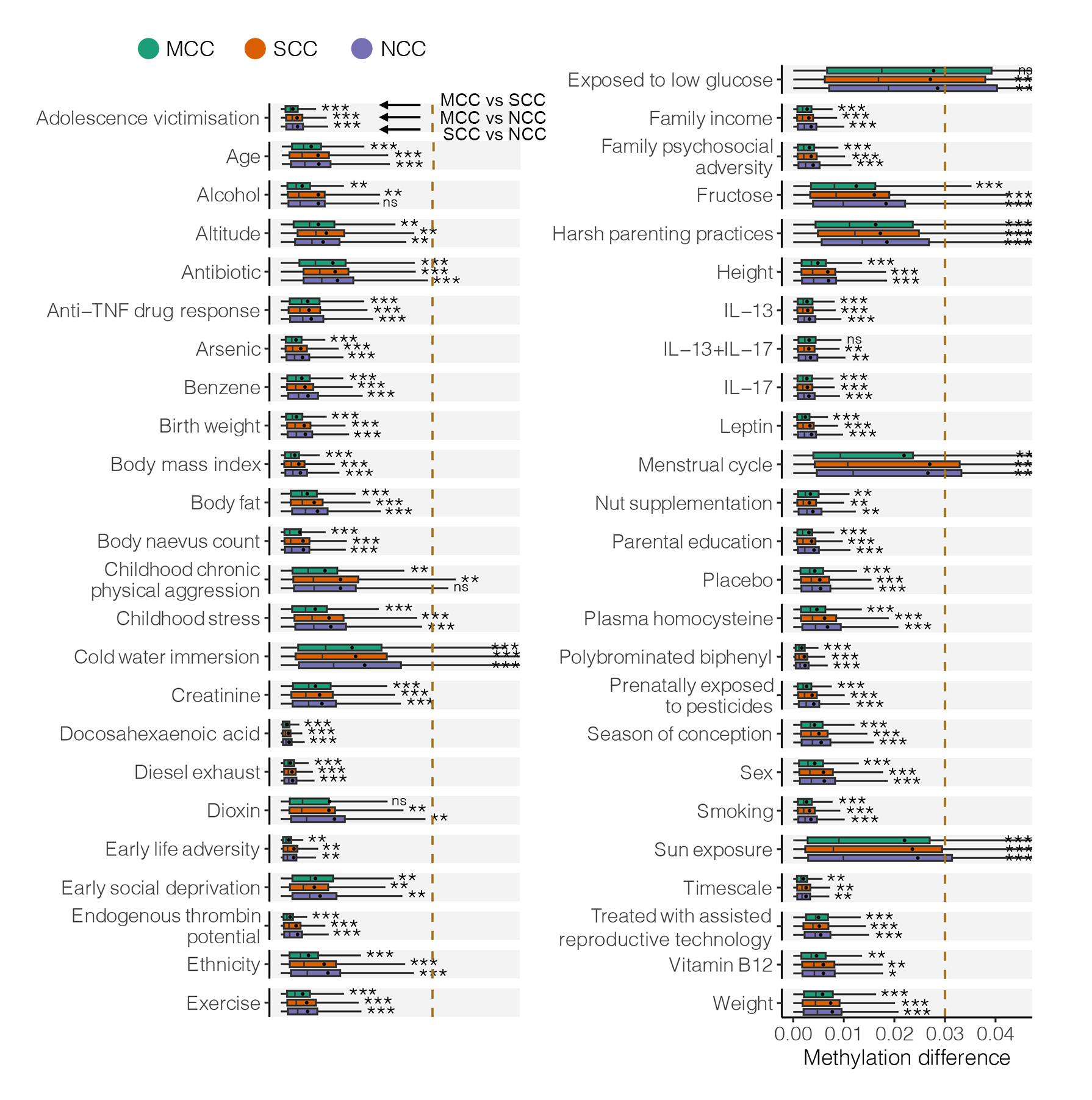

**Figure S16.** Methylation differences in MCCs between the groups of each demographic and environmental factor. Common factors, such as age, sex, ethnicity, and body mass index, were investigated in multiple tissues (results shown in Supplementary Table S4). The average sample size was 124 (range 8–440). Abbreviations: IL-13, interleukin-13; IL-17, interleukin-17; TNF, tumor necrosis factor. The dashed line represents the commonly used methylation change threshold of 0.03. Methylation difference was defined as the absolute CpG beta value change between groups. Mean values are presented as circles in each box plot. Outliers are not shown in box plots. Statistical significance is denoted separately for the comparisons of MCCs with SCCs and NCCs by asterisks (from top to bottom: MCC vs. SCC; MCC vs. NCC; SCC vs. NCC). * FDR < 0.05; ** FDR < 0.01; *** FDR < 0.001. To improve graphical display, the right limit of the x-axis is specified as the maximum value of the third quartile for methylation differences under demographic/environmental factors.

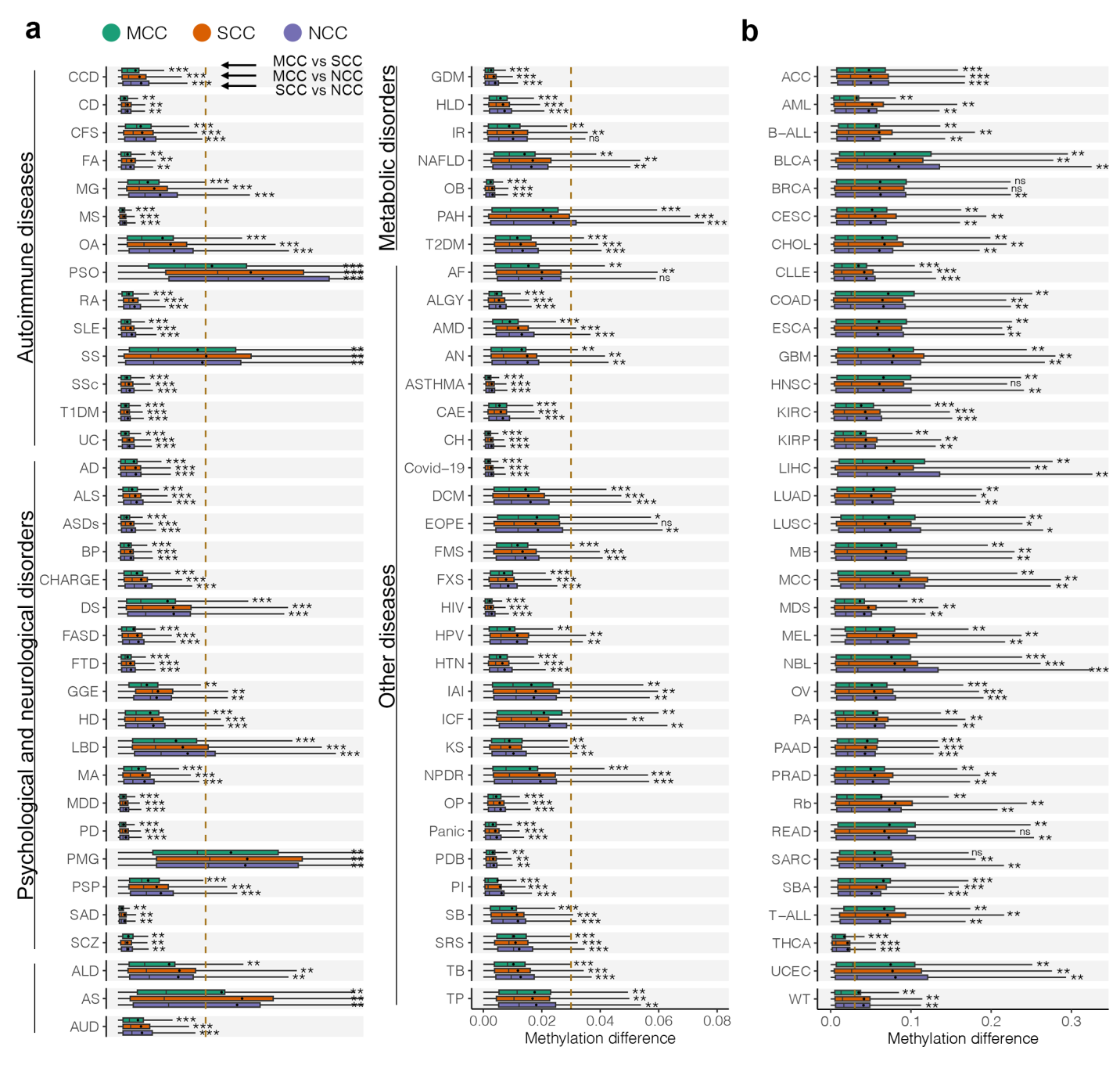

**Figure S17.** Methylation differences in MCCs between patient and healthy samples for each disease. **(a)** Box plots summarizing differential methylation differences between MCCs, SCCs, and NCCs in 69 noncancer diseases. The average sample size was 109 (range 10–689).

Abbreviations:

*Autoimmune diseases (n = 14):* CCD, celiac disease; CD, Crohn’s disease; CFS, chronic fatigue syndrome; FA, food allergy; MG, myasthenia gravis; MS, multiple sclerosis; OA, osteoarthritis; PSO, psoriasis; RA, rheumatoid arthritis; SLE, systemic lupus erythematosus; SS, Sjögren’s syndrome; SSc, systemic sclerosis; T1DM, type 1 diabetes mellitus; UC, ulcerative colitis.

*Psychological and neurological diseases (n = 18):* AD, Alzheimer’s disease; ALS, amyotrophic lateral sclerosis; ASDs, autism spectrum disorder; BP, bipolar disorder; CHARGE, CHARGE syndrome; DS, Down syndrome; FASD, fetal alcohol spectrum disorder; FTD, frontotemporal dementias; GGE, genetic generalized epilepsy; HD, Huntington’s disease; LBD, Lewy body dementia; MA, congenital microcephaly; MDD, major depressive disorder; PD, Parkinson’s disease; PMG, polymicrogyria; PSP, progressive supranuclear palsy; SAD, social anxiety disorder; SCZ, schizophrenia.

*Metabolic disorders (n = 10):* ALD, adrenoleukodystrophy; AS, atherosclerotic lesions; AUD, alcohol use disorders; GDM, gestational diabetes mellitus; HLD, hyperlipidemia; IR, insulin resistance; NAFLD, nonalcoholic fatty liver disease; OB, obesity; PAH, pulmonary arterial hypertension; T2DM, type 2 diabetes mellitus.

*Other diseases (n = 27):* AF, atrial fibrillation; ALGY, allergic diseases; AMD, age-related macular degeneration; AN, anencephalic fetus; ASTHMA, asthma; CAE, coronary artery ectasia; CH, congenital hypopituitarism; COVID-19, coronavirus disease 2019; DCM, dilated cardiomyopathy; EOPE, early onset preeclampsia; FMS, fibromyalgia; FXS, fragile X syndrome; HIV, human immunodeficiency virus infection; HPV, human papillomavirus infection; HTN, hypertension; IAI, intraamniotic infection; ICF, immunodeficiency, centromere instability, and facial anomalies syndrome; KS, Kabuki syndrome; NPDR, nonproliferative diabetic retinopathy; OP, osteoporosis; Panic, panic disorder; PDB, Paget’s disease of bone; PI, preterm infant; SB, spina bifida; SRS, Silver-Russell syndrome; TB, active pulmonary tuberculosis; TP, trisomy pregnancy.

The dashed line represents the commonly used methylation change threshold of 0.03. Methylation difference was defined as the absolute CpG beta value change between groups. Mean values are presented as circles in each box plot. Outliers are not shown in box plots. Statistical significance is denoted separately for the comparisons of MCCs with SCCs and NCCs by asterisks (from top to bottom: MCC vs. SCC; MCC vs. NCC; SCC vs. NCC). * FDR < 0.05; ** FDR < 0.01; *** FDR < 0.001. **(b)** Box plots summarizing differential methylation differences between MCCs, SCCs, and NCCs in 34 types of cancer. The average sample size was 255 (range 8–867).

Abbreviations: ACC, adrenocortical carcinoma; AML, acute myeloid leukemia; B-ALL, B-cell acute lymphoblastic leukemia; BLCA, bladder urothelial carcinoma; BRCA, breast invasive carcinoma; CESC, cervical squamous cell carcinoma and endocervical adenocarcinoma; CHOL, cholangiocarcinoma; CLLE, chronic lymphocytic leukemia; COAD, colon adenocarcinoma; ESCA, esophageal carcinoma; GBM, glioblastoma; HNSC, head and neck squamous cell carcinoma; KIRC, kidney renal clear cell carcinoma; KIRP, kidney renal papillary cell carcinoma; LIHC, liver hepatocellular carcinoma; LUAD, lung adenocarcinoma; LUSC, lung squamous cell carcinoma; MB, medulloblastoma; MCC, Merkel cell carcinoma; MDS, myelodysplastic syndrome; MEL, uveal melanoma; NBL, neuroblastoma; OV, ovarian cancer; PA, juvenile pilocytic astrocytoma tumor; PAAD, pancreatic adenocarcinoma; PRAD, prostate adenocarcinoma; Rb, retinoblastoma; READ, rectum adenocarcinoma; SBA, small bowel adenocarcinoma; SARC, sarcoma; T-ALL, T-cell acute lymphoblastic leukemia; THCA, thyroid carcinoma; UCEC, uterine corpus endometrial carcinoma; WT, Wilms’ tumor.

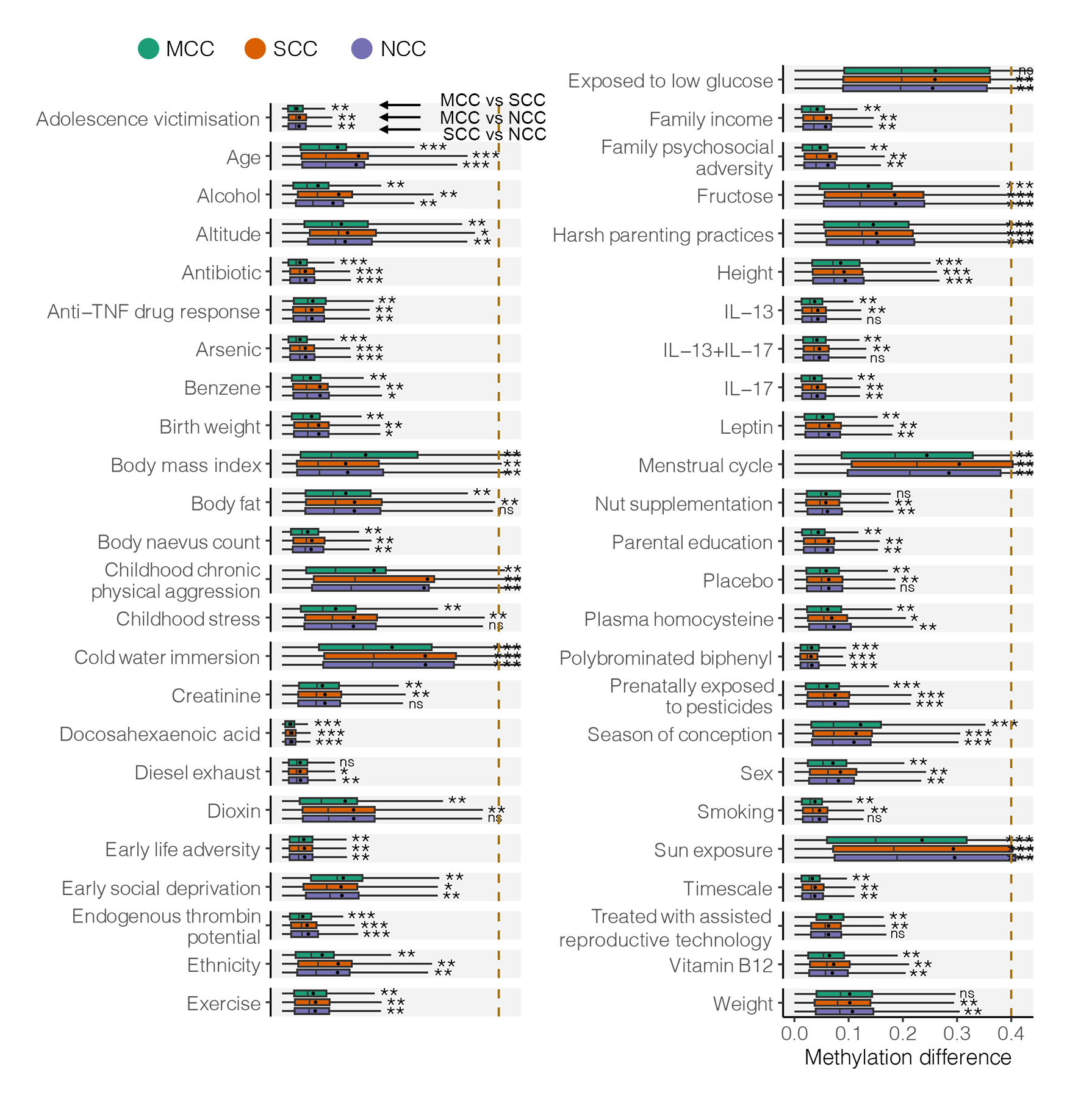

**Figure S18.** Validation of methylation differences in MCCs between the groups of each demographic and environmental factors using M values. The average sample size was 124 (range 8–440). The dashed line represents the suggested M change threshold of 0.4. Methylation difference was defined as the absolute CpG M value change between groups. Mean values are presented as circles in each box plot. Outliers are not shown in box plots. Statistical significance is denoted separately for the comparisons of MCCs with SCCs and NCCs by asterisks (from top to bottom: MCC vs. SCC; MCC vs. NCC; SCC vs. NCC). * FDR < 0.05; ** FDR < 0.01; *** FDR < 0.001.

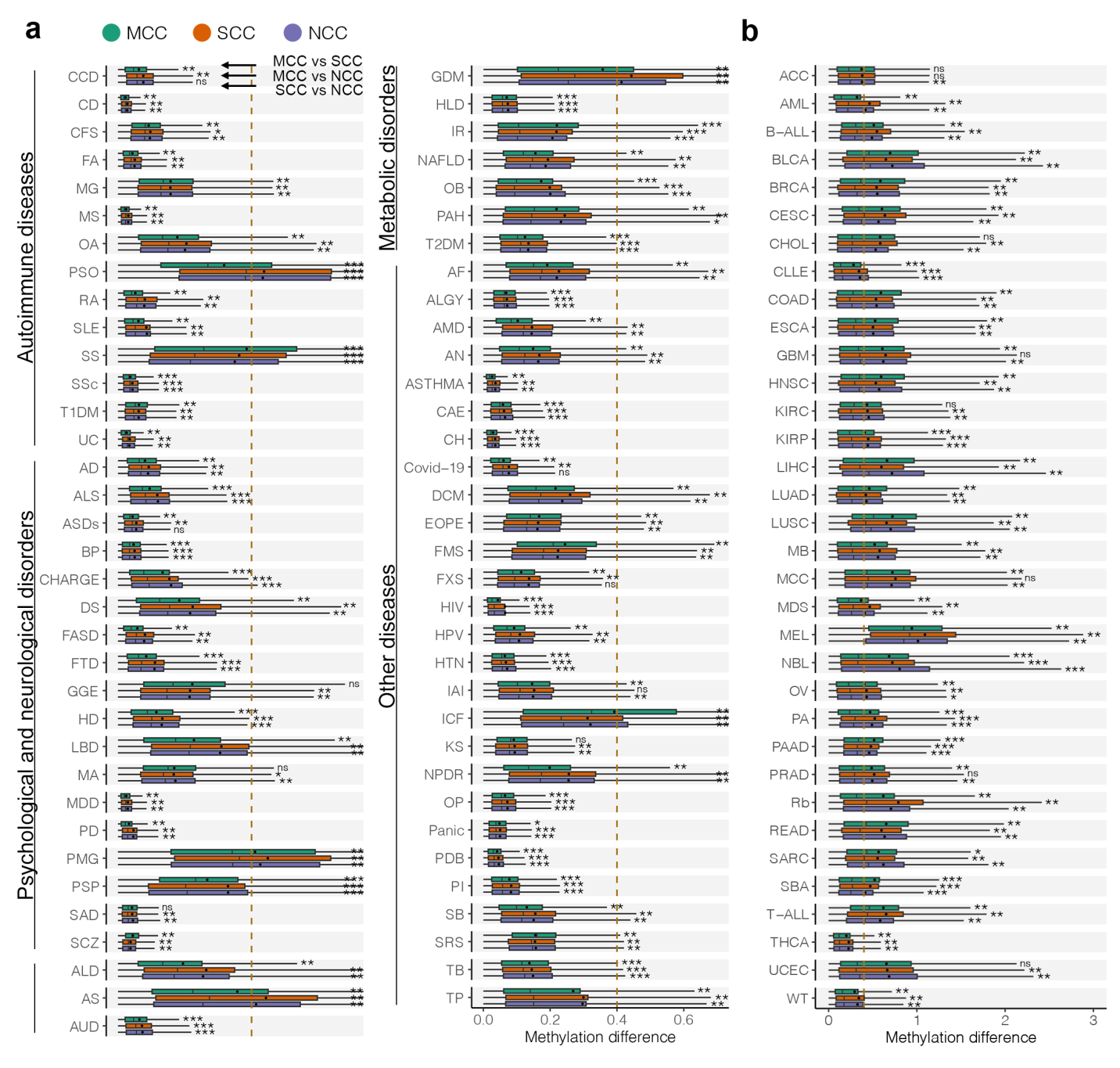

**Figure S19.** Validation of methylation differences in MCCs between the groups of non-cancer diseases, and cancers using M values. The dashed line represents the suggested M change threshold of 0.4. Methylation difference was defined as the absolute CpG M value change between groups. Mean values are presented as circles in each box plot. Outliers are not shown in box plots. Statistical significance is denoted separately for the comparisons of MCCs with SCCs and NCCs by asterisks (from top to bottom: MCC vs. SCC; MCC vs. NCC; SCC vs. NCC). * FDR < 0.05; ** FDR < 0.01; *** FDR < 0.001.

**(a)** Box plots summarizing differential methylation differences between MCCs, SCCs, and NCCs in each noncancer disease. The average sample size was 416 (range 212–689).

Abbreviations: FASD, fetal alcohol spectrum disorder; FTD, frontotemporal dementias; HIV, human immunodeficiency virus infection; MS, multiple sclerosis; OB, obesity; PD, Parkinson’s disease; PDB, Paget’s disease of bone; PSP, progressive supranuclear palsy; RA, rheumatoid arthritis; SCZ, schizophrenia.

**(b)** Box plots summarizing differential methylation differences between MCCs, SCCs, and NCCs in each type of cancer. The average sample size was 553 (range 419–867).

Abbreviations: B-ALL, B-cell acute lymphoblastic leukemia; BLCA, bladder urothelial carcinoma; BRCA, breast invasive carcinoma; HNSC, head and neck squamous cell carcinoma; KIRC, kidney renal clear cell carcinoma; LIHC, liver hepatocellular carcinoma; LUAD, lung adenocarcinoma; PRAD, prostate adenocarcinoma; THCA, thyroid carcinoma; UCEC, uterine corpus endometrial carcinoma.

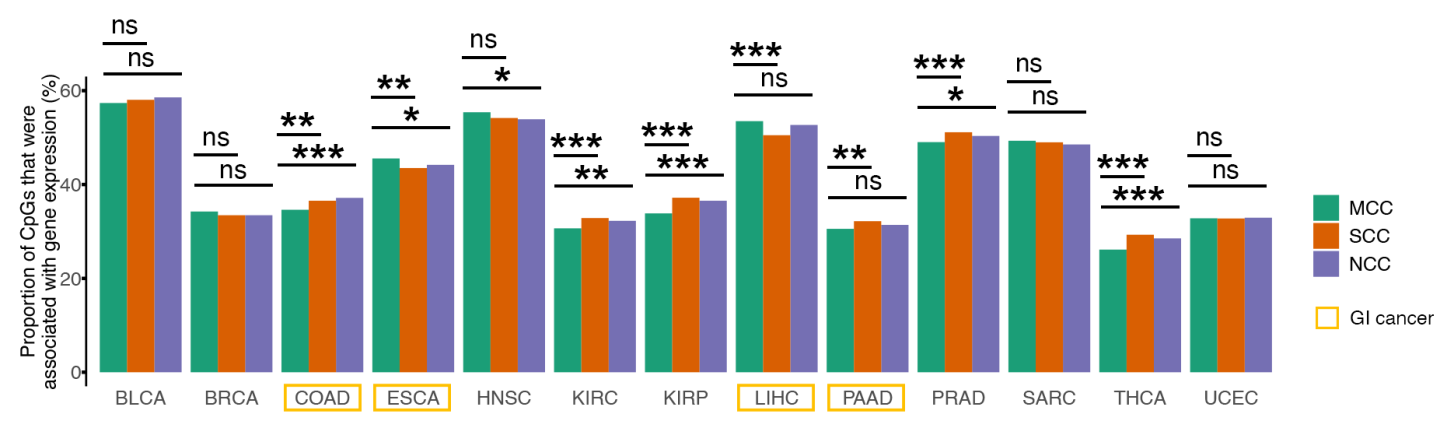

**Figure S20.** Comparison of proportions in MCCs, SCCs, and NCCs associated with gene expression in 13 cancers. Abbreviations: BLCA, bladder urothelial carcinoma; BRCA, breast invasive carcinoma; COAD, colon adenocarcinoma; ESCA, esophageal carcinoma; GI, gastrointestinal; HNSC, head and neck squamous cell carcinoma; KIRC, kidney renal clear cell carcinoma; KIRP, kidney renal papillary cell carcinoma; LIHC, liver hepatocellular carcinoma; PAAD, pancreatic adenocarcinoma; PRAD, prostate adenocarcinoma; SARC, sarcoma; THCA, thyroid carcinoma; UCEC, uterine corpus endometrial carcinoma. The two-proportions z-test was used to compare each pairwise combination of proportions. These cancers included four gastrointestinal cancers (COAD, ESCA, LIHC, and PAAD) and nine non-gastrointestinal cancers (BLCA, BRCA, HNSC, KIRC, KIRP, PRAD, SARC, THCA, and UCEC). * FDR < 0.05; ** FDR < 0.01; *** FDR < 0.001; ^ns^ nonsignificant.

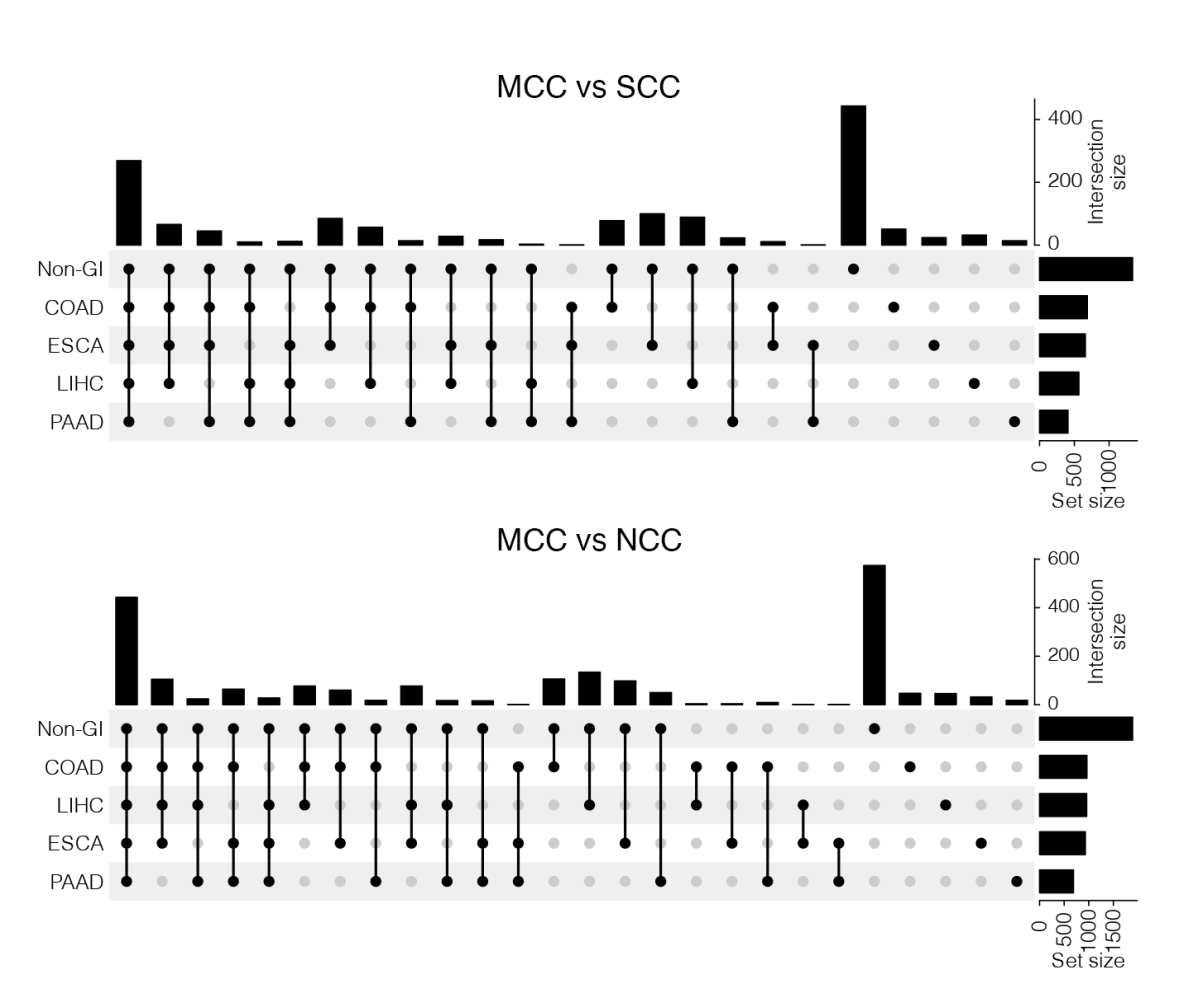

**Figure S21.** Upset plots of biological pathways across genes whose expression is associated with MCC methylation in various cancers. Non-gastrointestinal cancer (non-GI) is a combination of pathways from nine non-gastrointestinal cancers: bladder urothelial carcinoma (BLCA), breast invasive carcinoma (BRCA), head and neck squamous cell carcinoma (HNSC), kidney renal clear cell carcinoma (KIRC), kidney renal papillary cell carcinoma (KIRP), prostate adenocarcinoma (PRAD), sarcoma (SARC), thyroid carcinoma (THCA), and uterine corpus endometrial carcinoma (UCEC). The four gastrointestinal cancers were colon adenocarcinoma (COAD), esophageal carcinoma (ESCA), liver hepatocellular carcinoma (LIHC), and pancreatic adenocarcinoma (PAAD).

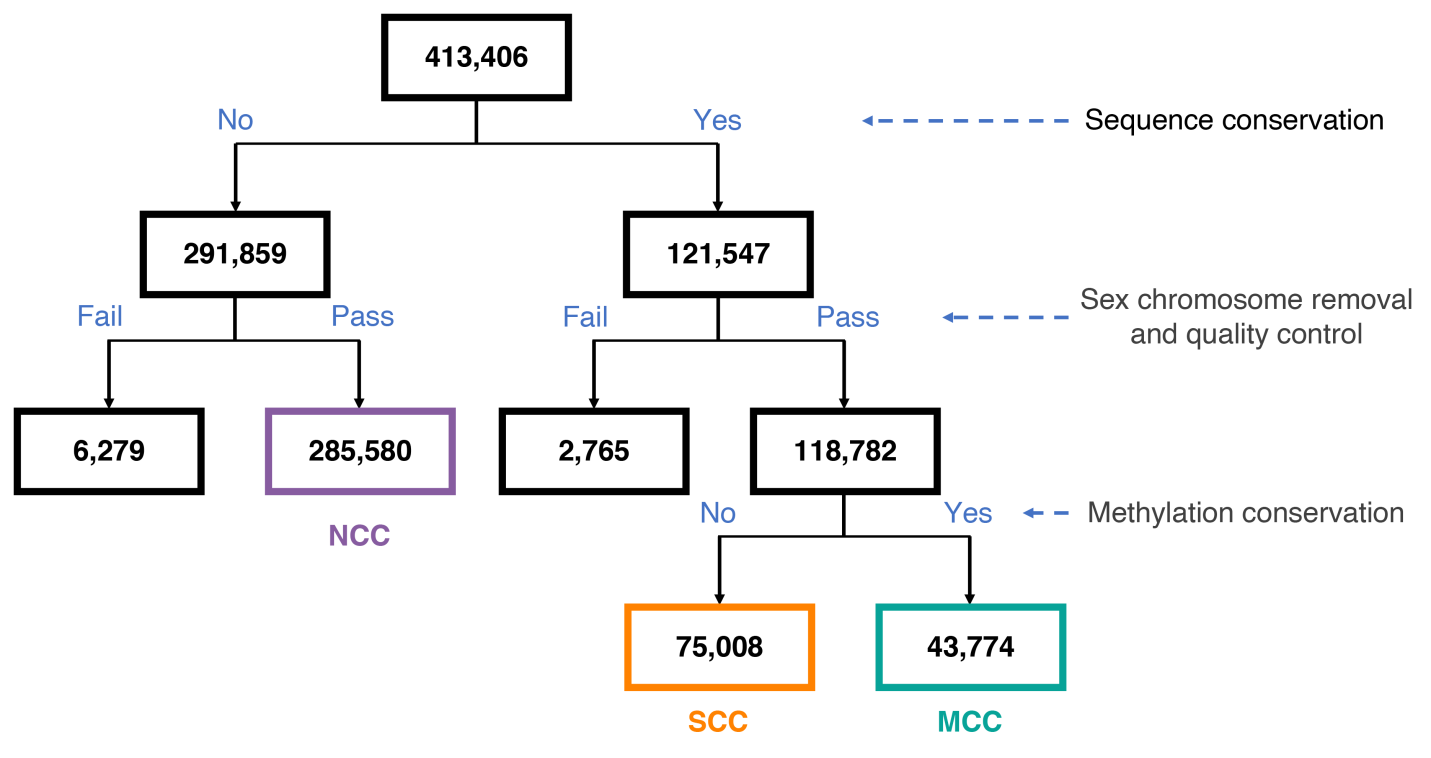

**Figure S22.** Flowchart of MCC discovery in additional CpGs on the EPIC array.

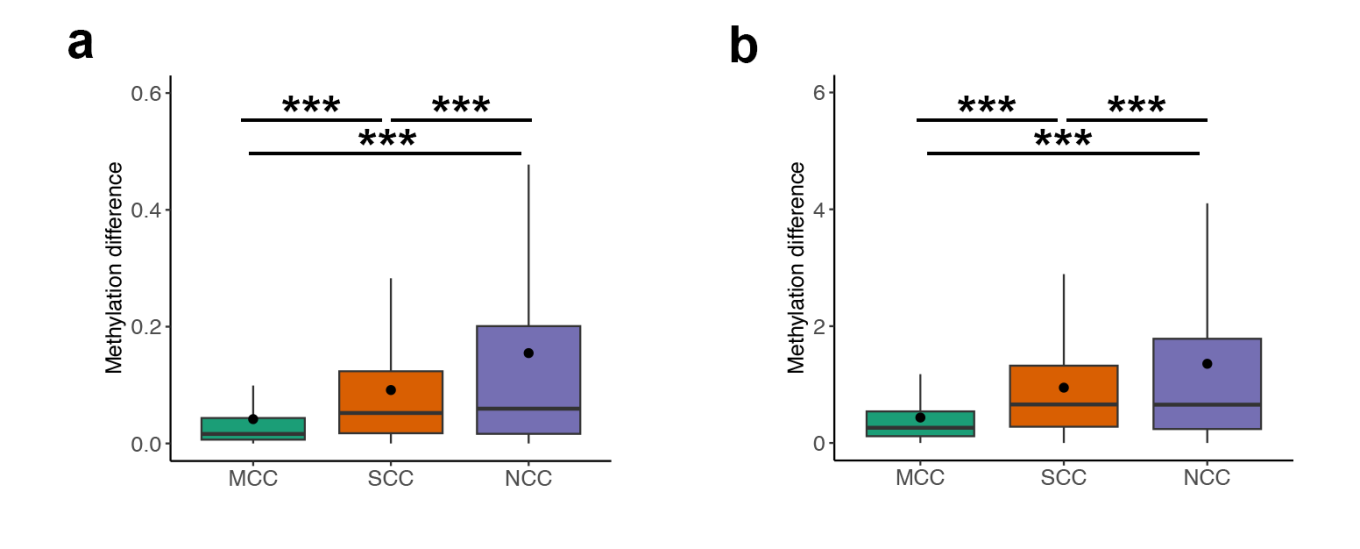

**Figure S23.** Validation of MCCs identified on the EPIC array in the cerebellum between chimpanzees (*n* = 8) and humans (*n* = 7) using **(a)** beta values and **(b)** beta-converted M values. Mean values are presented as circles in each box plot. Outliers are not shown in box plots. * FDR < 0.05; ** FDR < 0.01; *** FDR < 0.001; ^ns^ nonsignificant.

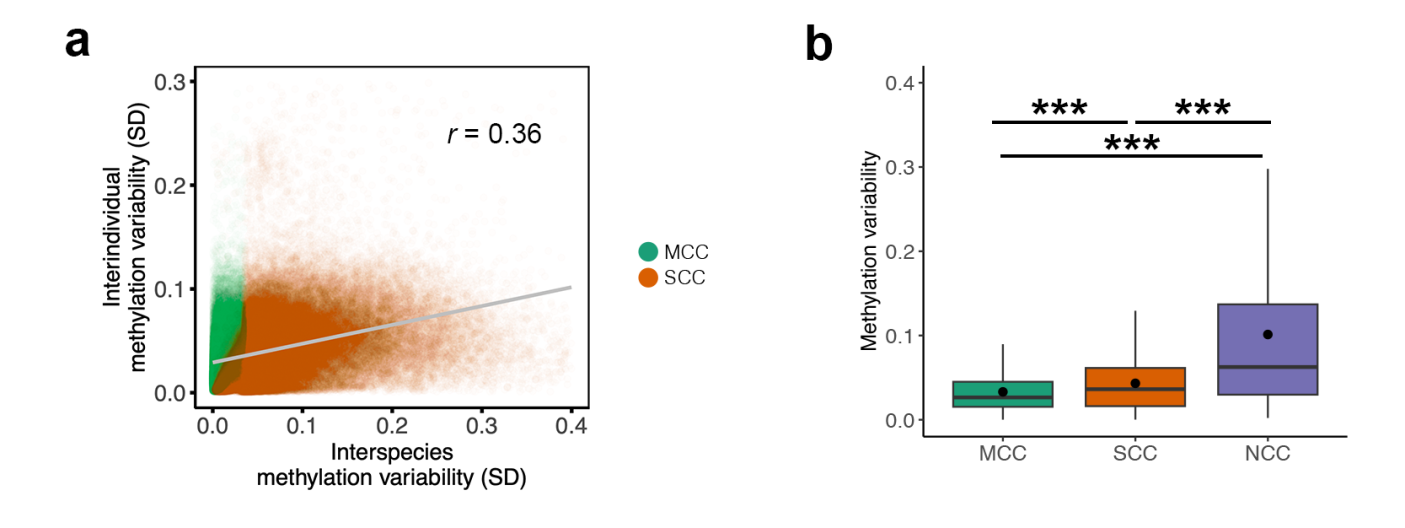

**Figure S24.** Positive association between methylation conservation and methylation stability in additional CpGs from the EPIC array. **(a)** Smooth scatterplots of interindividual and interspecies methylation variability within additional EPIC probes (*n* = 121,547) matched to all great ape genomes. Methylation variability was calculated as the SD of CpG methylation across samples (*n* = 7). **(b)** Box plots summarizing differential methylation variability between MCCs and SCCs in seven human prefrontal cortex samples. Methylation variability was calculated separately as the SD of CpG methylation (beta value) across samples. Mean values are presented as circles in each box plot. Outliers are not shown in box plots. * FDR < 0.05; ** FDR < 0.01; *** FDR < 0.001.

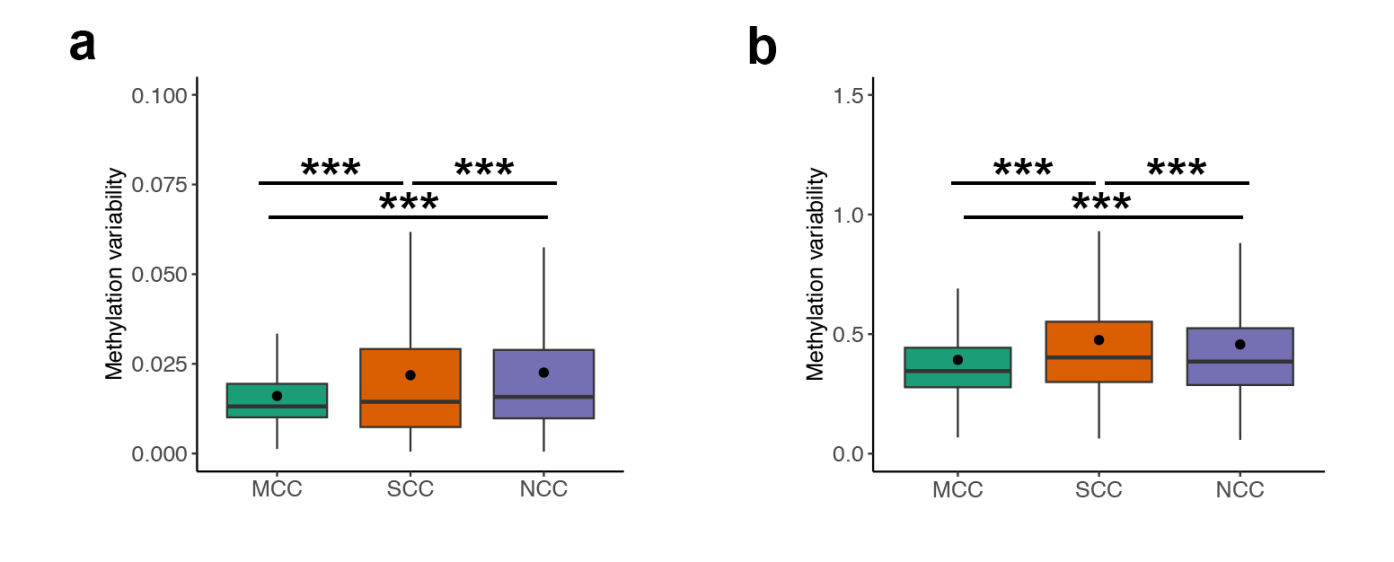

**Figure S25.** Box plots summarizing differential methylation variability (SD) between MCCs, SCCs, and NCCs in human blood samples (*n* = 504). Methylation variability was calculated as the SD of **(a)** beta values and **(b)** M values across samples in each tissue. Mean values are presented as circles in each box plot. Outliers are not shown in box plots. * FDR < 0.05; ** FDR < 0.01; *** FDR < 0.001.
